# A distinct core regulatory module enforces oncogene expression in KMT2A-rearranged leukemia

**DOI:** 10.1101/2021.08.03.454902

**Authors:** Taku Harada, Yaser Heshmati, Jérémie Kalfon, Juliana Xavier Ferrucio, Monika Perez, Jazmin Ewers, Andrew Kossenkov, Jana M. Ellegast, Joanna S. Yi, Allyson Bowker, Qian Zhu, Kenneth Eagle, Joshua M. Dempster, Guillaume Kugener, Jayamanna Wickramasinghe, Zachary T. Herbert, Charles H. Li, Jošt Vrabič Koren, David M. Weinstock, Vikram R. Paralkar, Behnam Nabet, Charles Y. Lin, Neekesh V. Dharia, Kimberly Stegmaier, Stuart H. Orkin, Maxim Pimkin

## Abstract

A small set of lineage-restricted transcription factors (TFs), termed core regulatory circuitry (CRC), control cell identity and malignant transformation. Here, we integrated gene dependency, chromatin architecture and TF perturbation datasets to characterize 31 core TFs in acute myeloid leukemia (AML). Contrary to a widely accepted model, we detected a modular CRC structure with hierarchically organized, partially redundant and only sparsely interconnected modules of core TFs controlling distinct genetic programs. Rapid TF degradation followed by measurement of genome-wide transcription rates revealed that core TFs directly regulate dramatically fewer genes than previously assumed. Leukemias carrying KMT2A (MLL) rearrangements depend on the IRF8/MEF2 axis to directly enforce expression of the key oncogenes MYC, HOXA9 and BCL2. Our datasets provide an evolving model of CRC organization in human cells, and a resource for further inquiries into and therapeutic targeting of aberrant transcriptional circuits in cancer.

## Introduction

Metazoan development depends on cell type-specific gene expression programs established and reinforced by combinatorial actions of spatiotemporally restricted transcription factors (TFs) (*1–4*). Although an average mammalian cell expresses several hundred TFs, only a small number, variably referred to as reprogramming, master or core regulatory (CR) TFs, are principally important for lineage specification (*5–9*). A generally accepted model posits that CR TFs positively regulate their own and each other’s genes, forming a network of interconnected feed-forward loops termed core regulatory circuitry (CRC) (*9–12*). CR TFs cooperatively enforce expression of key lineage genes by establishing extended, closely spaced enhancers with markedly high levels of histone acetylation and cofactor recruitment, termed super-enhancers (SEs) (*13–17*). SE patterns are specific markers of cell identity and have been exploited to infer developmental and oncogenic programs, as well as therapeutic vulnerabilities (*18–24*).

Progression of a normal cell to a neoplastic state is associated with transcriptional derangement (*11, 25, 26*). Mutations in TFs and chromatin regulators are common drivers of cancer (*27, 28*). In addition, cancer cells may be critically dependent on non-mutated TFs enforcing cancer-specific transcriptional programs in a process referred to as transcriptional addiction (*19, 25, 29–32*). The transcriptional circuitry of a malignant cell thus includes elements retained from its cell of origin as well as cancer-specific circuits established *de novo* or subverted from the transcriptional programs of other developmental stages or lineages (*33*). These context-specific circuitries cannot be predicted from the initiating tumorigenic mutation itself and have typically been identified through hypothesis-driven mechanistic exploration or comparative genome-wide screens. Meanwhile, to the extent that these *de novo* dependencies are cancer-restricted, they offer new opportunities for targeted therapy (*25, 26*).

Acute myeloid leukemia (AML) is the most common type of leukemia in adults and accounts for ∼20% of hematologic malignancy in children (*34, 35*). AML relies on deregulated expression of lineage TFs (*36, 37*) and is commonly caused by mutations in TFs and chromatin regulators (*27, 38, 39*). However, specific driver mutations do not correlate with enhancer architecture (*24, 33*), pointing to the existence of independently controlled, context-specific CRCs.

Here, we employ an integrative functional genomics analysis of the AML core transcriptional circuitry to systematically identify selective transcriptional vulnerabilities. Our study reveals four principles of AML CRC organization: hierarchy, modularity, divergence and redundancy. We demonstrate that divergent CRCs establish distinct subtypes of AML and underlie context-specific transcriptional addiction. We show that CRCs are stabilized by extensive redundancy and cross-compensation between co-expressed TF paralogs. We identify 4 TFs (RUNX2, IRF8, MEF2D and MEF2C) as selective transcriptional dependencies of leukemias carrying a KMT2A translocation and demonstrate that IRF8 and MEF2 proteins form a cancer-specific core regulatory module. We establish key leukemia oncogenes as direct transcriptional targets of the IRF8/MEF2 module, uncovering the mechanistic basis of leukemia addiction to these TFs. Our study significantly advances our understanding of transcriptional circuitries in cancer and provides an extensive data resource which will aid in the development of novel AML therapies.

## Results

### Recurrent SEs define distinct AML clusters

We assembled a cohort of AML cell lines (*n*=24), patient-derived xenograft (PDX) models (*40*) (*n*=38) and pediatric primary AMLs (*n*=19). We incorporated a previously published adult primary AML dataset (*24*) (*n*=49), for a total of 130 samples. We performed chromatin immunoprecipitation sequencing (ChIP-seq) for the enhancer histone mark H3K27ac and identified consensus SEs (*14*) (Figure 1A). This approach resulted in a total of 6868 distinct SEs, of which 4798 were recurrent in at least 2 samples and 2107 were shared between all sample types (Figures 1B, S1). To establish enhancer-gene connections on a genome scale we resolved the H3K27ac-mediated 2D genome architecture in a model AML cell line by HiChIP and calculated activity-by-contact scores (*41, 42*). We then integrated activity-by-contact with gene proximity and expression to assign SEs to target genes (Figure 1A). As a case study, we validated the predicted SE-gene connections in the *MYB* and *IRF8* loci by directly inhibiting enhancer activity (*43*) and confirming reduced expression of MYB and IRF8 (Figure S2).

**Figure 1.**
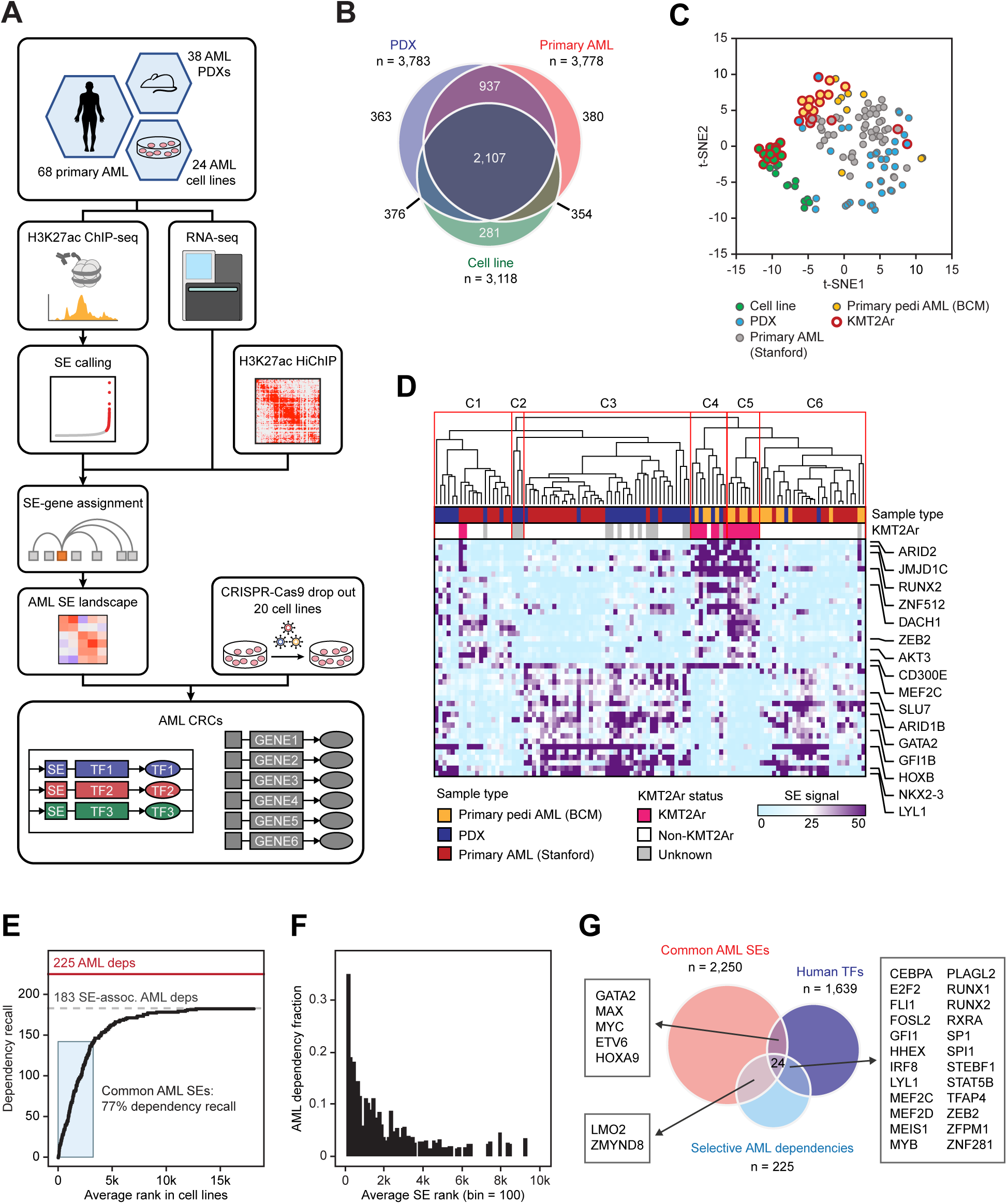
Integrative analyses of SE landscapes and enriched gene dependencies in AML. (A) Study schematic. (B) Overlap of distinct SEs in various sample types (C) Samples of all types are plotted according to the SE scores (4798 SEs recurrent in at least 2 samples) using t-distributed stochastic neighbor embedding (t-SNE). (D) Primary and PDX samples are hierarchically clustered using Pearson correlation of the scores of 4798 SEs recurrent in at least 2 samples. SEs with the largest average score difference between KMT2Ar and non-KMT2Ar leukemias are selectively shown. (E) A recall curve of association between selective AML dependencies and SEs. SE-associated genes are arranged on the horizontal axis in the order of decreasing average SE rank. The recall of selective AML dependencies is incrementally plotted on the vertical axis. (F) Top AML SEs are enriched for selective AML dependencies. SEs are binned according to their average rank (bin size = 100) and the fraction of SEs in each bin associated with genes representing selective AML dependencies is plotted on the vertical axis. (G) Overlap between common SEs (2250 SEs corresponding to 77% dependency recall), TFs and selective AML dependencies to define AML CRC. Refer to Figure S8 for details.

Cell lines differ from primary tumors in several key biological characteristics (*44–46*). PDX models, while capturing an underlying tumor’s biology more faithfully, undergo mouse-specific evolution (*47, 48*). Clustering algorithms placed cell lines into a separate group while primary AML samples and PDXs were generally similar (Figures 1C, S3-S4). Nonetheless, despite the divergence of global chromatin architecture, SEs linked to key lineage TFs tended to be preserved across the sample types (Figure S1B), indicating that cell lines may be acceptable models of core regulatory circuitry.

In agreement with a prior report (*24*), unsupervised clustering of leukemias in the SE space did not correlate with driver mutations except for samples carrying a KMT2A (MLL) translocation, which formed 2 related clusters almost exclusively composed of KMT2A-rearranged (KMT2Ar) leukemias (Figures 1D, S3-S4). A similar pattern was observed in cell lines (Figures 1C, S5), suggesting unique and reproducible differences in the transcriptional circuitry imparted by the KMT2A translocations. We attempted to model these divergent regulatory networks by calculating the connections (cliques) between SE-associated TFs (*10*), but this method resulted in a high number (>400) of potential core TFs within the AML spectrum (Figures S1D-F, S6), consistent with reported observations in other lineages (*49, 50*).

### Selective AML dependencies are strongly associated with SEs, define the AML CRCs

We reasoned that the AML core regulatory circuitry could be more specifically identified by asking which TFs are selectively required for AML survival (*12, 18*). We accessed the data from the Broad Cancer Dependency Map project, which includes genome-scale CRISPR-Cas9 loss-of-function screens of 18,333 genes in 769 cell lines, including 20 AML lines (*51–53*), allowing us to distinguish AML-specific vulnerabilities from universally essential housekeeping genes. We used a skewed-LRT test (*54*) to compare guide RNA drop out between AML and all other cell lines and identified 225 genes selectively essential in AML (Figure S7). Remarkably, while only a minority of SE-driven TFs were essential, >80% (183 out of 225) of selective AML dependencies were associated with at least one common SE (Figure 1E). The enrichment was the strongest among the highest-ranked SEs and precipitously diminished with decreasing average SE rank (Figure 1F). Henceforth, we defined top AML CR TFs as those that were selectively essential in AML and associated with common AML SEs. We defined common SEs as those that were ranked in the top 2250, corresponding to 77% dependency recall, thus prioritizing SEs with the strongest dependency association (Figure 1E). This approach resulted in 24 top CR TFs. Seven additional proteins did not meet one of the criteria but were included based on manual review of the data and validation experiments (Figures 1G, S8). We arbitrarily chose 11 TFs for validation in a low-throughput CRISPR/Cas9 drop out experiment. The observed patterns of growth inhibition generally matched the guide RNA drop out in the genome-wide screen (Figure S7E).

### Divergent CRCs mark context-specific transcriptional addictions

We hypothesized that the distinct SE patterns of the epigenetically defined leukemia clusters were established by divergent CRCs. Furthermore, since SE association is predictive of dependency, CR TFs selectively marked by strong SEs in distinct AML clusters should be predictive of cluster-specific vulnerabilities. Indeed, the activities of CR TF-associated SEs correlated with dependency (Figure 2A-B) and were generally more predictive of dependency than mRNA expression (Figure 2C). We identified distinct CRC signatures of the 6 SE-defined AML clusters. Leukemias carrying KMT2A rearrangements displayed significantly higher SE activities associated with IRF8, MEF2D, RUNX2, HOXA9, CEBPA, FOSL2 and MEIS1 (Figures 2D, S9A), which we termed KMT2Ar-specific CR TFs. As predicted, most of these CR TFs were associated with stronger dependency in KMT2Ar cell lines (Figure 2E). Conversely, RUNX1, ZFPM1 and GATA2 were characterized by higher SE activity and drop out scores in non-KMT2A cell lines. Interestingly, while MEF2C is a recognized dependency in KMT2Ar AML (*55–58*), it appears to have only a slight preference for KMT2Ar leukemias, while IRF8, MEF2D, RUNX2, HOXA9 and MEIS1 are strongly selective (Figure 2E). The CRC signature associated with KMT2Ar translocations was conserved between primary leukemias, PDXs and cell lines (Figure S9B).

**Figure 2.**
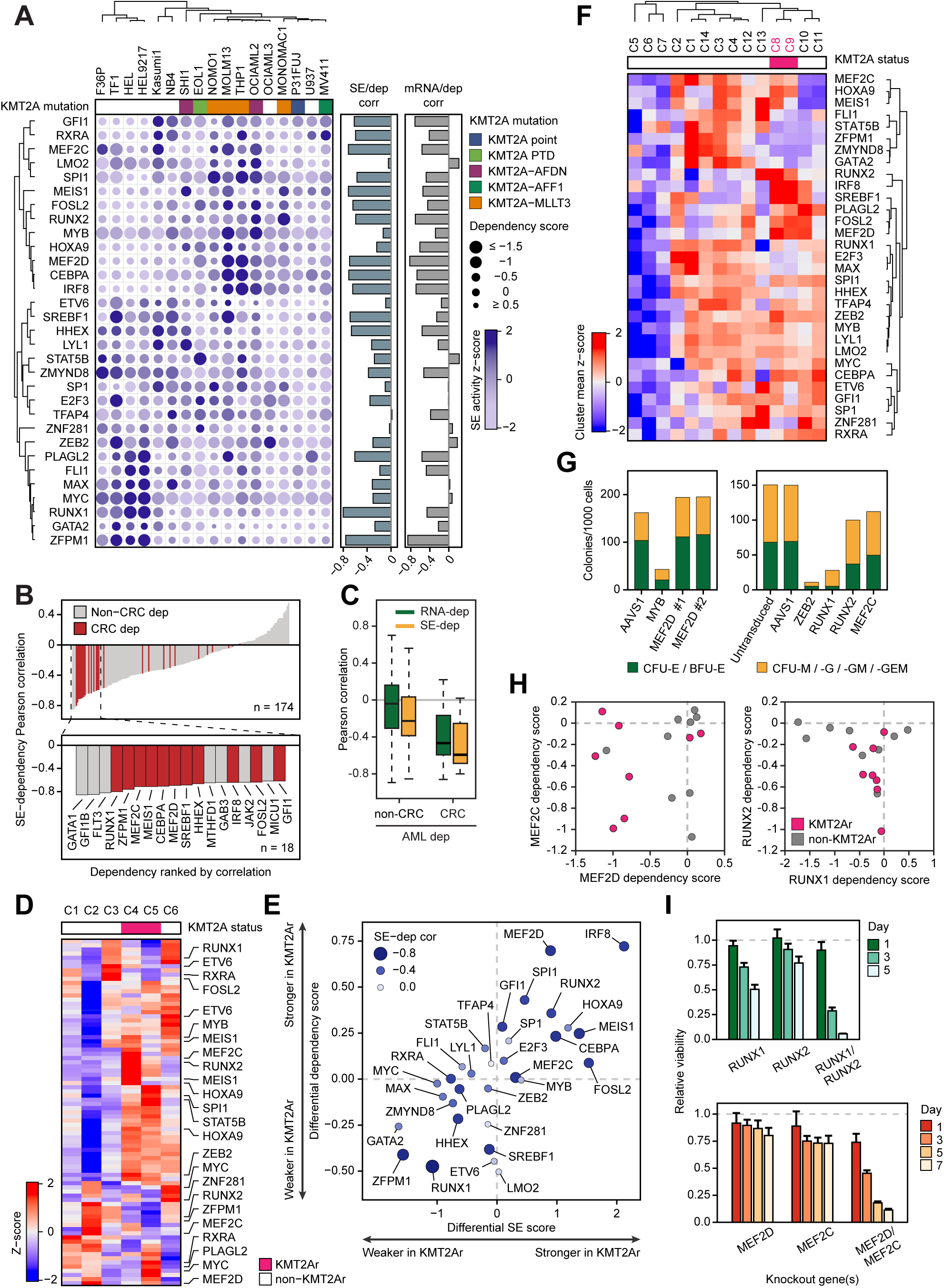
Divergent CRCs are diagnostic of context-specific AML vulnerabilities. (A) A bubble plot of AML CR TFs where color density represents associated SE scores in AML cell lines and bubble size represents dependency scores (negative scores correspond to stronger dependency). The columns and rows are hierarchically clustered according to Pearson correlation of SE scores. Side bars reflect correlation between SE scores and dependency, and mRNA expression and dependency, respectively. (B) SE-associated selective AML dependencies are ranked according to Pearson correlation between SE scores and dependency. CR TFs are highlighted in red. (C) Pearson correlation between SE scores and dependency scores, *versus* mRNA expression and dependency scores. (D) CR TF-associated SE signatures of epigenetically defined AML clusters. Cluster structure is the same as in Figure 1D. The heatmap visualizes average z-scores of CR TF-associated SEs in each cluster. Note that most CR TFs are associated with >1 SE; a full version of the plot is found in Figure S9A. (E) A differential plot of average SE scores versus average dependency scores in KMT2Ar *versus* non-KMT2Ar cell lines. (F) Patients from the BeatAML study (*27*) are hierarchically clustered based on Pearson correlation of CR TF mRNA expression. The heatmap visualizes average z-scores of CR TF mRNA expression in each cluster. A similarity matrix and a sample-level cluster map are found in Figure S10. (G) Human bone marrow-derived CD34^+^ cells from healthy donors were electroporated with *in vitro* assembled Cas9/sgRNA complexes targeting common and subtype-restricted CR TF as shown and plated on cytokine-supplemented methylcellulose media. Colonies were counted following a 14-day incubation period. An AAVS1 (“safe harbor”) targeting sgRNA was used as a control. (H) Scatter plots of paralog DepMap dependency scores in AML cell lines. Negative scores correspond to stronger dependency. (I) Synthetic lethality of TF paralogs. MV411 cells were electroporated with *in vitro* assembled Cas9/sgRNA complexes targeting one or both TF paralogs as shown, and cell viability was measured relative to an AAVS1 (“safe harbor”) control by quantification of ATP pools using a luciferin-based assay.

The SE activities associated with CR TF correlated with their mRNA expression (Figure 2A). Indeed, several of the KMT2Ar-specific CR TFs have been previously noted to be overexpressed in KMT2Ar leukemias (*58–63*). This prompted us to analyze patterns of CR TF expression in BeatAML, a large dataset of 510 genetically annotated and mRNA-sequenced primary AMLs (*27*). Hierarchical clustering of BeatAML samples by pairwise correlation of CR TF expression revealed 14 major clusters. Leukemias carrying KMT2A rearrangements formed two related clusters, C8 and C9, co-segregating with many leukemias having no major chromosomal rearrangements (Figures 2F, S10). High expression of IRF8, MEF2D, and RUNX2 was largely restricted to the KMT2Ar clusters, while expression of MEF2C, HOXA9, MEIS1 and SPI1 was more widely distributed (Figures 2F, S10B). Overall, analysis of mRNA expression revealed a CR TF expression signature that matched the SE signature of KMT2Ar AML and highlighted a large cohort of KMT2Ar-like leukemias. Thus, our findings indicate that differential SE patterns are broadly diagnostic of subtype-specific genetic vulnerabilities and reflect a functionally significant divergence of CRC structure in AML which can largely be reproduced from gene expression data.

We further characterized the 3 most strongly KMT2Ar-restricted CR TFs IRF8, MEF2D and RUNX2, whose roles in KMT2Ar leukemia had not been previously described. First, we confirmed the selective dependency of KMT2Ar leukemia on these TFs by measuring cell viability after CRISPR-Cas9 TF knockout in a panel of 6 cell lines. As expected, depletion of either IRF8, MEF2D or RUNX2 selectively inhibited cell lines carrying KMT2A translocations (Figure S9C). This prompted us to hypothesize that they were leukemia-specific transcriptional addictions (*25, 29*). Indeed, knockout of MEF2D in human CD34^+^ cells had no effect on myeloid colony formation, consistent with the published observation of normal hematopoiesis in MEF2D-null mice (*64*). Similarly, knockout of RUNX2 had little effect on colony formation (Figure 2G). In contrast, depletion of MYB, RUNX1 or ZEB2 resulted in a near-complete loss of colonies. While colony-forming assays are crude models of normal hematopoiesis, we conclude that KMT2Ar-specific CR TFs are relatively dispensable when compared to essential myeloid TFs and represent specific transcriptional addictions of KMT2Ar leukemia.

### Synthetic lethality reveals redundancy of co-expressed TF paralogs

The AML CRC includes two pairs of closely related TF paralogs, RUNX1/2 and MEF2C/D, characterized by highly variable dependency patterns. While RUNX2 and MEF2D are primarily KMT2Ar-specific addictions, dependency on these TFs is not universally associated with KMT2A translocations, and some cell lines are not critically dependent on either TF paralog (Figure 2H). We hypothesized that these paralogs were functionally redundant, allowing them to cross-compensate for each other’s loss. Indeed, simultaneous knockouts of MEF2C/D and RUNX1/2 were synthetically lethal in a cell line that displayed only moderate growth inhibition when either one paralog was depleted (Figure 2I).

### CRC hierarchy revealed by patterns of genomic co-occupancy and knockout response

We sought to investigate further the functional relationships between the members of the AML CRC, using as a model a cell line carrying a KMT2A-AFF1 (MLL-AF4) translocation and a FLT3-ITD mutation. First, we examined the genomic occupancy of 29 of the 31 CR TFs, as well as 7 additional TFs, 6 histone marks and 6 cofactors, by ChIP-seq. When possible, we conducted 2 or more replicate experiments using different primary antibodies and called high-confidence consensus peaks. Two proteins, MEF2D and GFI1, were pulled down using a FLAG affinity tag knock-in after we failed to produce high quality data with primary antibodies. Collectively, our dataset encompassed 109 ChIP-seq experiments representing 48 chromatin-bound proteins. We included published datasets for 6 additional proteins obtained in the same cell line. We detected broad co-occupancy of CR TFs in DNA elements, including promoters and enhancers (Figures 3, S11-S15). Indeed, among the 142,970 co-occupant sites containing 2 or more CR TFs, sites containing >10 CR TFs were significantly over-represented, with many sites harboring >20 co-binding CR TFs (Figure S12C). While most CR TFs demonstrated strong occupancy at the promoters, several TFs (FOSL2, HOXA9, GATA2, CEBPA) were selectively enriched at enhancers (Figure 3). We identified no specific SE signature. The TF paralogs RUNX1 and RUNX2 displayed highly concordant binding, while MEF2D and MEF2C demonstrated partially divergent patterns of chromatin occupancy. No cognate motifs were readily identified in approximately half of the CR TF binding (Figure S12D), and most TFs showed some degree of discordance between motif enrichment and TF binding across clusters (Figure 3). For example, RUNX factor binding was equally prevalent at enhancers and promoters but appeared to be less dependent on the RUNX motifs at the promoters. Overall, motif enrichment appeared to be significantly more focused than TF binding, with many enhancer clusters displaying strong enrichment for one or a few CR TF motifs. This is consistent with a hierarchical structure of TF binding at regulatory elements, where one or a few “founder TFs” nucleate an enhancer, causing additional TFs to be recruited in a cooperative fashion (*65, 66*).

**Figure 3.**
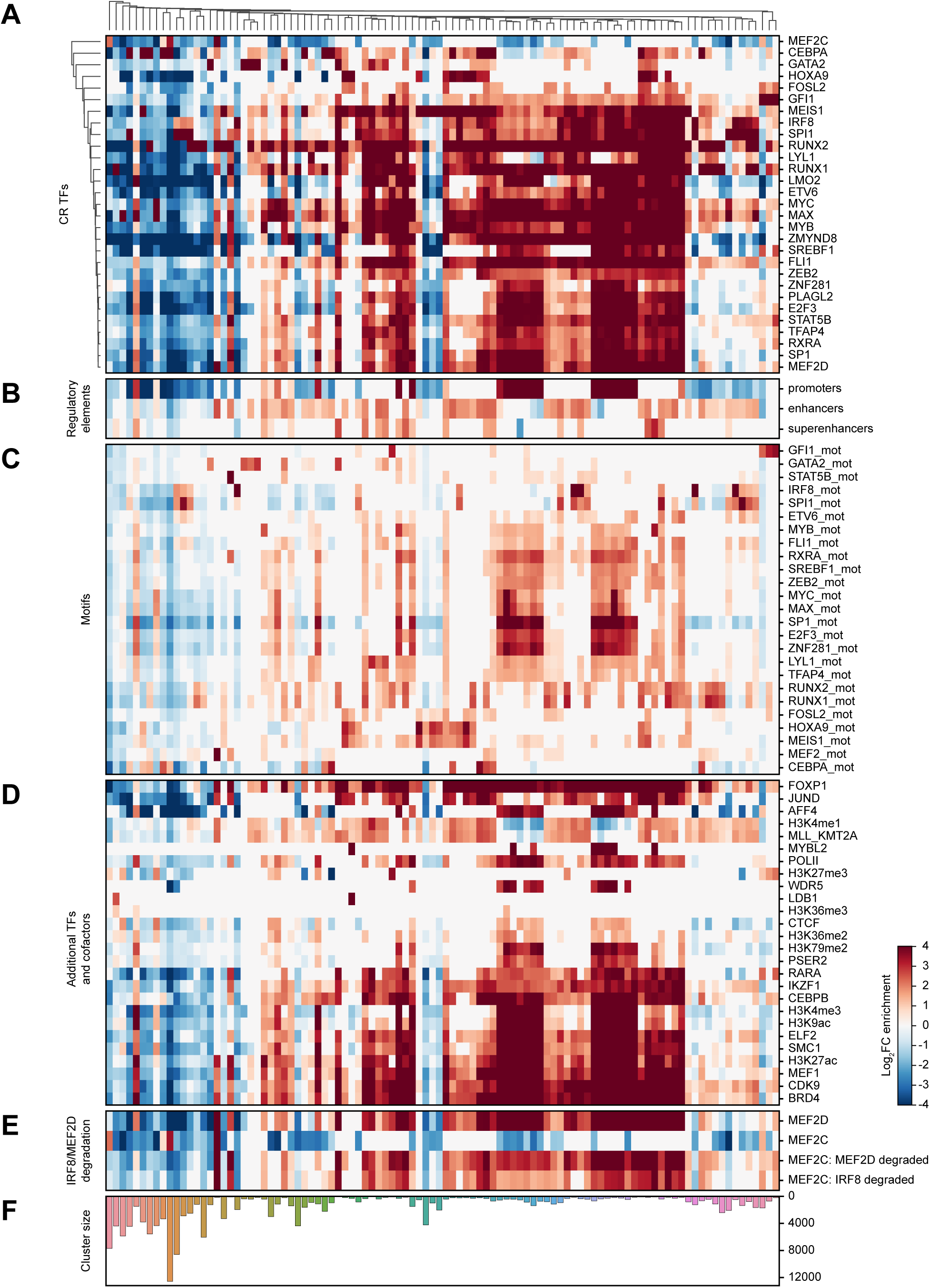
Genomic occupancy patterns of CR TFs reveal binding codes and hierarchy. A co-binding matrix of 142,970 binding sites in MV411 cells was created by merging CR TF binding peaks and eliminating sites with only one CR TF bound. The matrix was *k*-means clustered (*k*=100) on Pearson correlation of binary TF binding, using only the binding data for the 29 CR TFs (960,343 total binding events). Heatmap visualizes enrichment of TF binding and other features by cluster. Note that the white color represents average binding (i.e. neither positive or negative enrichment) rather than no binding. (A) Enrichment of CR TF binding. (B) Enrichment of regulatory DNA elements. (C) Enrichment of motifs corresponding to the CR TFs. (D) Enrichment of additional chromatin-associated proteins. (E) Related to Figures 6-7. Dynamics of MEF2D/MEF2C binding before and after MEF2D and IRF8 degradation. (F) Number of distinct co-binding sites in each cluster.

We examined the functional consequences of CR TF depletion by inactivating 17 CR TFs by CRISPR/Cas9 editing and measuring the transcriptional response by RNA-seq (Figure 4A, S16). First, we sought to evaluate the degree of global transcriptional response following each TF knockout by normalizing transcript counts to external spike-in controls (*67, 68*). Knockouts of key TFs at the top of the functional hierarchy (notably, MYC, SPI1, MAX and MYB) resulted in a global transcriptional collapse (Figure 4B). In contrast, knockouts of HOXA9, FLI1, SP1, RUNX1 and RUNX2) caused a modest net increase in transcription. Simultaneous knockouts of RUNX1/2 and MEF2C/D caused a profound decrease in global transcription, consistent with their synthetic lethality and confirming partially redundant functions of these paralogs. Similar trends were observed on quantitative measurements of genome-wide histone acetylation following the CR TF knockouts, consistent with a global collapse of the transcriptional machinery (Figure 4C).

**Figure 4.**
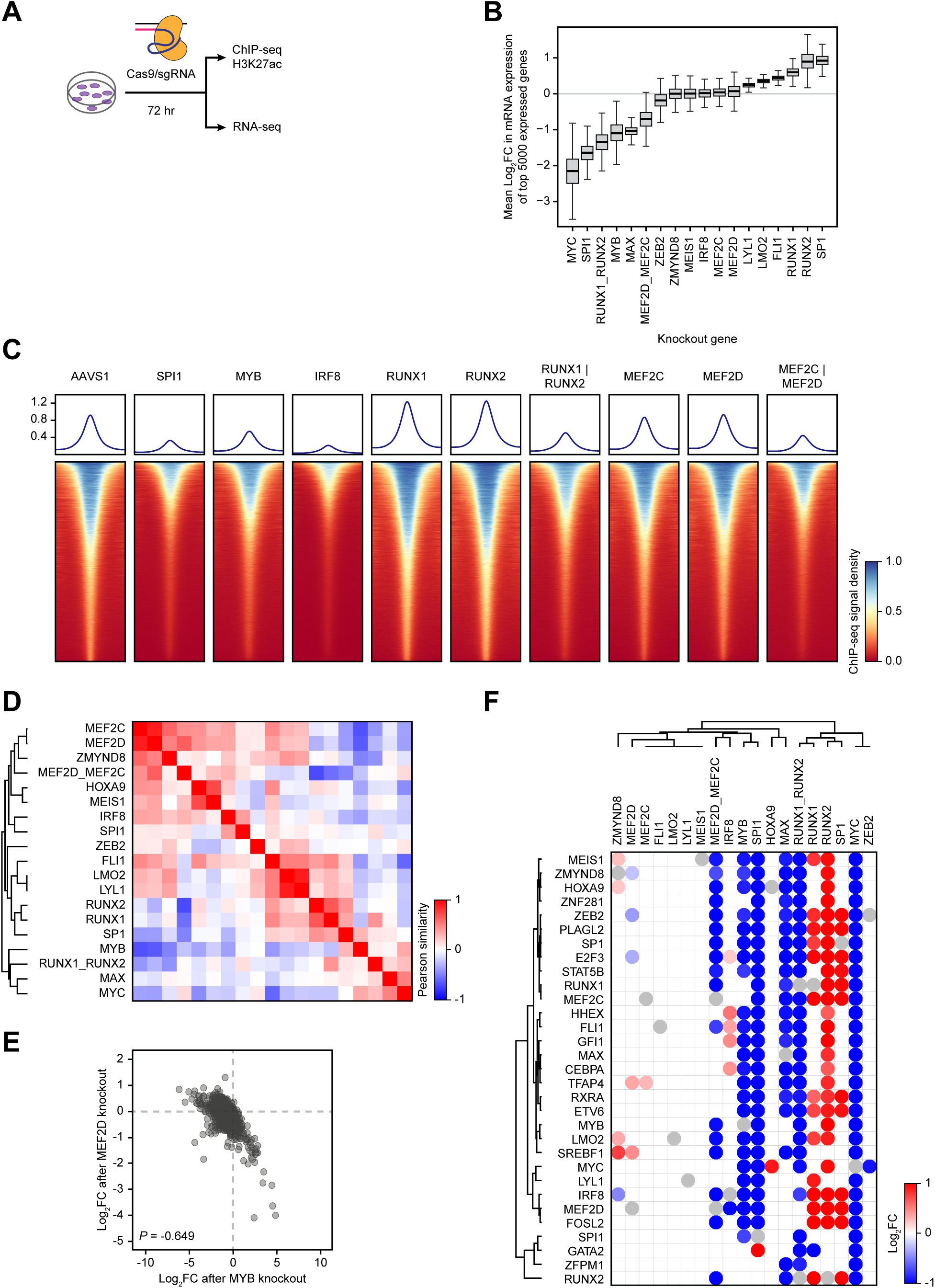
Global transcriptional effects of TF knockout reveal a hierarchical, modular and redundant CRC structure. (A) Experimental design. MV411 cells were electroporated with *in vitro* assembled Cas9/sgRNA complexes targeting 17 individual CR TFs as well as two simultaneous knockouts of TF paralogs. An AAVS1 (“safe harbor”) targeting sgRNA was used as a control. Genome-scale mRNA expression was measured by RNA-seq normalized to an external spike-in control. ChIP-seq with external spike-in control was used to measure changes in H3K27 acetylation. Validation of knockout efficiency by Western blot is found in Figures 6H (MEF2D/MEF2C) and S16. (B) Changes in the expression of top 5000 expressed genes after designated TF knockouts. Each TF knockout is compared to the AAVS1 control. (C) Changes in the genome wide H3K27ac levels after designated TF knockouts, measured by quantitative ChIP-seq using an external spike-in control. Density plots depict genome-wide histone acetylation after indicated TF knockouts. Each row visualizes spike-in normalized ChIP-seq signal around a single H3K27ac peak. (D) Pearson similarity matrix of TF knockouts hierarchically clustered by correlation between knockout-induced changes in the expression of the top 5000 expressed genes compared to the AAVS1 control. (E) Antagonistic actions of MEF2D and MYB are illustrated by cross-plotting transcriptional responses of the top 5000 expressed genes to the MYB and MEF2D knockouts. (F) A matrix of connectivity between CR TFs. Columns represent TF knockouts and rows represent response of the CR TF mRNA to the knockout, expressed as log2 fold change from the AAVS1 control. Only significant changes are shown with genome-wide adjusted *p*-value <0.05. Self-regulation cannot be evaluated by CRISPR knockout and is marked by grey circles.

Next, we asked whether CR TFs can be grouped into functional modules based on the patterns of co-regulated target genes. Indeed, unsupervised clustering of knockout-induced mRNA changes revealed a modular structure of TF-target relationships, where the CR TFs formed three partially antagonistic modules (Figure 4D). The KMT2Ar-specific CR TFs MEIS1, MEF2C, MEF2D and IRF8 formed one module along with ZMYND8. The module was strongly antagonistic with another module including MYB and MYC, with a particularly strong antagonism between the MEF2 factors and MYB (Figure 4E). Knockouts of TF paralogs had similar effects, consistent with partially redundant functions (Figure S16B). Similarly, TF pairs known to form chromatin-bound complexes (IRF8/SPI1, MYC/MAX, LYL1/LMO2 and HOXA9/MEIS1) showed similar patterns of chromatin binding and concordant transcriptional effects, consistent with their reported functional and structural associations (*69–77*).

Finally, we assessed the connectivity within the CRC and asked how each TF influences expression of the other members of the circuit (Figure 4F). Consistent with the genome-wide trends, loss of MYC, MAX, MYB and SPI1 resulted in a broad collapse of nearly the entire circuit, while other CR TFs affected few or no other members. The observation of limited CRC connectivity contrasts with the accepted model of CRC regulation where each member TF regulates expression of all other members (*10*).

### The IRF8/MEF2 axis regulates key oncogenes in KMT2Ar AML

We focused attention on the two most KMT2Ar-specific CR TFs IRF8 and MEF2D, whose overexpression has been associated with adverse outcomes (Figure S21J-K). The dependency scores and transcriptional effects of these two TFs display a high degree of correlation (Figures 2A, 4D, S7B, S17A), suggesting a functional link. However, measurements of mRNA pools after a gene perturbation cannot distinguish primary TF targets from secondary effects (*78*). To identify genes directly regulated by MEF2D and IRF8 we employed a targeted protein degradation strategy (*79, 80*). We constructed two AML cell lines expressing MEF2D and IRF8, respectively, fused to the FKBP12^F36V^ (dTAG) domain by a homozygous knock-in of the FKBP12^F36V^-coding DNA sequence into the endogenous TF-coding loci (Figures 5A-C, 7A-C). Treatment with dTAG^V^-1, a heterobifunctional small molecule that engages FKBP12^F36V^ and VHL (*80*), led to a near-complete loss of the fusion protein within 2 hours. The effects of TF degradation on cell viability and mRNA pools were quantitatively similar to the effects seen after CRISPR-Cas9-mediated gene knockouts (Figure S17B-E). Following degradation of MEF2D and IRF8 for 2 and 24 hours we measured the genome-wide rates of nascent mRNA synthesis by SLAM-seq (*81, 82*).

**Figure 5.**
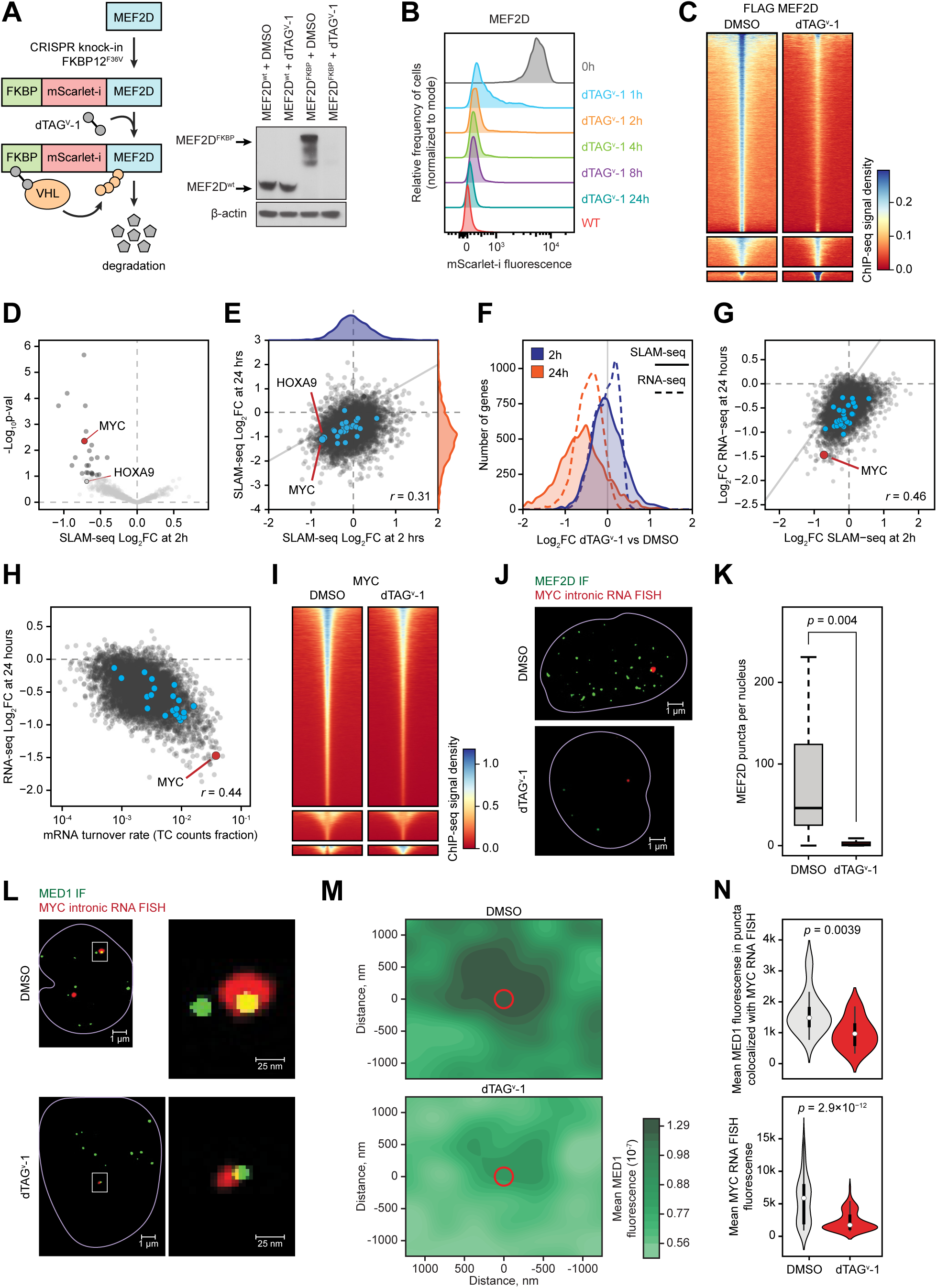
Direct transcriptional effects of MEF2D revealed by targeted degradation and SLAM-seq. (A) Schematic and Western blot of endogenous MEF2D tagging by CRISPR-HDR and subsequent targeted degradation of the fusion protein. (B) A time course of MEF2D degradation by FACS measurement of the fusion protein fluorescence. (C) Degradation of MEF2D reduces its genomic occupancy, as demonstrated by density plots of spike-in controlled anti-FLAG MEF2D ChIP-seq experiment showing genome-wide occupancy change after MEF2D degradation. Each row represents a single peak. (D) A volcano plot of genome-wide changes in nascent mRNA transcription measured by SLAM-seq after 2 hours of MEF2D degradation. (E) A cross-plot of genome-wide changes in nascent mRNA transcription measured by SLAM-seq after 2 *versus* 24 hours of MEF2D degradation demonstrates a poor correlation between early and late transcriptional responses, as well as signs of transcriptional collapse by 24 hours. (F) A distribution plot of genome-wide changes in nascent transcription rates (SLAM-seq) and mRNA pools (RNA-seq) after 2 and 24 hours of MEF2D degradation. (G) Correlation between changes in nascent RNA transcription (SLAM-seq) after 2 hours of MEF2D degradation *versus* changes in the mRNA pools (RNA-seq) after 24 hours of MEF2D degradation. (H) Correlation between steady state mRNA turnover rates approximated from the SLAM-seq TC count fraction *versus* changes in the mRNA pools (RNA-seq) after 24 hours of MEF2D degradation. (I) A density plot of spike-in controlled anti-MYC ChIP-seq experiment demonstrating reduced genome-wide MYC occupancy after MEF2D degradation. (J) Degradation of MEF2D for 2 hours results in a dramatic reduction of MEF2D puncta in the nucleus. Green color represents MEF2D immunofluorescence (IF) signal. Red color represents intronic MYC RNA FISH signal. (K) Quantitative analysis of MEF2D puncta after degradation. (L) Degradation of MEF2D for 2 hours results in decreased mediator recruitment and reduced MYC transcription. Green color represents MED1 IF signal. Red color represents intronic MYC RNA FISH signal. Yellow color represent overlap between red and green signal. (M) A density plot of multi-image analysis showing reduced mediator recruitment to foci of MYC transcription after degradation of MEF2D for 2 hours. The green color gradient represents kernel density estimation of aggregate MED1 IF signal in a cubic region of 1400 nm^3^ centered on the MYC RNA FISH condensates in each cell. The red circle represents the average size of the MYC RNA FISH puncta. (N) Quantitative analysis of MED1 puncta before and after degradation demonstrating reduced mediator fluorescence at the sites of MYC transcription after MEF2D degradation for 2 hours.

Degradation of MEF2D for 2 hours resulted in a significant decrease in transcription of only 23 genes (adjusted *p*-value <0.1; Figures 5D, S18-19), representing high-confidence direct MEF2D targets. Of these genes 7 encoded TFs (MYC, BHLHE40, KLF10, KLF6, KLF11, ZBTB33 and ZATB2) and 2 (MYC and TP53RK) were essential genes with dependency probability >0.5 (Figure S19A). Two additional CR TFs (ZEB2 and HOXA9) had borderline statistical significance at 2 hours and became significant by 24 hours, likely also representing direct MEF2D targets (Figure S19B). By 24 hours of MEF2D deprivation a global decrease in transcription was evident, consistent with a global transcriptional collapse (Figure 5E-G). However, the changes in the transcription rates of individual genes seen after 24 hours of MEF2D deprivation correlated poorly with the early changes, indicating that the majority of the late transcriptional responses were secondary events. Thus, although at least 8 CR TFs were significantly affected at 24 hours, including IRF8 and MEF2D itself, these appear to be secondary effects rather than direct MEF2D targets. The global decrease in mRNA pools by 24 hours, measured by RNA-seq, lagged behind the changes in transcription rates and correlated with mRNA turnover rates (Figure 5H).

Given the central role of MYC as a “transcriptional amplifier” (*83, 84*), reduced MYC activity, evidenced by decreased MYC binding across the genome after MEF2D degradation (Figure 5I), likely contributed to the global transcriptional collapse. Indeed, we found the MYC transcriptional signature to be strongly enriched among the genes affected by the MEF2D loss (Figure S20A). Furthermore, indirect inhibition of MEF2 function via inhibition of SIK3 by the small-molecule tool compound YKL-05-099 (*56*) had a similar direct transcriptional signature, including reduced MYC transcription, and demonstrated a modest synergy with pharmacologic MYC inhibition (Figure S20B-D). However, forced expression of MYC failed to rescue the growth phenotype of MEF2D degradation (Figure S20E-H), consistent with the observation that direct MEF2D targets include other essential genes.

We sought additional confirmation of a direct role of MEF2D in the regulation of MYC transcription. Utilizing super-resolution structured illumination microscopy (SIM) immunofluorescence (IF), we visualized MEF2D puncta in the nucleus. The MEF2D puncta co-localized with the sites of active MYC transcription highlighted by concurrent fluorescent *in situ* hybridization (FISH) for nascent MYC RNA (Figures 5J, 6B). We asked whether MEF2D regulated MYC expression by facilitating mediator recruitment. Indeed, degradation of MEF2D resulted in reduced intensity of MED1 puncta, along with a reduced intensity of the nascent RNA FISH signal, consistent with impaired MYC transcription (Figure 5J-N). These results confirmed that MEF2D directly activates MYC transcription by promoting mediator recruitment.

**Figure 6.**
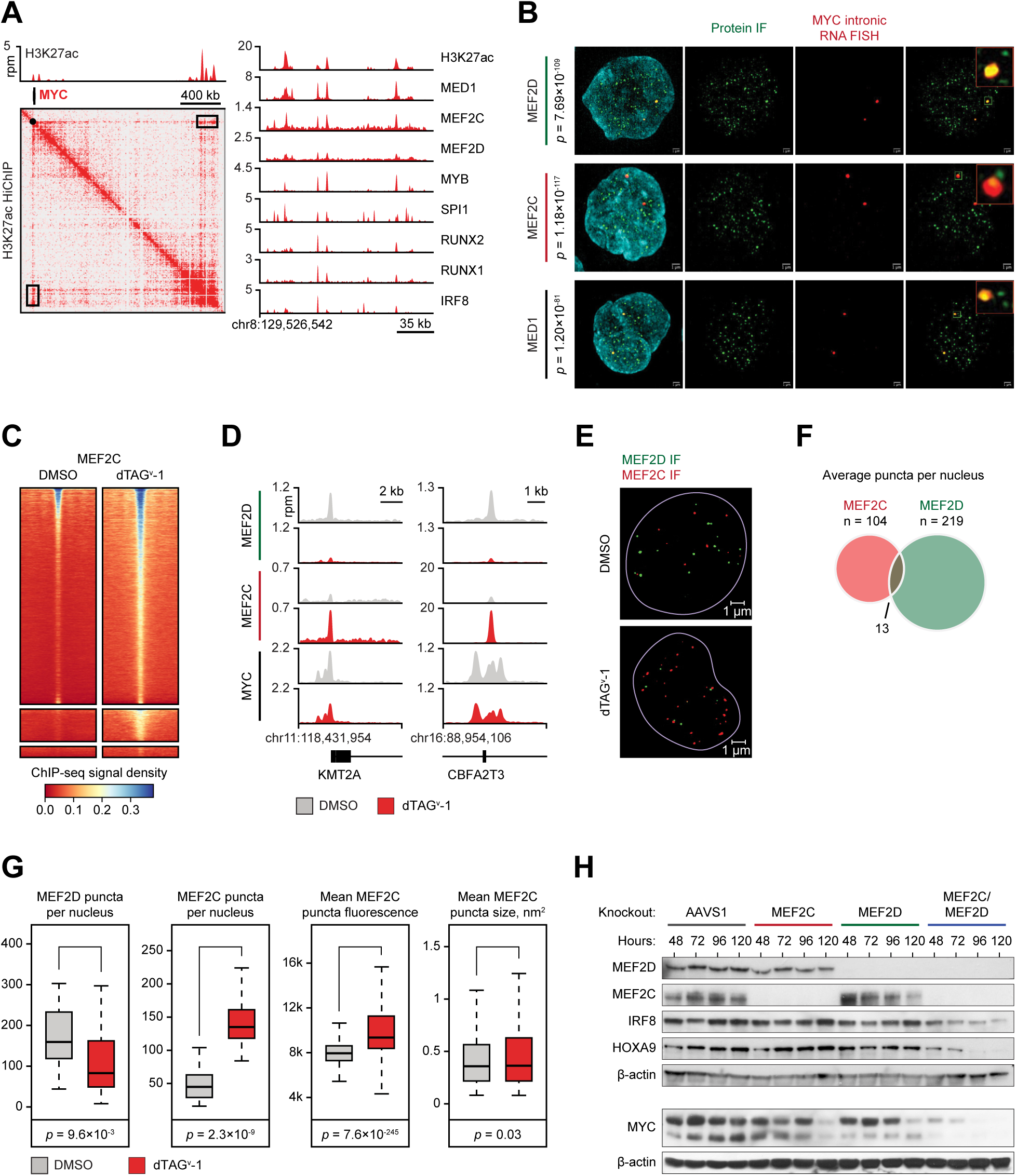
Compensation and competition between MEF2D and MEF2C. (A) A 2D HiChIP plot illustrating H3K27ac-mediated DNA contacts in the MYC locus and ChIP-seq tracks of CR TF binding at the MYC SE located ∼1.8 Mb downstream of the MYC promoter (black box on the HiChIP map). (B) SIM super-resolution confocal microscopy of MV411 cells with simultaneous immunofluorescence using primary antibodies against the designated proteins and intronic RNA FISH targeting nascent MYC transcripts. *p*-values reflect significance of co-localization of protein puncta with RNA FISH calculated by Fisher exact test. (C) A density plot of spike-in controlled anti-MEF2C ChIP-seq experiment demonstrating increased MEF2C occupancy after MEF2D degradation. Each row represents a single peak called from an anti-FLAG MEF2D ChIP-seq experiment in unmanipulated MV411 cells. Color gradient reflects MEF2C ChIP-seq signal in the MEF2D peaks. (D) ChIP-seq tracks demonstrating changes in MEF2D, MEF2C and MYC binding in two representative loci after MEF2D degradation. (E) MEF2D degradation results in increased number and intensity of MEF2C nuclear puncta. The images demonstrate immunofluorescence with antibodies against MEF2D (green) and MEF2C (red) before and after MEF2D degradation. (F) Overlap between MEF2D and MEF2C puncta in unperturbed cells on multi-image analysis. (G) Quantitative multi-image analyses of MEF2D and MEF2C puncta before and after MEF2D degradation shows increased number and intensity of MEF2C condensates after MEF2D degradation. (H) Western blot demonstrating changes in CR TF protein levels after single and combined MEF2D/MEF2C knockouts by CRISPR/Cas9.

We hypothesized that the modest immediate transcriptional response to MEF2D loss was due to partial compensation by MEF2C. Indeed, both paralogs localized to the MYC locus on SIM IF-FISH and ChIP-seq (Figure 6A-B). Remarkably, degradation of MEF2D caused redistribution of MEF2C to the genomic sites vacated by MEF2D (Figure 3E, 6C-D). Similarly, MEF2C nuclear puncta, which only partially overlapped with MEF2D puncta on SIM IF, increased in number and intensity following MEF2D degradation, despite a modest decrease in the total MEF2C level (Figures 6E-G, S18E). These results are consistent with genome-wide competition between MEF2D and MEF2C for chromatin binding and provide a mechanistic basis for their partially redundant functions. Consistent with these observations, simultaneous knockouts of MEF2D and MEF2C resulted in a more profound loss of MYC and HOXA9 compared to knockouts of either paralog alone (Figure 6H).

Targeted degradation of IRF8 for 2 hours resulted in a significant loss of MEF2D transcription, providing an explanation for the functional linkage between the two TFs (Figure 7A-E). Although MEF2D was the only affected TF, direct targets of IRF8 included 71 additional genes, of which 6 were essential, including the BCL2 oncogene (Figures S21-22). We found a single strong IRF8 binding site in the MEF2D SE (Figure 7F), prompting us to hypothesize that IRF8 activated MEF2D transcription by binding at this locus. We excised a ∼340 bp DNA sequence containing the IRF8 binding motif in the cell line carrying the FKBP-mScarlet-MEF2D fusion and measured mScarlet fluorescence as a reporter for MEF2D expression. Loss of the IRF8 binding sequence resulted in reduced fluorescence, indicating that the SE segment containing the IRF8 binding site is essential for maintaining MEF2D expression (Figure 7G). Notably, concurrent depletion of MEF2D and MEF2C resulted in decreased IRF8 levels (Figure 6H), while prolonged depletion of either IRF8 or MEF2D reduced the levels of MEF2C (Figures S18E, S21H). However, the absence of similar changes on SLAM-seq after rapid TF degradation indicated that these connections were indirect. We conclude that IRF8 directly activates transcription of MEF2D, while direct targets of MEF2D include key leukemogenic TFs HOXA9 and MYC (Figure 7H). Direct targets of both TFs include other essential genes, including IRF8 regulation of BCL2, an important therapeutic target in AML (*85*). Remarkably, both TFs directly regulate relatively small, non-overlapping sets of target genes, and most of the transcriptional changes detected by RNA-seq after a TF knockout represent secondary effects.

**Figure 7.**
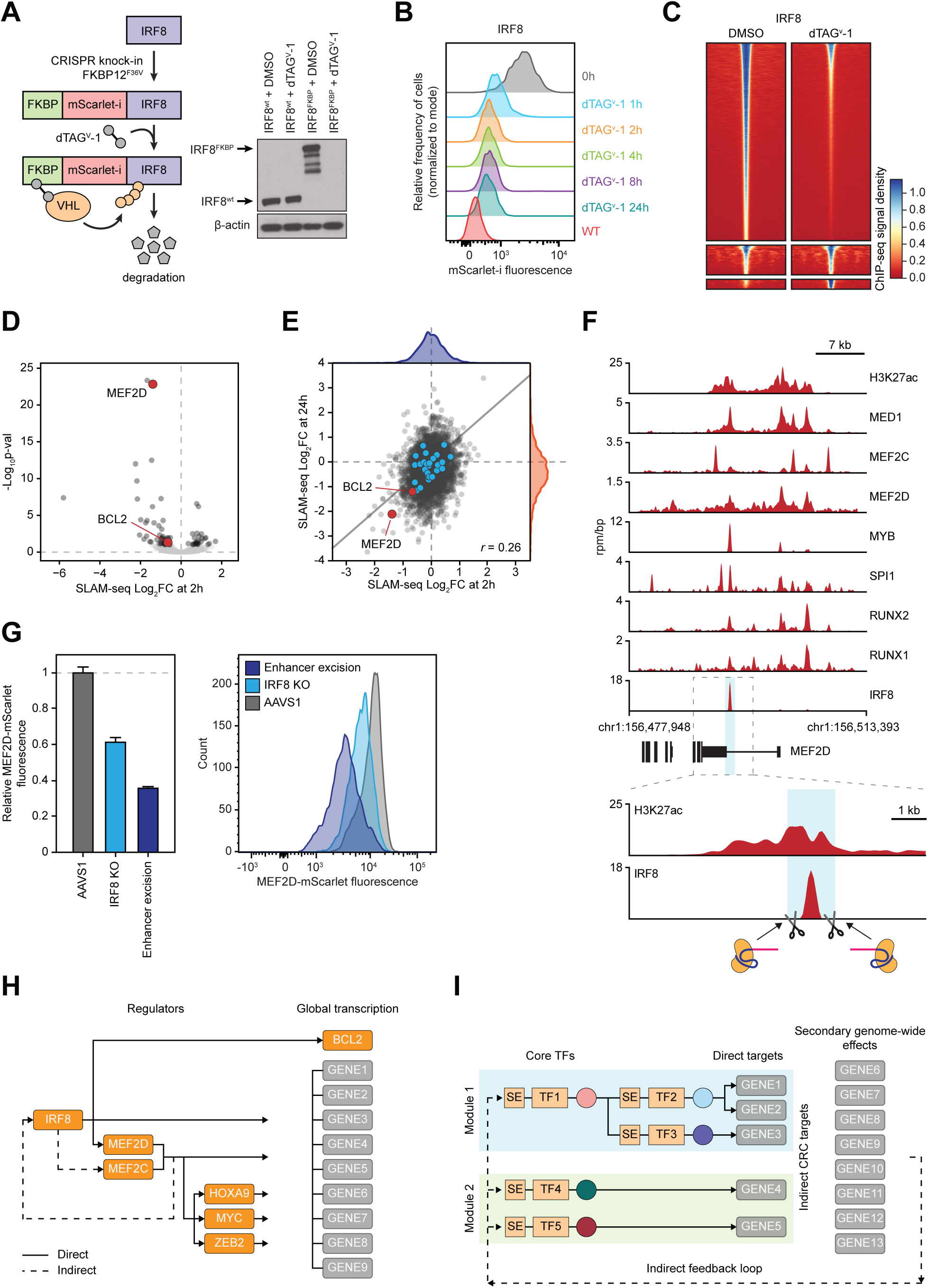
IRF8 directly regulates MEF2D. (A) Schematic and Western blot of endogenous IRF8 tagging by CRISPR-HDR and subsequent targeted degradation of the fusion protein. (B) A time course of IRF8 degradation by FACS measurement of the fusion protein fluorescence. (C) A density plot of spike-in controlled anti-IRF8 ChIP-seq experiment showing reduced genome-wide occupancy after degradation. (D) A volcano plot of genome-wide changes in nascent mRNA transcription measured by SLAM-seq after 2 hours of IRF8 degradation. (E) A cross-plot of genome-wide changes in nascent mRNA transcription measured by SLAM-seq after 2 *versus* 24 hours of IRF8 degradation demonstrates a poor correlation between early and late transcriptional response. (F) ChIP-seq tracks of CR TF binding at the MEF2D SE and schematic of CRISPR/Cas9 strategy for IRF8 binding site excision. (G) Changes in the MEF2D protein levels measured by mScarlet reporter fluorescence after IRF8 gene knockout *versus* excision of IRF8 binding site in the MEF2D locus. MV411 cells carrying the FKBP-mScarlet-MEF2D fusion were electroporated with Cas9/sgRNA complexes targeting the IRF8 gene, IRF8 binding site in the MEF2D SE, or AAVS1 control, respectively, and fluorescence was measured by FACS 72 hours after electroporation. (H) Schematic of the direct regulatory relationships in the IRF8/MEF2 axis. (I) Schematic of a modular CRC structure with few direct TF-TF and TF-target connections.

## Discussion

### A transcriptional dependency map for AML

A malignant cell depends on the flow of information from a cancer-initiating mutation, through a presumably small set of transcriptional regulators, to the effector genes whose altered expression drives the malignant phenotype. Accumulating evidence indicates that these cancer-specific transcriptional circuits can be therapeutically targeted (*25, 32, 86–89*). While most transcriptionally directed therapeutics target oncogenic TFs indirectly, there are increasing examples of direct TF inhibition, such as the targeted degradation of IKZF1/3 and ZMYM2 fusion oncoproteins by thalidomide analogs (*90, 91*). Our study provides a comprehensive characterization of AML transcriptional dependencies obtained from a combination of systems biology and mechanistic approaches. By restricting our CRC model to selectively essential TFs we focus on the circuits selectively critical for AML survival and thus most attractive for therapeutic reprogramming efforts. While many of them are recognized AML dependencies, such as MYB, SPI1, HOXA9, MEF2C and ZEB2 (*37, 56, 57, 77, 92–94*), others had not been functionally characterized in the context of AML. Our datasets and analyses provide a resource that will facilitate development of novel AML therapies.

### General principles of CRC organization

Our study reveals several principles of AML CRC organization that may be extrapolated to other cell types. First, CR TFs form a functional hierarchy. At the top of the hierarchy are universally essential lineage TFs, such as MYB and SPI1. These TFs play central roles in nucleating lineage-specific enhancers and their loss causes a profound transcriptional collapse. At the bottom of the hierarchy are subtype-restricted TFs regulating more narrow transcriptional programs. Second, CRCs are composed of functional modules of collaborating TFs that enforce a common transcriptional program. While each module participates in both gene activation and repression, distinct modules demonstrate largely antagonistic functions. Third, divergent CRCs establish distinct epigenetic subtypes of AML and form the basis of context-specific transcriptional addiction. Finally, CRCs are stabilized by partial functional redundancy of co-expressed TF paralogs. Elements of this organizational model have been reported in other cell types (*18–22, 95–100*).

Our findings have significant implications for our understanding of functional connectivity within a cell’s transcriptional network. In contrast to a widely accepted CRC model, where each member positively regulates its own expression and the expression of all other CR TFs (*9, 10, 101*), we find only partial connectivity within the AML CRC. Furthermore, some edges appear to be inhibitory, and many edges appear to reflect secondary effects of transcriptional collapse rather than direct regulation. This underscores the long-standing observation that only a fraction of TF binding events has functional significance (*102, 103*). Using MEF2D and IRF8, a pair of TFs with highly concordant dependency and transcriptional signatures, as a case study, we find unexpectedly small and non-overlapping sets of direct transcriptional targets. Instead, the functional similarity between MEF2D and IRF8 stems from the direct role of IRF8 in the enforcement of MEF2D expression. In addition, despite binding to their own regulatory elements, neither TF displays direct autoregulation. In contrast, the concordant transcriptional effects of SPI1/IRF8, MYC/MAX and HOXA9/MEIS1 appear to be largely driven by cooperativity at the enhancer level. Thus, the functional modules within the CRC structure are defined by a complex interplay of direct and indirect interactions, where direct connections include “vertical” TF-TF regulation and “horizontal” TF cooperativity. While the master myeloid TFs at the top of the functional hierarchy (such as MYB or SPI1) may directly regulate larger sets of genes, we suspect that many TF-TF and TF-target connections in the traditionally derived models of transcriptional networks are in fact indirect or functionally insignificant.

### Paralog redundancy and competition

Functional redundancy of co-expressed TF paralogs has been recognized as one of the mechanisms providing robustness to gene regulatory networks (*93, 104, 105*). For example, compensation by MEF2A attenuates loss of MEF2D in mouse cerebellum (*97*), while RUNX1 and RUNX3 have partially redundant functions in T-cell development (*98*). However, the mechanisms by which TF paralogs exert their similar but not identical functions are not well understood. Our study uncovers partially redundant functions of RUNX1/2 and MEF2C/D in KMT2Ar AML. While RUNX1 and RUNX2 display virtually identical genome-wide occupancy, MEF2C and MEF2D are characterized by partially divergent binding patterns consistent with distinct binding codes. Nonetheless, MEF2C compensates for MEF2D loss by re-locating to the MEF2D binding sites and displaying increased puncta in the nucleus. We suspect that paralog redundancy is a common feature of transcription networks and has important implications for the interpretation of gene dependency screens. It is also a limitation of the functional dependency approach to CRCs, as it restricts our definition of the CR TFs to those TFs whose loss cannot be fully compensated. Indeed, we identified RUNX1/2 and MEF2C/D as CR TFs because they displayed at least a modest effect on cell growth when knocked out alone. However, there are likely additional core TFs with profoundly redundant functions in the context of AML whose functional significance can only be uncovered in synthetic lethality screens. For example, direct targets of MEF2D include 3 KLF paralogs (KLF10, KLF6, KLF11). While none of the individual paralogs are essential in the genome-wide dependency screen, the simultaneous loss of all 3 proteins may be synthetically significant and contribute to the eventual transcriptional collapse triggered by the MEF2D degradation. Paradoxically, inactivating mutations in several essential TFs, such as SPI1/PU.1 and CEBPA, are common drivers of AML (*106*). An intriguing possibility is that their loss may be rescued by their respective paralogs expressed in a specific cellular context. If that were the case, the rescuing paralogs would become *de novo* transcriptional addictions, providing yet another opportunity for therapeutic intervention.

### Dynamic behaviors of TF and mediator puncta

The role of biomolecular condensates in the nucleus continues to be a subject of active research (*107–116*). While our study does not specifically address the role of liquid-liquid phase separation in transcription control, we used super-resolution microscopy as an orthogonal tool to understand TF dynamics. Consistent with prior reports, we demonstrate that TFs and mediator form distinct puncta co-localizing with foci of active transcription (*110, 112*). We show that degradation of MEF2D rapidly reduces transcription and mediator recruitment at the MYC locus. This is consistent with the results of SLAM-seq and provides additional proof of direct regulation of MYC transcription by MEF2D. Our observation of increased MEF2C puncta induced by MEF2D degradation further validates partial redundancy and compensation between the two paralogs. Additional studies are needed to characterize the specific mechanisms of paralog partitioning and competition in the context of chromatin structure.

### A context-specific core regulatory module enforces expression of common leukemia oncogenes

Traditional gene or messenger RNA disruption methods followed by measurements of mRNA pools cannot distinguish primary TF targets from secondary effects due to the slow-onset kinetics of TF deprivation and vast differences in mRNA and protein turnover rates (*78, 117, 118*). Recently, direct TF targets have been inferred from direct measurements of genome-wide transcription rates following rapid TF degradation (*81*). Using this approach, we demonstrate that the IRF8/MEF2 axis directly enforces expression of the key leukemia oncogenes MYC, HOXA9 and BCL2, each of which is a recognized therapeutic target. MYC is one of the most prevalent human oncogenes, while HOXA9 is a common driver of leukemia (*75, 93, 94, 119*). BCL2 is the target of venetoclax, one of the most promising novel therapies for AML (*85*). Our study clarifies the complex reported interactions between IRF8, HOXA9 and MEF2C/D proteins. HOXA9 is overexpressed in >50% of AML cases, and high expression of IRF8, MEF2C, MEF2D and HOXA9 has been linked to adverse outcomes (*75, 120–123*). While MEF2C is a transcriptional dependency of KMT2A-rearranged AML (*55–58*), the role of MEF2D in leukemia has remained relatively unexplored. We demonstrate that IRF8 directly activates expression of MEF2D, which then directly activates MYC and HOXA9 transcription in KMT2Ar AML. In agreement with our findings, the MEF2D paralog MEF2B has been reported to regulate MYC in diffuse large B-cell lymphoma (*124*). Notably, while MEF2D and MEF2C have partially overlapping functions, only MEF2D appears to be directly regulated by IRF8. Indeed, expression of MEF2C is more widely distributed in primary AML samples and correlates with the expression of HOXA9 in non-KMT2Ar AML (Figures 2F, S10B). Thus, MEF2C appears to regulate expression of HOXA9 in non-KMT2Ar leukemias, while the role of MEF2D is restricted to KMT2Ar AML. Notably, ectopic expression of MEF2C has been shown to induce leukemic transformation in *Irf8*^-/-^ mice (*125*). In light of our data, it appears that IRF8 knockout likely resulted in a loss of MEF2D expression, which was then functionally rescued by ectopic expression of MEF2C, promoting leukemogenesis. Our study provides an example of a specific class of driver mutations establishing a sub-network within the CRC hierarchy to enforce expression of common leukemia oncogenes, and an evolving model of CRC organization based on improved understanding of direct transcriptional regulation.

## Acknowledgements

M. Pimkin is supported by a Damon Runyon-Sohn Pediatric Cancer Fellowship and a Young Investigator Award from the Alex’s Lemonade Stand Foundation. This work was supported, in whole or in part, by research grants to M. Pimkin from the Curing Kids Cancer Foundation, When Everyone Survives Foundation, Pedals for Pediatrics, Children’s Cancer Research Fund, Children’s Leukemia Research Association, Hyundai Hope on Wheels, Kate Amato Foundation, and Boston Children’s Hospital Office of Faculty Development. K Stegmaier was supported by NIH 5R35 CA210030 and NIH P50 CA206963. J. M. Ellegast was supported by the Swiss National Science Foundation, the Lady Tata Memorial Trust, and the Pediatric Cancer Research Foundation, and is a Fellow of The Helen Gurley Brown Presidential Initiative (The Pussycat Foundation). N.V. Dharia was supported by the Julia’s Legacy of Hope St. Baldrick’s Foundation Fellowship. We thank Dr. E. Bresnick for sharing a custom-made GATA2 antibody. We thank Dr. R.A. Young for helpful discussions and advice, Drs. I.A. Klein, A. Dall’Agnese and L.S. Li for technical assistance, and Drs. R.A. Shivdasani, S.A. Armstrong and B.L. Ebert for critical reading of this manuscript.

## Author Contributions

Conceptualization: SHO, MPi

Computational methodology development and data analysis: JK, MPi, NVD, KE, MPe, JVK, JMD, GK, JW, AK, QZ, CYL, VRP

Experiments: TH, MPi, YH, JXF, JE, ZTH, JME, JSY, AB, CHL, BN

Visualization: MPe, TH, YH, JK, MPi, JXF

Funding acquisition: SHO, MPi

Supervision: SHO, MPi, KS, DW, CYL, NVD

Writing and review of the manuscript: all authors

## Declaration of Interests

K. Stegmaier has funding from Novartis Institute of Biomedical Research, consults for and has stock options in Auron Therapeutics, and has consulted for Kronos Bio and AstraZeneca on topics unrelated to this work. N.V. Dharia is a current employee of Genentech, Inc., a member of the Roche Group. J. Xavier Ferrucio is a current employee of Vor Biopharma. C.Y. Lin is a current employee of Kronos Bio. B. Nabet is an inventor on patent applications related to the dTAG system (WO/2017/024318, WO/2017/024319, WO/2018/148440, WO/2018/148443 and WO/2020/146250). K. Eagle has consulted for Third Rock Ventures and Flare Therapeutics on topics unrelated to this manuscript. All other authors declare no potential conflict of interest.

## Supplementary Figure Titles and Legends

**Figure S1, related to** **Figure 1. SE landscapes in human AML**

(A) Occurrence of distinct SEs across AML samples, including primary AMLs, PDXs and cell lines.

(B) Comparative analysis of SE ranks between cell lines and primary-derived AML samples (primary AMLs and PDXs) demonstrates overall similarity of SE ranks, including SEs associated with CR TFs (highlighted in red).

(C) Top SEs are enriched for TFs.

(D) Schematic of clique enrichment analysis to identify context-specific CRCs (*10*). For every SE-associated TF_a_, the in-degree was calculated as the number of TFs with a motif present in the nucleosome-depleted H3K27ac “valleys” of the TF_a_’s SE, and the out-degree was calculated as the number of TF SEs with a TF_a_ motif. Clique enrichment scores, defined as the fraction of all cliques within a sample of which that TF is a constituent, identify TFs with the highest network connectivity.

(E) Relationship between clique enrichment scores and samples with the clique for all identified cliques across AML samples (grey dots) and CR TFs defined by intersection of common SEs and selective AML dependencies (red dots).

(F) All TFs with at least one clique in at least one sample are ranked according to the product of median clique score and the number of samples where the clique was identified, with CR TFs highlighted in red. The analyses in (E) and (F) demonstrate generally poor specificity of clique analysis in identifying lineage-essential CR TFs, although selective AML dependencies are generally associated with higher clique enrichment scores (see also Figure S7C).

**Figure S2, related to** **Figure 1. Validation of SE-promoter assignments by dCas9-KRAB-MECP2 interference.** MV411 cells stably expressing dCas9-KRAB-MECP2 were transduced with lentiviral vectors encoding gRNAs targeting the indicated loci. Expression of the predicted target gene was measured by RT-qPCR. ChIP-seq tracks show H3K27 acetylation signal in MV411 cells.

(A) Predicted SE-promoter links in the MYB locus demonstrated by the H3K27ac HiChIP 2D contact map.

(B) Validation of predicted SE-promoter links in the MYB locus. The experiment was performed as one biological replicate.

(C) Predicted SE-promoter links in the IRF8 locus demonstrated by the H3K27ac HiChIP 2D contact map.

(D) Validation of predicted SE-promoter links in the IRF8 locus. The experiment was performed in 3 biological replicates.

**Figure S3, related to** **Figure 1. A SE-based classification of AML.** The heatmap is a full version of the heatmap in Figure 1D. Primary and PDX samples (columns) are hierarchically clustered with complete linkage using Pearson correlation of the scores of 4798 SEs recurrent in at least 2 samples (rows). The bars at the top reflect presence of coding mutations (grey color designates unknown) and sample type.

**Figure S4, related to** **Figure 1. A SE-based classification of AML: similarity matrix.** The heatmap is a similarity matrix of the heatmap in Figure S3. Primary and PDX samples are hierarchically clustered with complete linkage using Pearson correlation of the scores of 4798 SEs recurrent in at least 2 samples. The heatmap reflects pairwise Pearson correlation of SE scores. The bars at the top reflect presence of coding mutations (grey color designates unknown) and sample type.

**Figure S5, related to** **Figure 1. A SE-based classification of AML: cell lines.** The heatmap is a similarity matrix of the cell lines clustered with complete linkage using Pearson correlation of the scores of 4798 SEs recurrent in at least 2 samples. The heatmap reflects pairwise Pearson correlation of SE scores. The bars at the top reflect presence of coding mutations and sample type.

**Figure S6, related to** **Figure 1. A clique-based classification of AML.** The heatmap represents clique enrichment scores clustered with complete linkage using Pearson correlation. The bars at the top reflect presence of coding mutations and sample type.

**Figure S7, related to** **Figure 1. Selective AML gene dependencies**

(A) MYB as an example of a selective AML dependency. AML cell lines have higher dependency scores compared to non-AML cell lines in the Broad DepMap screens.

(B) A heatmap of 225 selective AML dependencies reflecting probability of dependency inferred from the skewed LRT scores. Rows and columns are hierarchically clustered on Pearson correlation with complete linkage.

(C) SE-associated selective AML dependencies have higher clique scores compared to other SE-associated genes.

(D) Lineage TFs are AML dependencies characterized by high FASE scores (reflecting SE scores and frequency across AML samples). The plot uses 4798 SEs recurrent in at least 2 samples.

(E) Validation of selective AML dependencies identified in the genome-wide screen. MV411 cells stably expressing Cas9 were transduced with GFP-expressing lentiviral vectors encoding gRNAs targeting the indicated genes (2 gRNAs per vector targeting the same gene). Fraction of GFP-positive cells was followed over time.

**Figure S8, related to** **Figure 1. List of CR TFs identified by intersecting selective AML dependencies and common AML SEs.** The table shows gene characteristics, criteria for exclusion/inclusion and data availability (ChIP-seq in unmanipulated MV411 cells and spike-in controlled RNA-seq following TF knockout).

**Figure S9, related to** **Figure 2. Divergence of CR TF-associated SEs in primary AML samples**

(A) A full version of the heatmap in Figure 2D. CR TF-associated SE signatures of epigenetically defined AML clusters. Cluster structure is the same as in Figure 1D. The heatmap visualizes average z-scores of CR TF-associated SEs in each cluster. Note that most CR TFs are associated with multiple distinct SEs.

(B) Differential scores of SEs associated with divergent CRCs by sample type demonstrating similar patterns of CRC divergence.

(C) Validation of RUNX2, MEF2D and IRF8 as selective dependencies of KMT2Ar leukemia using 3 cell lines carrying a KMT2A translocation versus 3 non-KMT2Ar cell lines. The cells were electroporated with *in vitro* assembled Cas9/sgRNA complexes targeting the indicated TF genes, and cell viability was measured relative to an AAVS1 (“safe harbor”) control by quantification of ATP pools using a luciferin-based assay. MYB-targeting sgRNAs were used as a positive control.

**Figure S10, related to** **Figure 2. Classification of primary AML samples based on mRNA expression of CR TFs.** 510 samples from the BeatAML dataset (*27*) were hierarchically clustered using Pearson correlation of mRNA expression of the 31 CR TFs with complete linkage.

(A) A similarity matrix of the 510 BeatAML samples reflecting Pearson correlation between CR TF mRNA expression.

(B) A sample-level hierarchically clustered map of the 510 BeatAML samples visualizing mRNA expression levels of the 31 CR TFs.

**Figure S11, related to** **Figure 3. Dense co-binding of CR TFs at AML SEs.** ChIP-seq binding tracks of 29 CR TFs and several additional proteins (H3K27ac, MED1, BRD4, SMC1, CTCF) in 3 representative loci. 2D plots represent H3K27ac HiChIP connectivity maps with black boxes marking the locations of the ChIP-seq tracks.

**Figure S12, related to** **Figure 3. Co-binding patterns of AML CR TFs.** Analysis of a binary co-binding matrix of 142,970 binding sites created by merging CR TF binding peaks and eliminating sites with only one CR TF bound.

(A) Number of binding sites with *n* CR TFs bound illustrating a progressive decline in the prevalence of binding sites with increasing number of co-binding TFs.

(B) A model of predicted numbers of co-binding sites with *n* TFs binding, assuming a random distribution of TF binding peaks within the co-binding matrix of 142,970 binding sites.

(C) Enrichment of co-binding sites with *n* TFs bound compared to the model that assumes random distribution of TF binding in (B). Sites with 4-10 co-binding CR TFs are negatively enriched (i.e. under-represented compared to a model of random binding distribution), while enrichment of sites with >10 CR TFs binding increases progressively with increasing *n*.

(D) Fraction of TF binding events where an underlying motif was detected.

**Figure S13, related to** **Figure 3. t-SNE analysis of co-binding patterns of AML CR TFs.** A t-SNE plot computed from the binary co-binding matrix of 142,970 binding sites created by merging CR TF binding peaks and eliminating sites with only one CR TF bound. CR TFs are marked in red, while additional proteins, histone marks and regulatory elements are marked in grey.

**Figure S14, related to** **Figure 3. Analysis of co-binding patterns of AML CR TFs using enrichment of pairwise peak overlaps.** The heatmap represents log_2_ fold pairwise enrichments of TFs and chromatin marks computed by Fisher’s exact test on the expected *vs*. observed overlap of peaks in each row with the peaks in each column.

**Figure S15, related to** **Figure 3. Pairwise peak overlaps.** The heatmap represents fractional overlap of row peaks with column peaks. The bottom heatmap represents CR TFs. The top heatmap represents additional proteins and regulatory elements.

**Figure S16, related to** **Figure 4. CRC validation by CRISPR/Cas9 TF knockout followed by mRNA-seq**

(A) Validation of gene knockouts by Western blotting. MV411 cells were electroporated with pre-assembled Cas9/sgRNA complexes targeting the indicated genes and protein depletion was verified by Western blot 72 hours post electroporation. Western blots visualizing MEF2D and MEF2C knockouts are found in Figure 6H.

(B) A t-SNE plot visualizing relationships between CR TF knockouts. The plot was computed from a matrix of log2 fold changes in the expression of top 5000 expressed genes measured by RNA-seq following the indicated CR TF knockouts. Adjacency on the plot illustrates similarity of global transcriptional response.

**Figure S17, related to** **Figures 5**-**6. Application of targeted TF degradation to elucidate direct transcriptional effects of MEF2D and IRF8**

(A) IRF8 dependency correlates with MEF2D dependency in AML cell lines.

(B) Targeted degradation of IRF8 results in loss of cell viability. Cells carrying a fusion-based IRF8 degron were incubated with DMSO *vs.* dTAG^v^-1 and cell viability was followed by quantification of ATP pools using a luciferin-based assay.

(C) Targeted degradation of MEF2D results in loss of cell viability. The experiment was carried out as in (B).

(D) CRISPR/Cas9 knockout of MEF2D results in a loss of cell viability that is quantitatively similar to the effects of MEF2D degradation. MYB knockout was used as positive control.

(E) A combined Pearson similarity matrix of TF knockouts and degron-based experiments hierarchically clustered by correlation between changes in the expression of the top 5000 expressed genes compared to the AAVS1/DMSO controls. The plot shows the expected similarity between degron- and knockout-induced changes. Note that the mRNA-seq was performed 72 hours after TF knockouts vs. 2-24 hours after targeted degradation.

**Figure S18, related to** **Figure 5. Volcano plots of SLAM-seq and mRNA-seq following MEF2D degradation and regulation of MEF2C by MEF2D.** CR TFs are highlighted in blue.

(A) SLAM-seq 2 hours after MEF2D degradation.

(B) SLAM-seq 24 hours after MEF2D degradation.

(C) mRNA-seq 2 hours after MEF2D degradation.

(D) mRNA-seq 24 hours after MEF2D degradation.

(E) Western blot showing reduced MEF2C levels after MEF2D degradation.

**Figure S19, related to** **Figure 5. A table summary of SLAM-seq and RNA-seq data following MEF2D degradation**

(A) Genes demonstrating a statistically significant change in transcription rate (SLAM-seq adjusted *p*-value <0.1) at 2 hours following MEF2D degradation.

(B) Changes in CR TF transcription after MEF2D degradation.

**Figure S20, related to** **Figure 5. A functional interaction between MYC and MEF2D**

(A) Gene set enrichment analysis of the genes showing significant changes in transcription rates after MEF2D degradation using Ehrichr (*126*).

(B) Intersection of gene sets demonstrating significant changes in transcription (adjusted *p*-value <0.05) measured by SLAM-seq after MEF2D degradation and indirect MEF2C/D inhibition by YKL-05-099 treatment.

(C) Excess over bliss synergy matrix demonstrating synergy between SIK/MEF2 inhibitor YKL-05-099 and bromodomain inhibitor JQ1. Red color demonstrates synergy.

(D) Excess over bliss synergy matrix demonstrating synergy between SIK/MEF2 inhibitor YKL-05-099 and MYC inhibitor MYCi361. Red color demonstrates synergy.

(E) Cloning strategy for a doxycycline-inducible lentiviral vector expressing MYC-P2A-TagBFP.

(F) Induction of MYC expression demonstrated by TagBFP fluorescence after addition of doxycycline.

(G) Forced exogenous expression of MYC fails to rescue the cell viability loss following MEF2D degradation.

(H) As a positive control, forced exogenous expression of MYC completely rescues MYC knockout by CRISPR/Cas9. Exogenous MYC expression is induced by doxycycline.

**Figure S21, related to** **Figure 6. Targeted degradation of IRF8**

(A) Volcano plot of SLAM-seq 2 hours after IRF8 degradation.

(B) Volcano plot of SLAM-seq 24 hours after IRF8 degradation. CR TFs are highlighted in blue.

(C) Volcano plot of RNA-seq 2 hours after IRF8 degradation. CR TFs are highlighted in blue.

(D) Volcano plot of RNA-seq 24 hours after IRF8 degradation. CR TFs are highlighted in blue.

(E) A cross-plot of genome-wide changes in mRNA pools measured after 2 *vs.* 24 hours of IRF8 degradation demonstrates a poor correlation between early and late transcriptional response.

(F) A cross-plot of genome-wide changes in mRNA pools (mRNA-seq) *vs.* transcription rates (SLAM-seq) measured after 2 hours of IRF8 degradation.

(G) A distribution plot of genome-wide changes in nascent transcription rates (SLAM-seq) and mRNA pools (RNA-seq) after 2 and 24 hours of IRF8 degradation.

(H) Western blot showing reduced MEF2C and MEF2D levels after IRF8 degradation.

(I) Western blot showing reduced MEF2C and MEF2D levels 24 hours after IRF8 knockout.

(J) Kaplan-Meyer plot of overall survival in TCGA AML patients with MEF2D expression above the 60^th^ percentile (red line) and below the 40^th^ percentile (blue line).

(K) Kaplan-Meyer plot of overall survival in TCGA AML patients with IRF8 expression above the 60^th^ percentile (red line) and below the 40^th^ percentile (blue line).

**Figure S22, related to** **Figure 6. A table summary of SLAM-seq and RNA-seq data following IRF8 degradation.** Genes demonstrating a statistically significant change in transcription rate (SLAM-seq adjusted *p*-value <0.1) at 2 hours following IRF8 degradation are shown.

## METHODS

### EXPERIMENTAL MODEL AND SUBJECT DETAILS

#### Cell lines and patient-derived xenograft samples

AML cell lines were cultured in the RPMI-1640 media containing 10% fetal bovine serum and regularly tested to be free of *Mycoplasma spp*. Patient-derived xenograft samples (PDX) were obtained from the DFCI PRoXe repository (https://proxe.shinyapps.io/PRoXe/). PDX cytogenetics and molecular characteristics were obtained from the PRoXe database (*40*).

#### Primary AML samples

Samples from children with AML were obtained with informed consent, according to protocols approved by the Institutional Review Board of Baylor College of Medicine. Patients’ cytogenetics and molecular characteristics were collated from clinical chart records. At the time of collection, mononuclear cells were enriched by density centrifugation and cryogenically preserved. Samples selected for inclusion contained high purity of blasts (>85%). Cells were preserved in conditioned media from the human bone marrow stromal cell line HS5 (collected after 2 days of plating) diluted 1:1 with RPMI+10% FBS and 1% Penicillin/Streptomycin. Freshly thawed cells were cross-linked with formaldehyde and used for ChIP-seq.

### METHOD DETAILS

#### CRISPR/Cas9 gene knockouts by RNP electroporation

Synthetic modified sgRNA constructs were purchased from Synthego (Redwood City, CA). Ribonucleoprotein (RNP) assembly was performed by mixing 2-3 sgRNAs (a total of 120 pmol) with 8.5 μg recombinant Cas9 (Invitrogen A36499). The resulting RNP mix was electroporated into 0.3×10^6^ MV411 cells using a Lonza 4D Nucleofector, program DJ-100, in 20 μl Nucleocuvette strips (Lonza V4XC-2032). Unless otherwise noted, cells were incubated in media for 72 hours post-electroporation before subsequent analyses. Knockout efficiency was confirmed by Western blotting and PCR amplification followed by indel analysis. A guide RNA targeting the AAVS1 “safe harbor” locus was used as a negative control (*127*). Cell viability was measured at indicated times post-electroporation using the CellTiter-Glo luminescent cell viability assay (Promega G7570).

#### CRISPR/Cas9 gene knockouts by lentiviral transduction

We used lentiviral delivery of gRNAs for initial validation of 11 arbitrarily chosen AML dependencies (Figure S7E). Lentiviral vectors encoding gRNAs (bicistronic vectors with 2 gRNAs per vector: pLV[2gRNA]-EGFP:T2A:Puro-U6>[hgRNA#1]-U6>[gRNA#2]) were purchased from VectorBuilder (Cyagen Biosciences, Santa Clara, CA) and packaged into lentivirus. Cas9-expressing AML cell lines were transduced by spinfection as described (*128*). The guide dropout was measured every 2 days for a total of 22 days as the percentage of GFP+ cells and normalized to the percentage of GFP+ cells 48 hours transduction.

#### Chromatin immunoprecipitation sequencing (ChIP-seq)

ChIP-seq for hematopoietic TFs was performed using 100×10^6^ exponentially growing MV411 cells per experiment. For histone ChIP-seq, 2.5×10^6^ cells were used. Cells were fixed with 1% formaldehyde for 10 minutes at room temperature, quenched with 125 mM glycine for 5 minutes, and washed 3 times with PBS. Nuclei were isolated using Nuclei EZ Isolation buffer (Sigma NUC-101) and resuspended in 10 mM Tris-HCl, pH 8.0, 1 mM EDTA, 0.1% SDS with 1x HALT protease inhibitor (ThermoFisher 78430). Chromatin was fragmented by sonication on an E220 Covaris focused sonication machine using 1 ml glass AFA tubes (Covaris 520135) and the following parameters: 140 mV, 5% duty factor, 200 cycles/burst for 14 minutes. The rest of the ChIP-seq protocol was performed as previously described (*23*) and in accordance with the Encode guidelines (*129*). ChIPseq libraries were prepared using Swift S2 Acel reagents (Swift 21096) on a Beckman Coulter Biomek i7 liquid handling platform from approximately 1 ng of DNA according to manufacturer’s protocol and using 14 cycles of PCR amplification. Sequencing libraries were quantified by Qubit fluorometer and Agilent TapeStation 2200. Library pooling and indexing was evaluated by shallow sequencing on Illumina MiSeq. Subsequently, libraries were sequenced on Illumina NextSeq 500 or NovaSeq 6000 by the Molecular Biology Core facilities at the Dana-Farber Cancer Institute.

For quantitative ChIP-seq analysis of H3K27 acetylation and TF binding we used *Drosophila* chromatin/antibody spike-in control as previously described (*130*). For H3K27ac ChIP-seq, 4 μg of anti-H3K27ac antibody, 2 μg of spike-in antibody and 20 ng of spike-in chromatin (Active Motif 61686 and 53083, respectively) were added to chromatin prepared from 2.5×10^6^ MV411 cells 72 hours after RNP-mediated TF knockout. For TF ChIP-seq, 10 μg of anti-TF antibody, 5 μg of spike-in antibody and 50 ng of spike-in chromatin were added to chromatin prepared from 100×10^6^ MV411 cells. The rest of the ChIP-seq experiment was performed in the standard fashion. After ChIP-seq reads were mapped to the *Drosophila* genome and the hg38 human genome in parallel, human tag counts were normalized to *Drosophila* tag counts.

#### HiChIP

HiChIP with primary antibodies against H3K27ac (Abcam ab4729) was performed as described (*41, 131*). MV411 cells (3 replicates of 30×10^6^ cells each) were cross-linked with 1% formaldehyde in PBS at room temperature for 10 minutes, quenched with 125 mM glycine for 5 minutes, washed with PBS 3 times and flash-frozen. Cross-linked cell pellets were thawed on ice and resuspended in 10 μL ice-cold Hi-C lysis buffer (10 mM Tris-HCl pH 8.0, 10 mM NaCl, 0.2% IGEPAL CA-630, 1X cOmplete Protease Inhibitor Cocktail (Roche 11697498001)) and incubated for 30 minutes at 4°C rotating. Nuclei were pelleted by centrifugation at 2500 x g for 5 minutes at 4°C, discarding supernatant, and washed with 500 μL ice-cold Hi-C lysis buffer. Nuclei were resuspended in 100 μL 0.5% SDS and incubated at 62°C for 10 minutes, then quenched by addition of 335 μL 1.5% Triton X-100 and incubation at 37°C for 15 minutes. To digest chromatin, 50 μL NEB Buffer 2 and 375 U MboI (NEB R0147M) were added and allowed to incubate for 2 hours at 37°C rotating, before heat inactivating at 62°C for 20 minutes. Sticky ends were filled-in by adding 37.5 uL 0.4 mM biotin dATP (Invitrogen 19524016), 1.5 μL 10 mM dCTP, 1.5 μL 10 mM dGTP, 1.5 μL 10 mM dTTP, and 10 μL 5 U/μL DNA Polymerase I, Large (Klenow) Fragment (NEB M0210L), and incubating for 1 hour at 37°C rotating. Proximity ligation was performed by adding 150 μL 10X NEB T4 Ligase Buffer, 125 μL 10% Triton X-100, 7.5 μL 20 mg/mL BSA, 10 μL 400 U/μL T4 DNA Ligase (NEB M0202L), and 655.5 μL water, and incubating for 4 hours at room temperature rotating. Proximity ligated chromatin was sheared by sonication using a Covaris S220 sonicator in 1 mL Covaris milliTube using settings: fill level 10, duty cycle 5, peak intensity power 140, cycles per burst 200, time 6 minutes. Sonicated chromatin was clarified by centrifugation at 16,100 x g for 15 minutes at 4°C, discarding the pellet, and pre-cleared by incubation with 60 μL Dynabeads Protein G (Invitrogen 10004D) for 1 hour at 4°C rotating. Chromatin immunoprecipitation was performed by incubating the pre-cleared chromatin with 75 μL Dynabeads Protein G bound with 7.5 μg H3K27ac antibody (Abcam ab4729) overnight at 4°C rotating. The beads were then washed twice with 1 mL sonication buffer (50 mM HEPES-KOH pH 7.5, 140 mM NaCl, 1 mM EDTA pH 8.0, 1 mM EGTA pH 8.0, 1% Triton X-100, 0.1% sodium deoxycholate, 0.1% SDS), once with 1 mL high salt sonication buffer (50 mM HEPES-KOH pH 7.5, 500 mM NaCl, 1 mM EDTA pH 8.0, 1 mM EGTA pH 8.0, 1% Triton X-100, 0.1% sodium deoxycholate, 0.1% SDS), once with 1 mL LiCl wash buffer (20 mM Tris-HCl pH 8.0, 1 mM EDTA pH 8.0, 250 mM LiCl, 0.5% IGEPAL CA-630, 0.5% sodium deoxycholate, 0.1% SDS), and once with 1 mL 50 mM NaCl in TE buffer. Immunoprecipitated chromatin was eluted by resuspending the beads in ChIP elution buffer (50 mM Tris-HCl pH 8.0, 10 mM EDTA pH 8.0, 1% SDS) and incubating at 65°C for 15 minutes. Eluted chromatin was treated with 2.5 uL 33 mg/mL RNase A (Sigma R4642) for 2 hours at 37°C followed by 10 uL 20 mg/mL proteinase K (Invitrogen 25530049) for 45 minutes at 55°C, then incubated at 65°C for 5 hours to reverse cross-links. DNA was purified using Zymo ChIP DNA Clean and Concentrator Kit (Zymo D5205) and eluting with 14 μL water. 5 μL Dynabead MyOne Streptavidin C1 beads (Invitrogen 65001) were washed with 1 mL Tween wash buffer (5 mM Tris-HCl pH 7.5, 0.5 mM EDTA pH 8.0, 1M NaCl, 0.05% Tween-20) and resuspended in 10 μL 2X Biotin binding buffer (10 mM Tris-HCl pH 7.5, 1 mM EDTA pH 8.0, 2M NaCl). To capture biotinylated DNA, 10 μL purified DNA was added to the beads and incubated for 15 minutes at room temperature with intermittent agitation, discarding supernatant. Beads were washed twice with 500 μL Tween wash buffer at 55°C for 2 minutes shaking at 1000 rpm. For tagmentation, beads were resuspended in 25 μL Nextera Tagment DNA Buffer (2X) (Illumina FC-121-1030), 3.25 μL TDE1 per 50 ng DNA, and resuspension buffer (RSB) up to 50 μL final volume; and allowed to incubate at 55°C for 2 minutes shaking at 1000 rpm. The reaction was quenched by adding 500 μL 50 mM EDTA and incubating at 50°C for 30 minutes. Beads were then washed twice with 500 μL 50 mM EDTA incubating at 50°C for 3 minutes, twice with 500 μL Tween wash buffer incubating at 50°C for 2 minutes, and once with 500 μL 10 mM Tris-HCl pH 7.5. To prepare HiChIP libraries, beads were resuspended in 15 μL Nextera PCR Master Mix, 5 μL PCR Primer Cocktail, 5 μL index primer 1, 5 μL index primer 2, and 20 μL water. Hi-ChIP libraries were amplified by PCR program: 72°C for 5 minutes, 98°C for 1 minute, [98°C for 15 seconds, 63°C for 30 seconds, 72°C for 1 minute, repeated for 10 cycles], 72°C for 1 minute, 4°C hold. HiChIP libraries were purified using the Zymo DNA Clean and Concentrator kit (Zymo D4013) and eluted using two volumes of 10 μL water each time. Purified HiChIP libraries were subjected to 50 bp paired-end sequencing on an Illumina HiSeq 2500.

#### RNA-seq

For RNA-seq experiments the total cellular RNA was extracted using the QuickRNA kit (Zymo Research R1054). Purified total RNA was mixed with the ERCC ExFold RNA Spike-in mix (Invitrogen 4456740). RNA sequencing libraries were prepared on a Beckman Coulter Biomek i7 liquid handling platform using Roche Kapa mRNA HyperPrep strand specific sample preparation kits (Roche 08098123702) from 200 ng of purified total RNA according to the manufacturer’s protocol. Library quantification and Illumina sequencing were performed as described in the ChIP-seq section above.

#### Western blotting

Whole-cell lysates were prepared in RIPA buffer (Boston Bio-Products BP-115-500) with protease inhibitor cocktail (ThermoFisher 23225). Lysates were boiled in Laemmli buffer (BioRad 1610737), separated by SDS-PAGE, and transferred and blocked using standard methodology. HRP-conjugated anti-mouse and anti-rabbit IgG secondary antibodies were used for imaging (BioRad 1706515 and 1706515) with an enhanced chemiluminescence substrate (PerkinElmer NEL104001EA) according to manufacturers’ instructions.

#### Targeted TF degradation

MV411 cells were modified by CRISPR-HDR to express N-terminal FKBP12 fusions of IRF8 and MEF2D, respectively. For each knock-in, donor DNA constructs were chemically synthesized and cloned into the pUC19 or pAAV-MCS2 plasmids obtained from Addgene (Watertown, MA). The donors included 400-800 bp homology arms flanking the inserted DNA sequence encoding the FKBP12 degradation tag as well as mScarlet, FLAGx3 and HA tags. MV411 cells were electroporated with Cas9/sgRNA complexes targeting the HDR insertion site (with sgRNA protospacer sequence spanning the insertion site). Electroporation was performed using Lonza SF Cell Line 4D Nucleofector (V4XC-2032). RNP complexes were formed by mixing 8.5 μg of TrueCut Cas9 Protein v2 (Invitrogen A36499) and 120 pmol sgRNA. 0.3×10^6^ cells were washed with PBS and resuspended in 20 μL of SF Cell Line solution (Lonza). The cells were combined with the RNP mix and 0.6 μg pDNA donor and electroporated using program DJ-100. After a 5-7 day incubation period the cells were sorted for mScarlet fluorescence. Single clones were then obtained by single-cell dilution microwell plating and screened for bi-allelic donor insertion by PCR. Clones were validated by Western blotting and Sanger sequencing. TF degradation was induced by adding 500 nM of dTAG^v^-1 as previously described (*80*) and followed by FACS measurement of mScarlet fluorescence and Western blotting.

#### SLAM-seq

Thiol (SH)-linked alkylation for the metabolic sequencing of RNA (SLAM-seq) was performed as described (*82*). Briefly, a total of 2.5×10^6^ MV411 cells per replicate were incubated with 500 nM dTAG^V^-1 for 2 or 24 hours. S^4^U labeling was performed by adding S^4^U to a final concentration of 100 μM for an additional hour. Cells were flash-frozen and total RNA was extracted using Quick-RNA MiniPrep (ZYMO Research) according to the manufacturer’s instructions except including 0.1 mM DTT to all buffers. ERCC ExFold RNA Spike-in mix (Invitrogen) was added to purified RNA. Thiol modification was performed by 10 mM iodoacetamide treatment followed by quenching with 20 mM DTT. RNA was purified by ethanol precipitation and mRNA-seq was performed as described above.

#### Immunofluorescence with RNA FISH

Glass coverslips were coated at 37°C with 0.01% of poly-L-lysine solution (Sigma-Aldrich, P4707) for 30 minutes. Cells were plated on the pre-coated coverslips and grown for 24 hours followed by fixation using 3.7% paraformaldehyde (VWR, BT140770) in PBS for 10 minutes. Cells were washed with PBS twice followed by permeabilization using 70% ethanol overnight. Cells were washed with Wash buffer A (20% Stellaris RNA FISH Wash Buffer A (Biosearch Technologies, Inc., SMF-WA1-60), 10% Deionized Formamide (EMD Millipore, S4117) in RNase-free water (Life Technologies, AM9932) for 3 minutes. Coverslips were incubated in Hybridization buffer (90% Stellaris RNA FISH Hybridization Buffer (Biosearch Technologies, SMF-HB1-10) and 10% Deionized Formamide) containing diluted primary antibodies (see table x) and 25 mM RNA probe (Human MYC_intron with Quasar® 570 Dye, ISMF-2066-5, Stellaris) for 4.5 hours at 37°C in a humidified chamber. Cells were then incubated in anti-rabbit 488 antibody (A-11008, Invitrogen) at a concentration of 1:100 in buffer A for 30 minutes at 37°C and then in Buffer A containing the same secondary antibody plus 20 mm/mL Hoechst 33258 (Life Technologies, H3569) for an additional 30 minutes at 37°C, followed by a 5-minute wash in Wash buffer B (Biosearch Technologies, SMF-WB1-20). The coverslips were then mounted onto glass slides with Vectashield (VWR, 101098-042) and sealed with nail polish (Electron Microscopy Science Nm, 72180). Immunofluorescence without RNA FISH was done using the same protocol except a mouse primary antibody against MEF2C was added instead of the RNA FISH probe, and an anti-mouse secondary antibody was used in addition to the anti-rabbit secondary antibody.

Images were acquired at the Harvard Center for Biological Imaging (HCBI) using lattice-based structured illumination microscopy (lattice SIM) on an ELYRA 7 super-resolution microscope (Carl Zeiss Microscopy, Jena, Germany), using a 63x/1.4 objective and imaging on pco.edge 4.2 sCMOS cameras (a dual camera setup with motorized precision alignment). Image acquisition, post-processing and primary analysis was conducted with Carl Zeiss Microscopy ZEN software. Images were post-processed using standard SIM settings with automatic channel alignment and subsequent Maximum Intensity Projection.

#### Exogenous MYC expression

For exogenous MYC expression a synthetic DNA sequence encoding MYC fused to TagBFP via a P2A linker was cloned into the pLVX-TetOne-Puro vector (Takara Bio 631849) and the construct was packaged into a lentiviral vector. MV411 cells were transduced with the lentivirus and selected with 1 μg/mL puromycin. RNP-mediated knockouts were carried out as described above. MYC expression was induced by adding 1 μg/mL doxycycline immediately after RNP electroporation and verified by Western blot (not shown) and TagBFP fluorescence. Cell viability was followed by the CellTiter-Glo luminescent cell viability assay (Promega G7570).

#### Direct inhibition of enhancer activity

For direct inhibition of enhancer activity we used the dCas9–KRAB–MeCP2 system (*43*). sgRNAs targeting predicted enhancers in the vicinity of *MYB* and *IRF8* genes were cloned in a lentiviral expression vector (LRG2.1_Puro) and packaged into lentivirus. MV411 cells were first transduced with a lentiviral vector expressing dCas9–KRAB–MeCP2 and selected by incubation with 10 μg/mL blasticidin. The dCas9–KRAB–MeCP2-expressing cells were transduced with lentiviral gRNA-expressing vectors and additionally selected by incubation with 2 μg/mL puromycin for 48 hours. Expression of the target genes (*MYB* and *IRF8*) was measured by ΔΔCt TaqMan qPCR and normalized to β-actin.

### EXTERNAL DATASETS

RNA-seq BAM files for the BeatAML project (*27*) were provided by Oregon Health & Science University and processed through the CCLE RNA processing pipeline (STAR/RSEM, described at https://github.com/broadinstitute/ccle_processing). Reads were normalized to transcripts per million (TPM) and filtered for protein coding genes. The expression values were transformed to log2(TPM+1).

H3K27ac ChIP-seq data from primary AML samples (*24*) were downloaded from Sequence Read Archive (SRA) under accession number SRP103200 and processed using the AQUAS pipeline (https://github.com/kundajelab/chipseq_pipeline) with minor modifications and according to the ENCODE3 guidelines. We used data from 49 samples that passed our quality criteria.

Genetic dependency data are available for download at the Broad DepMap portal database (https://depmap.org/portal/download/). Data release 20q1 was used for this study.

ATAC-seq data in MV411 cells were downloaded from https://www.ncbi.nlm.nih.gov/sra?term=SRX5608489.

The following external ChIP-seq datasets were downloaded from SRA: SRR5054413, SRR5054417, SRR5054418, SRR5054416, SRR5054423, SRR3617110, SRR4423332, SRR4423334, SRR4423333, SRR7266894, SRR7266893, SRR7266892, SRR7266885, SRR7266884, SRR7266883, SRR7266874, SRR7266875, SRR7266876, SRR5863449, SRR5863448, SRR5863451, SRR5863452, SRR2532559, SRR2532562, SRR2532564, SRR3465915, SRR3465916, SRR3331425, SRR3331423.

### QUANTIFICATION AND STATISTICAL ANALYSIS DETAILS

#### ChIP-seq data analysis

Quality control, mapping and analysis of the ChIP-seq data was performed using the nf-core pipeline (https://github.com/nf-core/chipseq). Differential binding of the same protein under two conditions was computed using the *diffpeak* function of the MACS2 pipeline (https://github.com/macs3-project/MACS). Spike-in controlled experiments were mapped to the *Drosophila* genome and the hg38 human genome in parallel, and human tag counts were normalized to *Drosophila* tag counts as described (*67, 130*).

#### HiChIP data analysis

HiChIP datasets were processed using HiC-Pro (https://github.com/nservant/HiC-Pro) with default settings. Briefly, HiChIP paired-end reads were aligned using Bowtie2 with the following parameters*: --very-sensitive - L 30 --score-min L, -0.6, -0.2 --end-to-end --reorder*. The restriction sites were obtained by scanning for MboI restriction enzyme fragments across the human genome. Valid interactions pairs (*validPairs*) were converted to a .hic file using the hicpro2juicebox.sh script from the utility tool of HiC-Pro. The generated .hic file contains interaction matrices at fragment and base pair resolutions. Interaction maps were visualized with Juicebox (https://aidenlab.org/juicebox/).

#### SE calling and gene assignment

SE calling and merging was performed essentially as described (*132*). Briefly, H3K27ac ChIP-seq reads were aligned to hg19 genome using BWA-ALN. Duplicate reads were removed using Picard MarkDuplicates (https://broadinstitute.github.io/picard/) and samtools (http://www.htslib.org). Fragment length was estimated using the spp package (https://cran.r-project.org/web/packages/spp/index.html). Broad peaks were called using MACS2 with a *p*-value cutoff of 1×10^-5^ using the estimated fragment length. For SE calling each sample was run through ROSE2 (https://github.com/linlabbcm/rose2) excluding 2500 bp around TSSs (-t 2500) and the hg19 encode blacklisted regions. SE regions were then merged and ROSE2 was re-run on all sample using the merged regions, producing the signal matrix, which was then normalized by median signal. Two or more replicate ChIP-seq experiments were performed for the vast majority of samples, and SE scores were averaged between the replicates. SE coordinates were then lifted to hg38 using the USCE Genome Browser LiftOver tool (http://genome.ucsc.edu/cgi-bin/hgLiftOver) and all subsequent analyses were conducted using the hg38 genome assembly.

To assign SEs to genes we used a modified activity-by-contact (ABC) procedure (*42*). First, we identified H3K27ac HiChIP loops with one end within 5000 bp of the transcription start site. Separately, we aggregated H3K27ac ChIP-seq and ATAC-seq activity within 2500 bp on either side of the non-TSS end of the HiC loop. We then calculated the ABC strength score as a product of the HiChIP loop frequency and the geometric mean of H3K27ac ChIP-seq and ATAC-seq activities. This procedure yielded ca. 2.71 million loops, each associated with a particular gene by having one end near a TSS. We then trimmed the ABC loop set to remove (a) loops with both ends within 5000 bp of the TSS (accounting for ∼60,000 loops), (b) loops that overlapped with blacklist areas (as identified in https://github.com/Boyle-Lab/Blacklist/tree/master/lists/hg38-blacklist.v2.bed.gz, accounting for ∼10,000 loops), and (c) loops with ABC scores below the 89th percentile genome-wide (∼ 2,350,000 additional loops). Adjacent non-TSS regions were then stitched together. This resulted in 238,220 ABC regions. Subsequently, H3K27ac ChIP-seq peaks for all samples were re-mapped to the 238,220 ABC-defined regions using bamliquidator (https://github.com/BradnerLab/pipeline/wiki/bamliquidator). For each region, we extracted H3K27ac area under the curve in each sample and normalized by total H3K27ac across all the regions in that sample to yield a normalized enhancer signal. We then calculated the Pearson correlation coefficient between (1) normalized enhancer signals and (2) mRNA expression (in log2[TPM+1]) for each ABC region, separately across the PDX and cell line sample sets. Effective ABC region/gene associations were chosen when either correlation coefficient was > 0.3, resulting in a total of 23170 associations. The ABC associations were merged with the standard proximity associations computed by ROSE2. Overall, our algorithm yielded 6868 distinct SEs assigned to 11866 genes with an average of 3.7 gene associations per SE.

#### Identification of selective AML dependencies

Identification of selective AML dependencies was performed essentially as described (*45*). Data from genome-scale CRISPR-Cas9 loss-of-function screens of 74,378 guide RNA species targeting 18,333 human genes in 769 cell lines (including 20 AML cell lines) were downloaded from the Broad DepMap portal database (https://depmap.org/portal/download/). Data release 20q1 was used for this study (https://figshare.com/articles/dataset/DepMap_20Q1_Public/11791698). Gene dependency scores were calculated as previously described (*52*). Briefly, the abundance of each guide RNA at the time of infection to its abundance after 21 days of cell culture was compared and aggregated into a single score per gene. The relative drop out ratio of each gene was then normalized to negative controls (score = 0, representing non-essential genes) and positive controls (score = −1, reflecting the median score of common essential genes). For each gene the dependency probability was estimated as the likelihood that the gene represented a phenotype similar to positive controls (*51*) (Figure S7A). The common essential genes were identified as those genes that were ranked in 90% of cell lines above a cutoff determined from the central minimum in the histogram of gene ranks in their 90th percentile least dependent line as previously described (*51*).

To identify genetic dependencies that had *skewed* distributions across the cell lines screened, normLRT scores were calculated (*54*). The log likelihood ratio of fitting to a skewed distribution for the dependency scores of each gene with the skew-t parametric family of skew-elliptically contoured distribution for the error term. The log likelihood ratio of fitting to a normal distribution was calculated for the dependency scores of each gene. The normLRT score is twice the difference of the log of the likelihood ratio of fitting to a skewed distribution and the log of the likelihood ratio of fitting to a normal distribution. *Skewed* gene dependencies were defined as those with normLRT scores greater than or equal to 100 and left-sided skew, as indicated by a mean gene effect score less than the median gene effect score.

To identify *enriched* dependencies in AML, a two-class comparison was performed between the gene effect scores for AML cell lines (in-group, n=20) and the remainder of all other cell lines in the screen (out-group, n=749) as previously described (Dharia et al., 2021). Briefly, a linear model was fitted to the gene effect scores divided in the in-group and out-group. Next, t-statistics and log-odds ratios of differential gene effect were computed. Effect size was calculated as the difference in the mean gene effect dependency score in the in-group compared to that in the out-group. In addition to two-sided P values, one-sided ‘left’ P values were calculated to identify gene dependency effects that were more negative (more dependent) in the in-group than in the out-group, and one-sided ‘right’ P values were calculated to identify those that were less dependent in the in-group than in the out-group. All P values were corrected for multiple hypothesis testing using the Benjamini–Hochberg correction, and these adjusted P values were reported as q values. AML *enriched* genetic dependencies were identified as those with a q value less than 0.05 with a negative effect size (the mean of dependency gene effect score was more negative in the in-group than in the out-group).

We then used this dataset to define a list of 225 selective AML dependencies by identifying genes which met all of the following criteria (Figure S7B): 1) probability of dependency >0.5 in 3 or more AML cell lines; 2) not classified as a common essential gene in the screen, and 3) either classified as an *enriched* dependency OR a *skewed* dependency.

#### Merging of TF ChIP-seq replicates

We developed CREME, a fast post-processing ChIP-seq replicate merging algorithm that does not need any raw sequencing data. CREME uses as input sets of BED files and bigWig tracks for each protein, and outputs a merged BED file, as well as merging quality metrics. Given a set of replicates, CREME first computes a consensus set of peaks by taking the union of all of their peaks and considering any peak ≤150 bp away from another to be in overlap. The overlapping peaks are then merged and assigned the mean of their signals and the product of their p-values. CREME then computes a similarity score as follows:

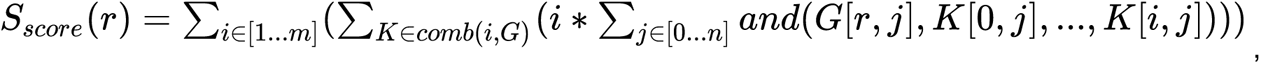

where:

*G* is an *m*\**n* binary matrix of *m* replicates with *n* consensus peaks and a value of 1 if replicate *m_i_* has a peak on consensus peak *m_i_* and 0 otherwise,

*r* is one of the replicates,

*combi(i, G)* is a list of all possible matrices made from taking *i* replicates from matrix *G* without replacement,

*and()* is a binary operation returning 1 if all passed elements are 1 else 0.

The highest scoring sample not labelled as a bad-quality replicate will be selected as the main replicate (replicate A). Bad-quality replicates are user provided annotations, *e.g.* found by visual inspection of bigWig tracks, a threshold on FRiP scores or any other QC method. If the second best scoring replicate (replicate B) and the main replicate have less than 30% peak overlap CREME marks that ChIP as failed and only returns the main replicate.

Then, for each replicate, CREME searches for new peaks using a modified MACS2 peak calling algorithm with a lower discovery threshold (a KL distance of 8). We assume that the peaks are ‘events’ in a Poisson process occurring across the genome and compute the difference between the region of interest and its entire chromosome, using the KL divergence between two poisson distributions originating from these two regions. The signal for these events is extracted from the bigWig files. If the KL distance is above the threshold we consider the region as a peak. Using this approach, CREME will attempt to call peaks in replicate B using peak loci that were called in replicate A and do not overlap with replicate B. The same is then repeated in reverse, attempting to call peaks in replicate A using non-overlapping peak regions from replicate B. If, despite the additional peak calling, we still detect less than 30% overlap between replicates, replicate B is discarded. Otherwise, the newly called peaks are added to the merged peak set. CREME then repeats this procedure with additional replicates ranked in the order of their quality. The final merged BED file contains all consensus peaks of the main replicates (including its newly found peaks) and any other peaks present in at least 2 replicates.

#### RNA-seq data analysis

RNA-seq data were processed using the CCLE workflow of STAR (v2.6.1c) + RSEM (v1.3.0) with hg38 reference genome, enriched with ERCC92 v29 reference (https://storage.googleapis.com/ccle_default_params/Homo_sapiens_assembly38_ERCC92.fasta; https://storage.googleapis.com/ccle_default_params/STAR_genome_GRCh38_noALT_noHLA_noDecoy_ERCC_v29_oh100.tar.gz; https://storage.googleapis.com/ccle_default_params/rsem_reference_GRCh38_gencode29_ercc.tar.gz) and Gencode v29 reference gene regions (https://storage.googleapis.com/ccle_default_params/references_gtex_gencode.v29.GRCh38.ERCC.genes.collapsed_only.gtf). Scaling factors were computed from ERCC spike-ins using the ERCCdashboard R package (https://bioconductor.org/packages/release/bioc/html/erccdashboard.html). Scaling was applied only to samples where ERCC displayed a mean scaling of at least twice the size of standard error. Differential analysis of RNA-seq data was performed using DESeq2 (v1.26.0). (https://bioconductor.org/packages/release/bioc/html/DESeq2.html), using the ERCC pseudogenes to rescale the data by using *the run_estimate_size_factors control_genes* parameter.

#### SLAM-seq data analysis

A modified version of the slamdunk pipeline was used for SLAM-seq processing. Differential analysis of SLAM-seq data was performed using DESeq2 on the TC converted transcripts (*tccounts*) and total read counts (*totalcounts*). First, mean *totalcounts* were used to compute scaling factors via the DESeq2 *run_estimate_size_factors geoMeans* parameter. Then, the *tccounts* and *totalcounts* were normalized to the ERCC pseudogene counts using DEseq2 *getSizeFactors* and *setSizeFactors* functions.

#### Motif analysis

Motif analysis was done using the MEME (v5.1.0) suite with its HOCOMOCOv11_full_HUMAN_mono database of human motifs (https://meme-suite.org/meme/db/motifs). We used a custom genome build of MV411 cells computed on the hg38 human genome with mutations called by GATK4’s Haplotype Caller (https://dockstore.org/workflows/github.com/broadinstitute/depmap_omics/cnn-variant-filter:master) from the CCLE 30x whole genome sequencing dataset (https://www.ncbi.nlm.nih.gov/sra/SRX5449767[accn]). Motifs were called using open region of the genome using publicly available MV411 ATAC-seq data (SRA SRX5608489). Found motifs were merged into the consensus set of peaks (the co-binding matrix above), using genepy *simpleMergePeaks* function, merging any motifs <100 bp away from a peak’s center.

#### SIM microscopy

The ZEN machine learning package was used for image processing (https://www.zeiss.com/microscopy/int/products/microscope-software/zen.html#downloads). We used ZEN 3.0 Black for image processing, including channel alignment, SIM processing and image subsetting, and ZEN 3.1 Blue for data segmentation and formatting. The segmentation (puncta recognition) algorithm was trained on a subset of raw images labelled manually.

First, 2D (discoid) puncta of individual z-stacks were aggregated into 3D spheroids if the distance between the centers of the discoid puncta were less than the size of an average discoid. Puncta present only in one z-stack were discarded. In the rare cases when >3 RNA FISH foci were detected per nucleus due to background noise, only the top 3 in terms of total fluorescence were counted, as MV411 cells carry 3 copies of the MYC gene.

The co-localization enrichment was computed by Fisher’s exact test between the expected number of red (RNA FISH) puncta co-localizing with the green (protein IF) *versus* the actual measured number. The expected (random) distribution of puncta was computed by defining the total volume of the nucleus occupied by the green puncta.

