## Supplemental Figures S1-S22 for "A distinct core regulatory module enforces oncogene expression in KMT2A-rearranged leukemia"

### Figure S1

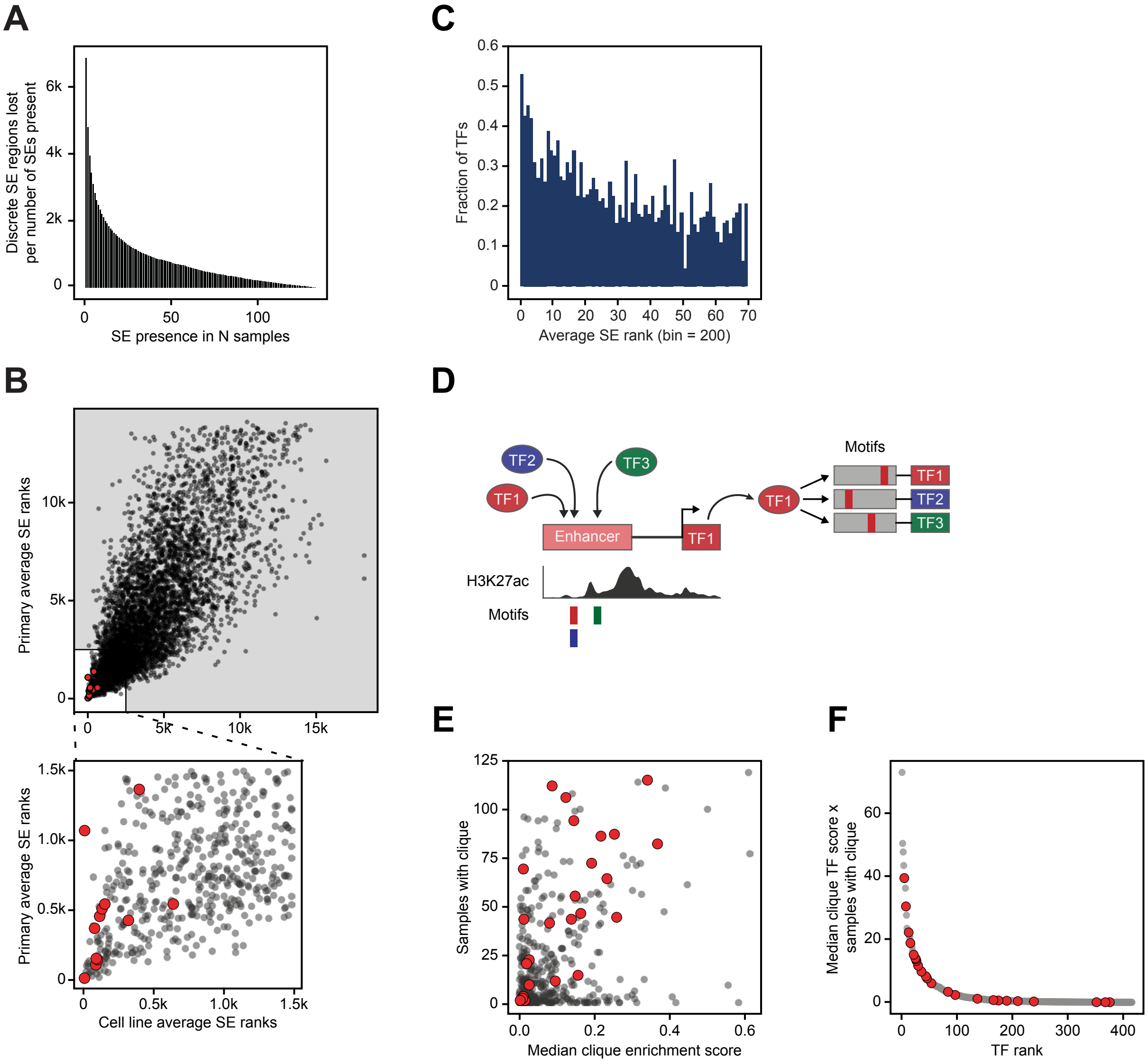
A
Discrete SE regions lost per number of SEs present
SE presence in N samples
C
Fraction of TFs
Average SE rank (bin = 200)
B
Primary average SE ranks
Cell line average SE ranks
D
TF2
TF3
TF1
Enhancer
Motifs
H3K27ac
E
Samples with clique
Median clique enrichment score
F
Median clique TF score x samples with clique
TF rank

### Figure S2

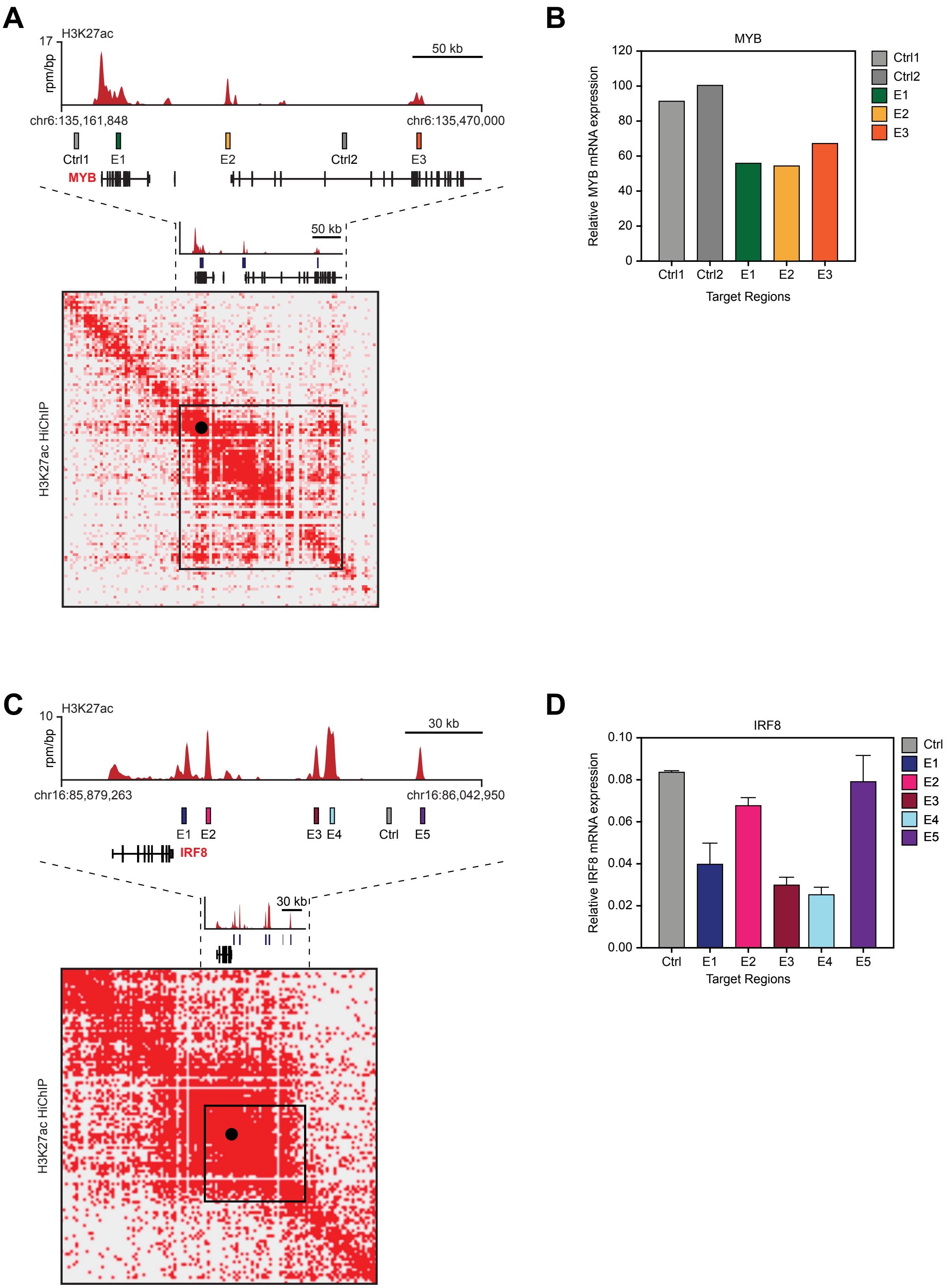
A
H3K27ac
17
rpm/bp
50 kb
chr6:135,161,848
chr6:135,470,000
Ctrl1
E1
E2
Ctrl2
E3
MYB
50 kb
H3K27ac HiChIP
B
MYB
Relative MYB mRNA expression
120
100
80
60
40
20
0
Ctrl1
Ctrl2
E1
E2
E3
Target Regions
C
H3K27ac
10
rpm/bp
30 kb
chr16:85,879,263
chr16:86,042,950
E1
E2
E3
E4
Ctrl
E5
IRF8
30 kb
H3K27ac HiChIP
D
IRF8
Relative IRF8 mRNA expression
0.10
0.08
0.06
0.04
0.02
0.00
Ctrl
E1
E2
E3
E4
E5
Target Regions

### Figure S3

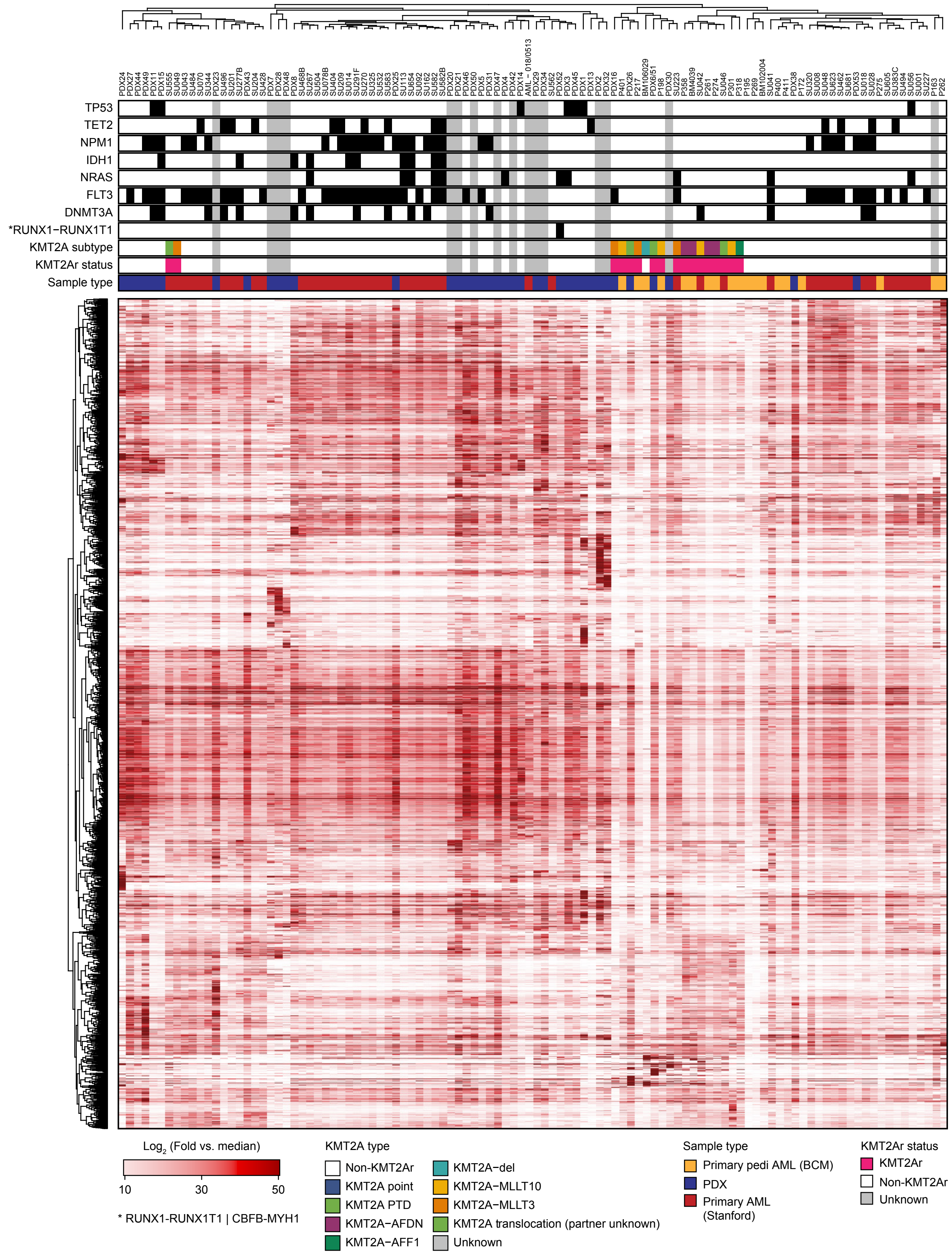

PDX24 PDX27 PDX44 PDX49 PDX11 PDX15 SU555 SU049 SU043 SU484 SU070 SU344 PDX23 SU496 SU201 SU277B PDX43 SU204 SU428 PDX7 PDX28 PDX48 PDX8 SU468B SU267 SU504 SU078B SU404 SU209 SU014 SU291F SU270 SU325 SU532 SU583 PDX25 SU113 SU654 SU092 SU162 SU582B PDX20 PDX21 PDX46 PDX50 PDX5 PDX31 PDX47 PDX4 PDX42 PDX14 AML - 018/0513 PDX29 PDX34 SU562 PDX52 PDX3 PDX45 PDX1 PDX13 PDX2 PDX32 PDX16 P401 PDX26 P217 BM106029 PDX6/51 P198 PDX30 SU223 P358 BM4039 SU042 P261 P274 SU046 P301 P318 P195 P269 BM102004 SU041 P400 P411 PDX38 P172 SU320 SU008 SU048 SU623 SU462 SU681 PDX53 SU018 SU028 P275 SU605 SU383C SU494 SU056 SU001 SU227 P163 P262
TP53
TET2
NPM1
IDH1
NRAS
FLT3
DNMT3A
*RUNX1−RUNX1T1
KMT2A subtype
KMT2Ar status
Sample type
Log2 (Fold vs. median)
10
30
50
* RUNX1-RUNX1T1 | CBFB-MYH1
KMT2A type
Non-KMT2Ar
KMT2A point
KMT2A PTD
KMT2A−AFDN
KMT2A−AFF1
KMT2A−del
KMT2A−MLLT10
KMT2A−MLLT3
KMT2A translocation (partner unknown)
Unknown
Sample type
Primary pedi AML (BCM)
PDX
Primary AML (Stanford)
KMT2Ar status
KMT2Ar
Non-KMT2Ar
Unknown

### Figure S4

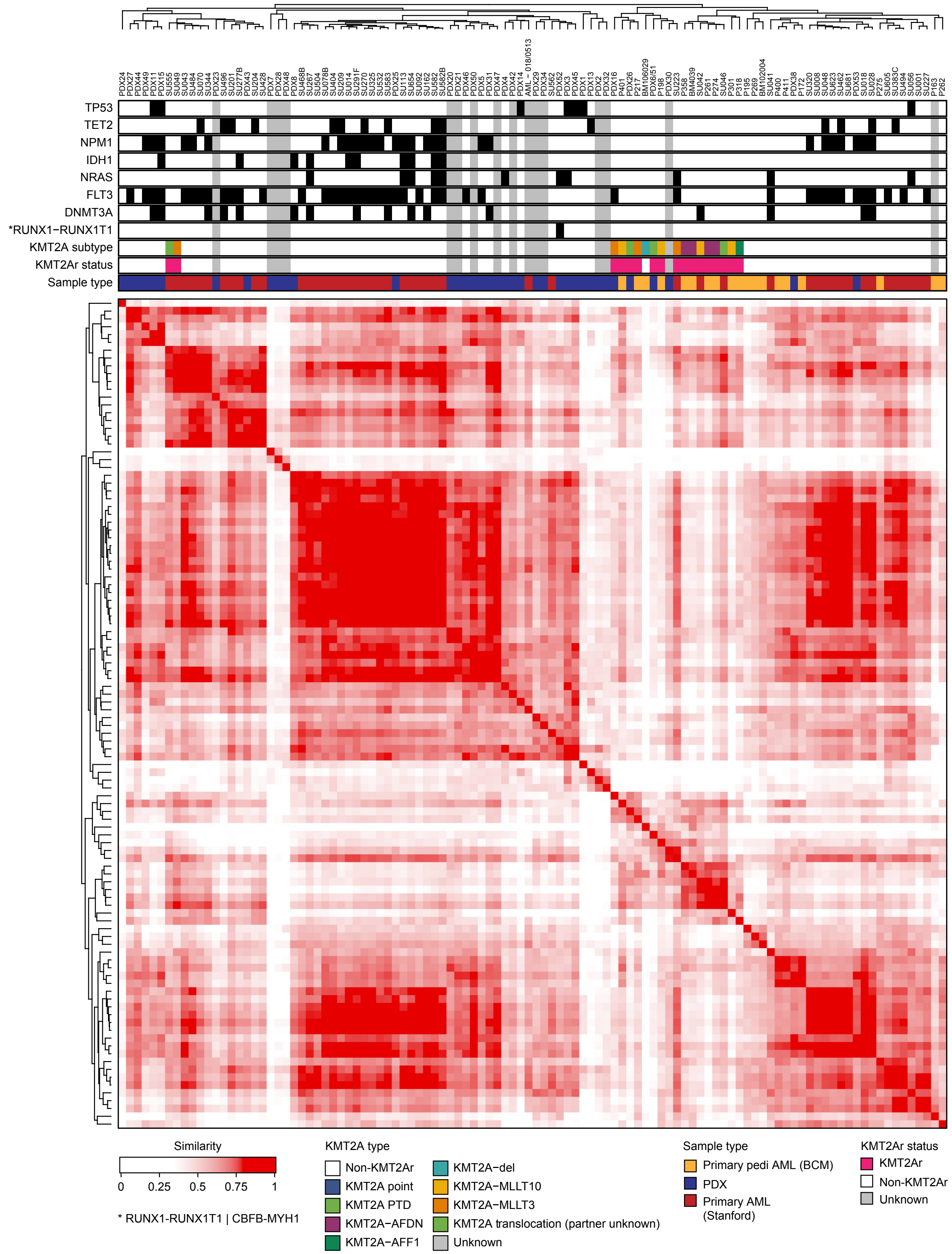
TP53
TET2
NPM1
IDH1
NRAS
FLT3
DNMT3A
*RUNX1−RUNX1T1
KMT2A subtype
KMT2Ar status
Sample type
Similarity
0
0.25
0.5
0.75
1
* RUNX1-RUNX1T1 | CBFB-MYH1
KMT2A type
Non-KMT2Ar
KMT2A point
KMT2A PTD
KMT2A−AFDN
KMT2A−AFF1
KMT2A−del
KMT2A−MLLT10
KMT2A−MLLT3
KMT2A translocation (partner unknown)
Unknown
Sample type
Primary pedi AML (BCM)
PDX
Primary AML (Stanford)
KMT2Ar status
KMT2Ar
Non-KMT2Ar
Unknown

### Figure S5

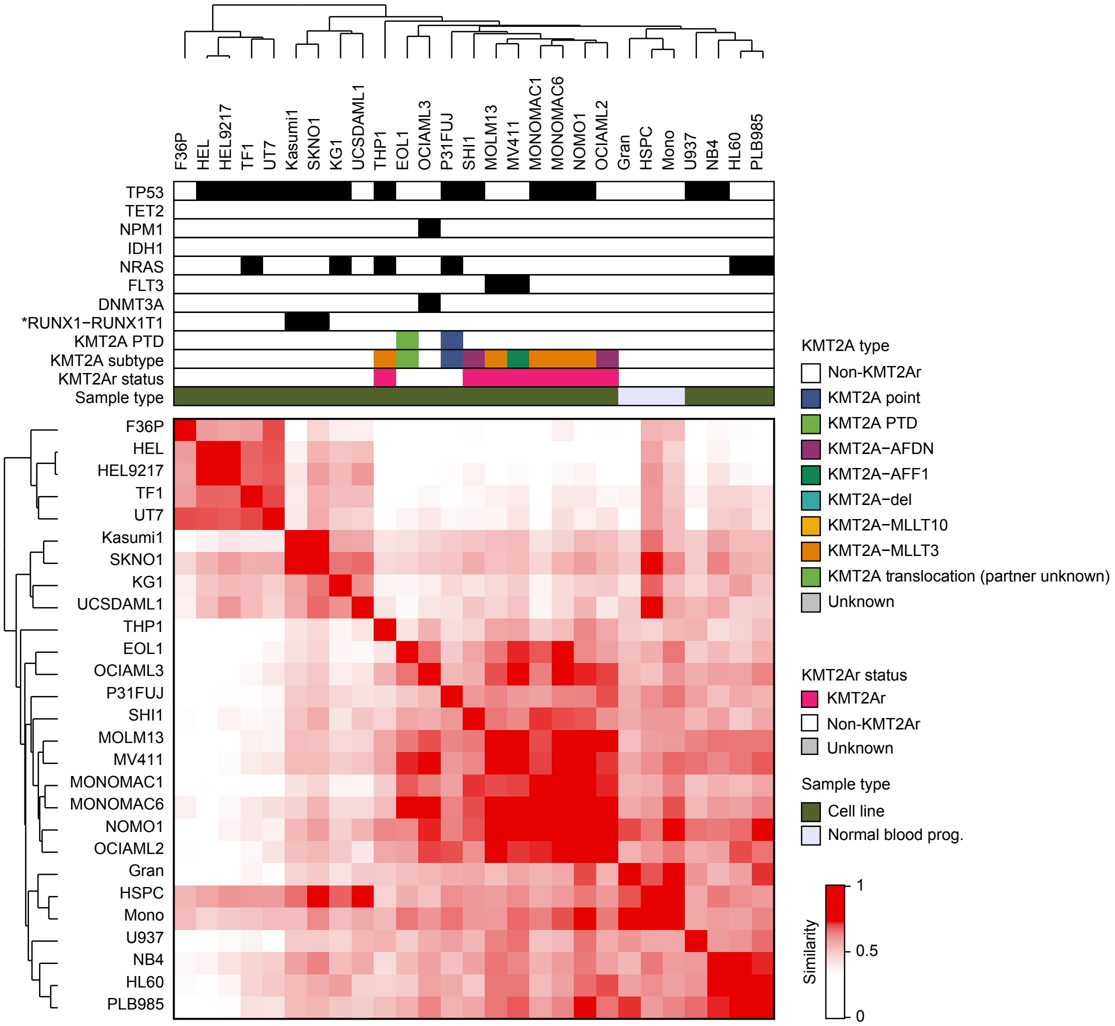


* RUNX1-RUNX1T1 | CBFB-MYH1

### Figure S6

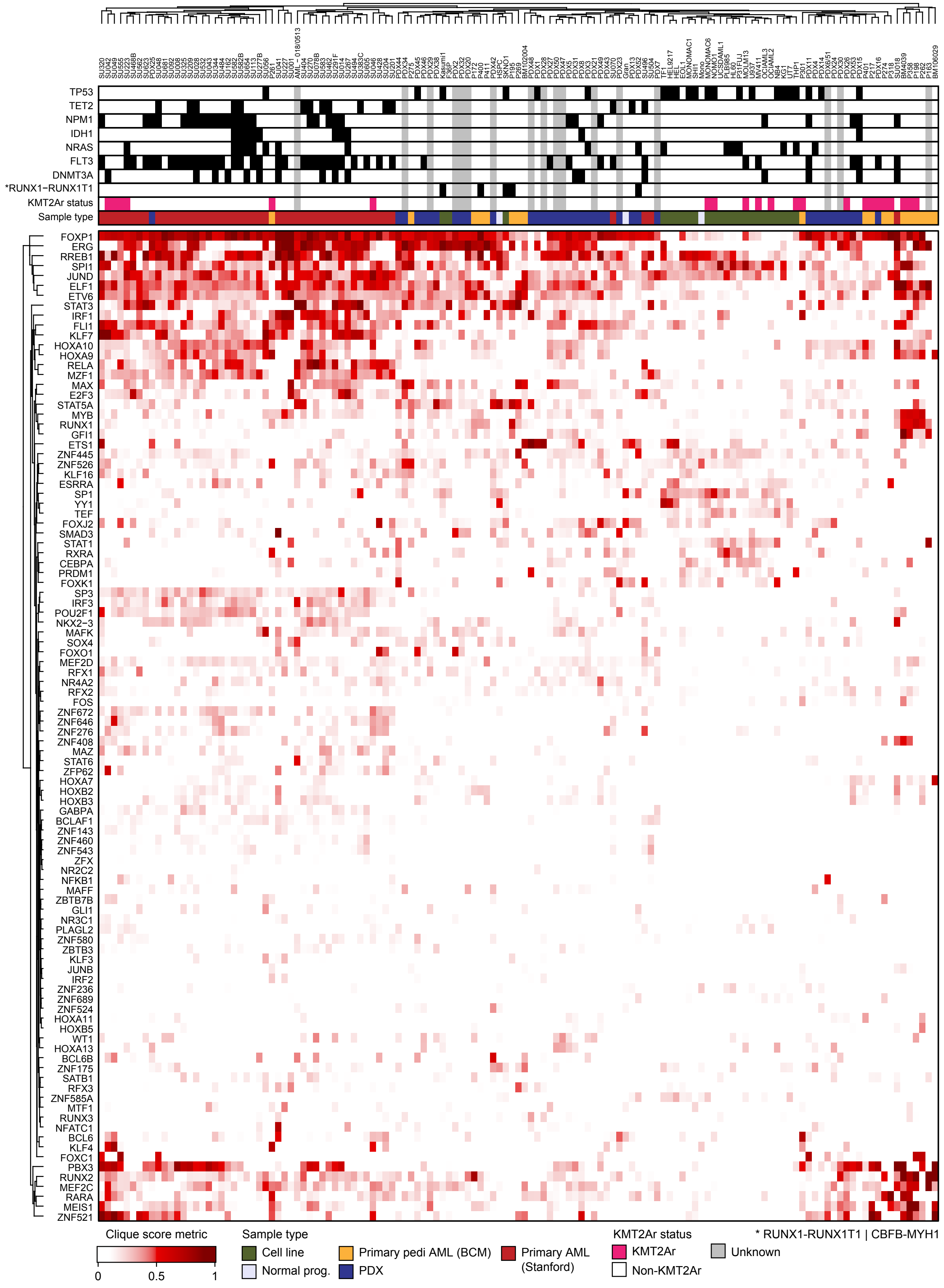

TP53
TET2
NPM1
IDH1
NRAS
FLT3
DNMT3A
*RUNX1−RUNX1T1
KMT2Ar status
Sample type
FOXP1
ERG
RREB1
SPI1
JUND
ELF1
ETV6
STAT3
IRF1
FLI1
KLF7
HOXA10
HOXA9
RELA
MZF1
MAX
E2F3
STAT5A
MYB
RUNX1
GFI1
ETS1
ZNF445
ZNF526
KLF16
ESRRA
SP1
YY1
TEF
FOXJ2
SMAD3
STAT1
RXRA
CEBPA
PRDM1
FOXK1
SP3
IRF3
POU2F1
NKX2−3
MAFK
SOX4
FOXO1
MEF2D
RFX1
NR4A2
RFX2
FOS
ZNF672
ZNF646
ZNF276
ZNF408
MAZ
STAT6
ZFP62
HOXA7
HOXB2
HOXB3
GABPA
BCLAF1
ZNF143
ZNF460
ZNF543
ZFX
NR2C2
NFKB1
MAFF
ZBTB7B
GLI1
NR3C1
PLAGL2
ZNF580
ZBTB3
KLF3
JUNB
IRF2
ZNF236
ZNF689
ZNF524
HOXA11
HOXB5
WT1
HOXA13
BCL6B
ZNF175
SATB1
RFX3
ZNF585A
MTF1
RUNX3
NFATC1
BCL6
KLF4
FOXC1
PBX3
RUNX2
MEF2C
RARA
MEIS1
ZNF521
Clique score metric
0
0.5
1
Sample type
Cell line
Normal prog.
Primary pedi AML (BCM)
PDX
Primary AML (Stanford)
KMT2Ar status
KMT2Ar
Non-KMT2Ar
Unknown
* RUNX1-RUNX1T1 | CBFB-MYH1

### Figure S7

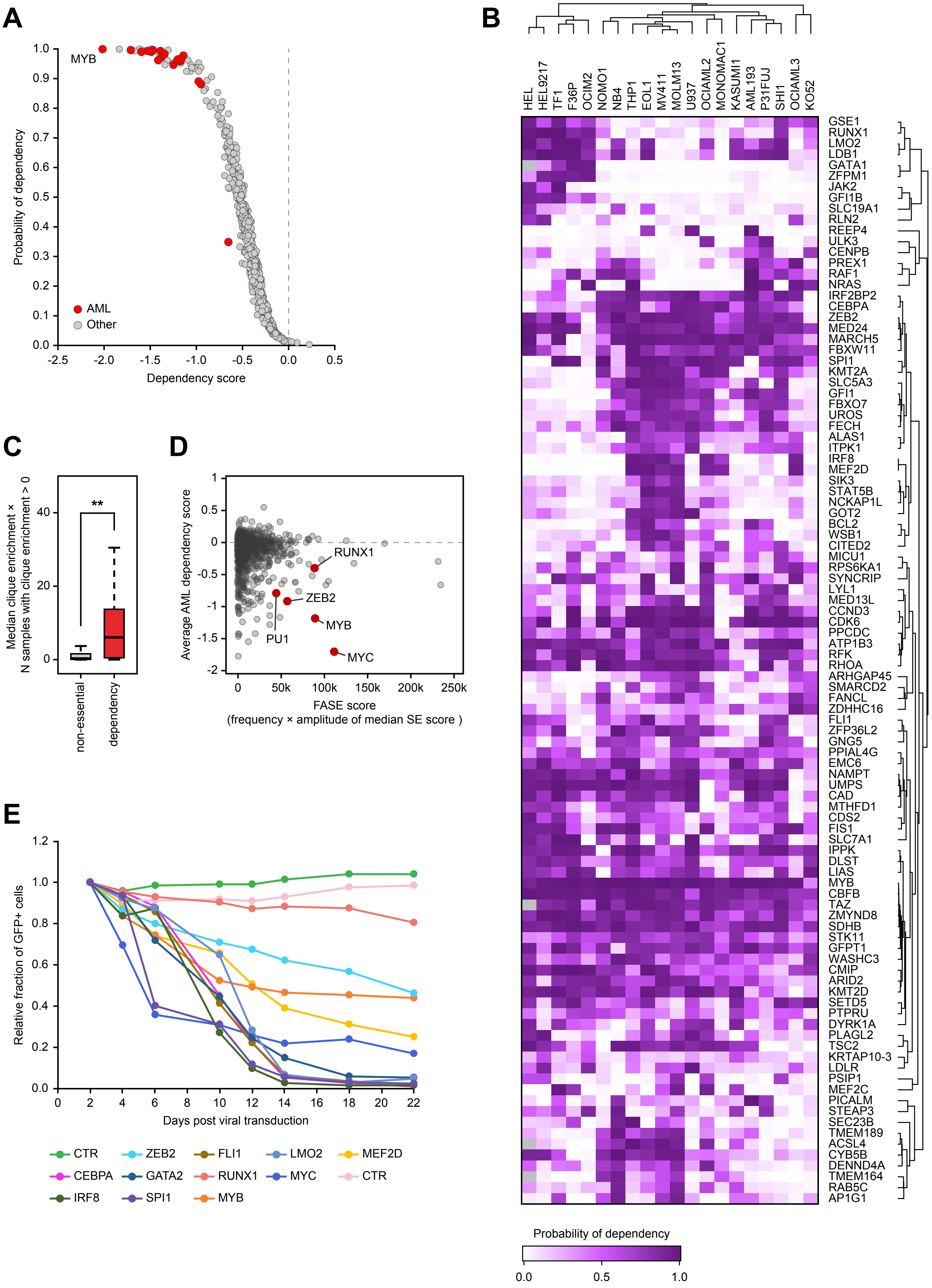

A
B
C
D
E
MYB
AML
Other
Probability of dependency
Dependency score
**
Median clique enrichment × N samples with clique enrichment > 0
non-essential
dependency
Average AML dependency score
FASE score
(frequency × amplitude of median SE score )
RUNX1
ZEB2
PU1
MYB
MYC
Relative fraction of GFP+ cells
Days post viral transduction
CTR
ZEB2
FLI1
LMO2
MEF2D
CEBPA
GATA2
RUNX1
MYC
CTR
IRF8
SPI1
MYB
HEL
HEL9217
TF1
F36P
OCIM2
NOMO1
NB4
THP1
EOL1
MV411
MOLM13
U937
OCIAML2
MONOMAC1
KASUMI1
AML193
P31FUJ
SHI1
OCIAML3
KO52
GSE1
RUNX1
LMO2
LDB1
GATA1
ZFPM1
JAK2
GFI1B
SLC19A1
RLN2
REEP4
ULK3
CENPB
PREX1
RAF1
NRAS
IRF2BP2
CEBPA
ZEB2
MED24
MARCH5
FBXW11
SPI1
KMT2A
SLC5A3
GFI1
FBXO7
UROS
FECH
ALAS1
ITPK1
IRF8
MEF2D
SIK3
STAT5B
NCKAP1L
GOT2
BCL2
WSB1
CITED2
MICU1
RPS6KA1
SYNCRIP
LYL1
MED13L
CCND3
CDK6
PPCDC
ATP1B3
RFK
RHOA
ARHGAP45
SMARCD2
FANCL
ZDHHC16
FLI1
ZFP36L2
GNG5
PPIAL4G
EMC6
NAMPT
UMPS
CAD
MTHFD1
CDS2
FIS1
SLC7A1
IPPK
DLST
LIAS
MYB
CBFB
TAZ
ZMYND8
SDHB
STK11
GFPT1
WASHC3
CMIP
ARID2
KMT2D
SETD5
PTPRU
DYRK1A
PLAGL2
TSC2
KRTAP10-3
LDLR
PSIP1
MEF2C
PICALM
STEAP3
SEC23B
TMEM189
ACSL4
CYB5B
DENND4A
TMEM164
RAB5C
AP1G1
Probability of dependency

### Figure S8

| CR TF | Enriched AML dependency | Top 2245 SEs / 77% recall | TF (Lambert *et al.*) | ChIP-seq | RNA-seq after k/o | Which criteria not met | Reason for inclusion/exclusion |
|---|---|---|---|---|---|---|---|
| CEBPA | Y | Y | Y | Y | N | | |
| E2F3 | Y | Y | Y | Y | N | | |
| FLI1 | Y | Y | Y | Y | Y | | |
| FOSL2 | Y | Y | Y | Y | N | | |
| GFI1 | Y | Y | Y | Y | N | | |
| HHEX | Y | Y | Y | N | N | | |
| IRF8 | Y | Y | Y | Y | Y | | |
| LYL1 | Y | Y | Y | Y | Y | | |
| MEF2C | Y | Y | Y | Y | Y | | |
| MEF2D | Y | Y | Y | Y | Y | | |
| MEIS1 | Y | Y | Y | Y | Y | | |
| MYB | Y | Y | Y | Y | Y | | |
| PLAGL2 | Y | Y | Y | Y | N | | |
| RUNX1 | Y | Y | Y | Y | Y | | |
| RUNX2 | Y | Y | Y | Y | Y | | |
| RXRA | Y | Y | Y | Y | N | | |
| SP1 | Y | Y | Y | Y | Y | | |
| SPI1 | Y | Y | Y | Y | Y | | |
| SREBF1 | Y | Y | Y | Y | N | | |
| STAT5B | Y | Y | Y | Y | N | | |
| TFAP4 | Y | Y | Y | Y | N | | |
| ZEB2 | Y | Y | Y | Y | Y | | |
| ZFPM1 | Y | Y | Y | N | N | | |
| ZNF281 | Y | Y | Y | Y | N | | |
| LMO2 | Y | Y | N | Y | Y | Not classified as a TF by Lambert *et al.*; acts as a scaffolding cofactor for TF complexes; no evidence of sequence-specific DNA binding | Transcriptional regulator; leukemia oncogene; strong dependency and SE |
| ZMYND8 | Y | Y | N | Y | Y | Not classified as a TF by Lambert at al; acts as a transcriptional corepressor; no evidence of sequence-specific DNA binding | Strong AML Dependency; one of the top AML SEs |
| GATA2 | N | Y | Y | Y | N | Weak drop out scores in the screen | Low throughput validation by CRISPR knockout shows strong dependency; appears to be a false-negative in the genome-wide screen |
| MAX | N | Y | Y | Y | Y | Common essential gene, not a preferential AML dependency | Forms a heterodimer with MYC; strong dependency and SE |
| MYC | N | Y | Y | Y | Y | Classified as a common essential gene | AML oncogene; strong dependency enrichment in AML despite classification as a common essential; one of the top SEs |
| ETV6 | N | Y | Y | Y | N | Weak dependency scores in the screen | One of the top AML SEs; unable to achieve knock-out in CRISPR validation experiments, possibly a false-negative in the genome-wide screen |
| HOXA9 | N | Y | Y | Y | Y | Weak dependency scores in most AML cell lines | AML oncogene; strong SE; likely a false-negative in the genome-wide screen due to HOXA cross-compensation |
| CENPB | Y | Y | Y | N | N | | Component of centromeric heterochromatin participating in centromere formation and kinetochore assembly; excluded as unlikely to be a TF |
| ARID2 | Y | Y | Y | N | N | | Member of the SWI/SNF chromatin remodeling complex; excluded as unlikely to be a sequence-specific TF |

### Figure S9

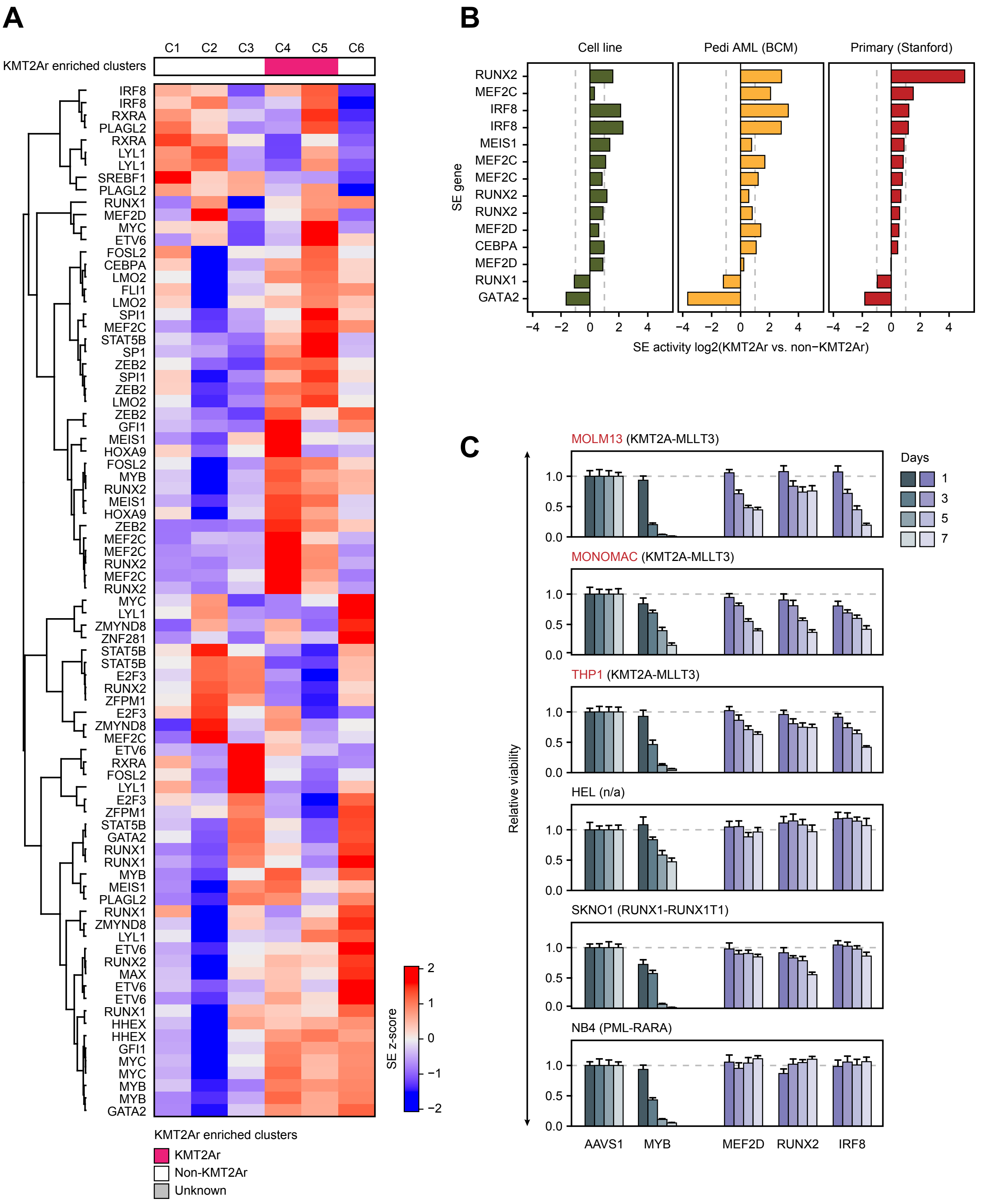

A
C1 C2 C3 C4 C5 C6
KMT2Ar enriched clusters
IRF8
IRF8
RXRA
PLAGL2
RXRA
LYL1
LYL1
SREBF1
PLAGL2
RUNX1
MEF2D
MYC
ETV6
FOSL2
CEBPA
LMO2
FLI1
LMO2
SPI1
MEF2C
STAT5B
SP1
ZEB2
SPI1
ZEB2
LMO2
ZEB2
GFI1
MEIS1
HOXA9
FOSL2
MYB
RUNX2
MEIS1
HOXA9
ZEB2
MEF2C
MEF2C
RUNX2
MEF2C
RUNX2
MYC
LYL1
ZMYND8
ZNF281
STAT5B
STAT5B
E2F3
RUNX2
ZFPM1
E2F3
ZMYND8
MEF2C
ETV6
RXRA
FOSL2
LYL1
E2F3
ZFPM1
STAT5B
GATA2
RUNX1
RUNX1
MYB
MEIS1
PLAGL2
RUNX1
ZMYND8
LYL1
ETV6
RUNX2
MAX
ETV6
ETV6
RUNX1
HHEX
HHEX
GFI1
MYC
MYC
MYB
MYB
GATA2
SE z-score
2
1
0
−1
−2
KMT2Ar enriched clusters
KMT2Ar
Non-KMT2Ar
Unknown
B
Cell line
Pedi AML (BCM)
Primary (Stanford)
SE gene
RUNX2
MEF2C
IRF8
IRF8
MEIS1
MEF2C
MEF2C
RUNX2
RUNX2
MEF2D
CEBPA
MEF2D
RUNX1
GATA2
−4 −2 0 2 4
SE activity log2(KMT2Ar vs. non−KMT2Ar)
C
MOLM13 (KMT2A-MLLT3)
MONOMAC (KMT2A-MLLT3)
THP1 (KMT2A-MLLT3)
HEL (n/a)
SKNO1 (RUNX1-RUNX1T1)
NB4 (PML-RARA)
Relative viability
1.0
0.5
0.0
Days
1
3
5
7
AAVS1
MYB
MEF2D
RUNX2
IRF8

### Figure S10

A

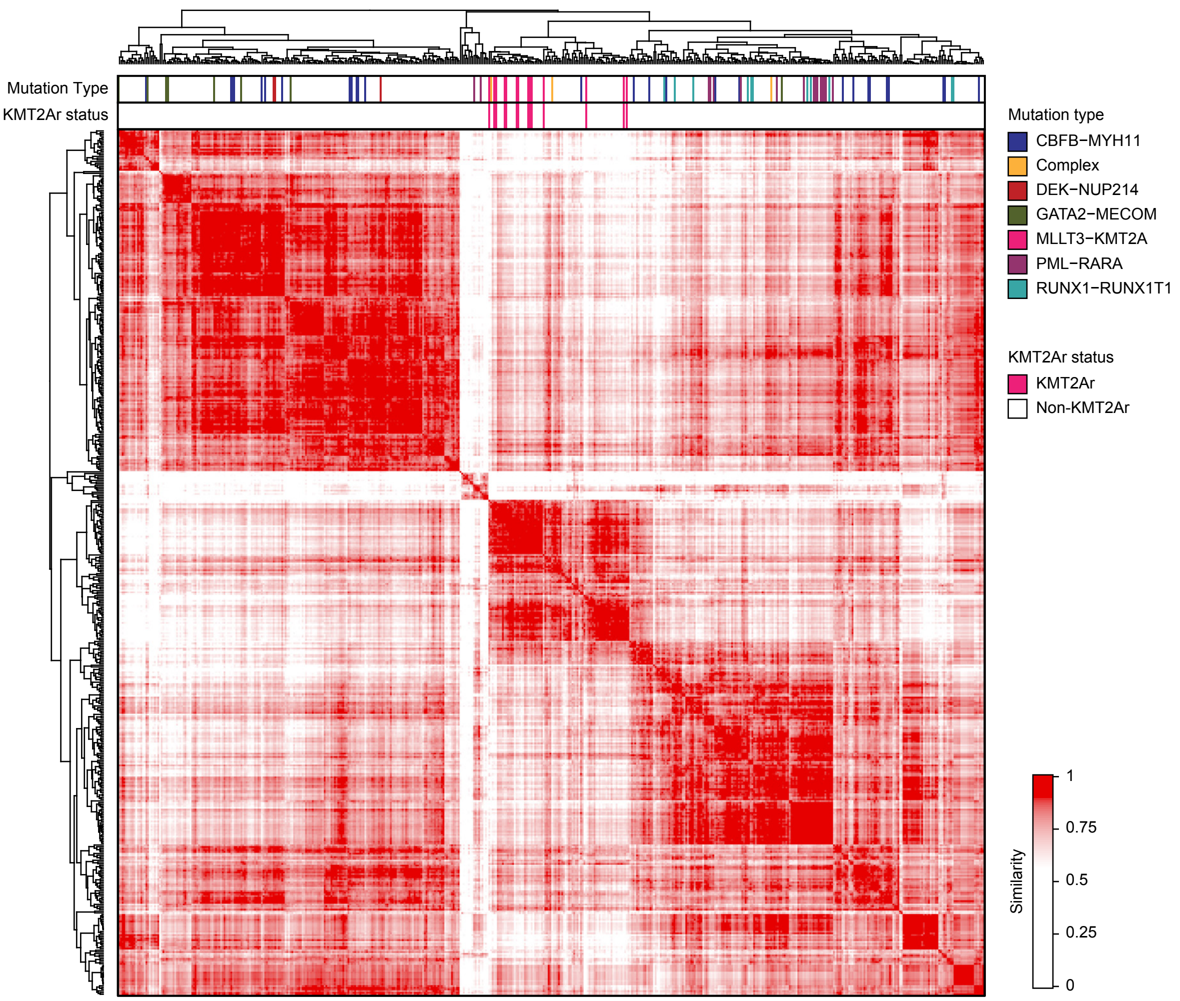


B

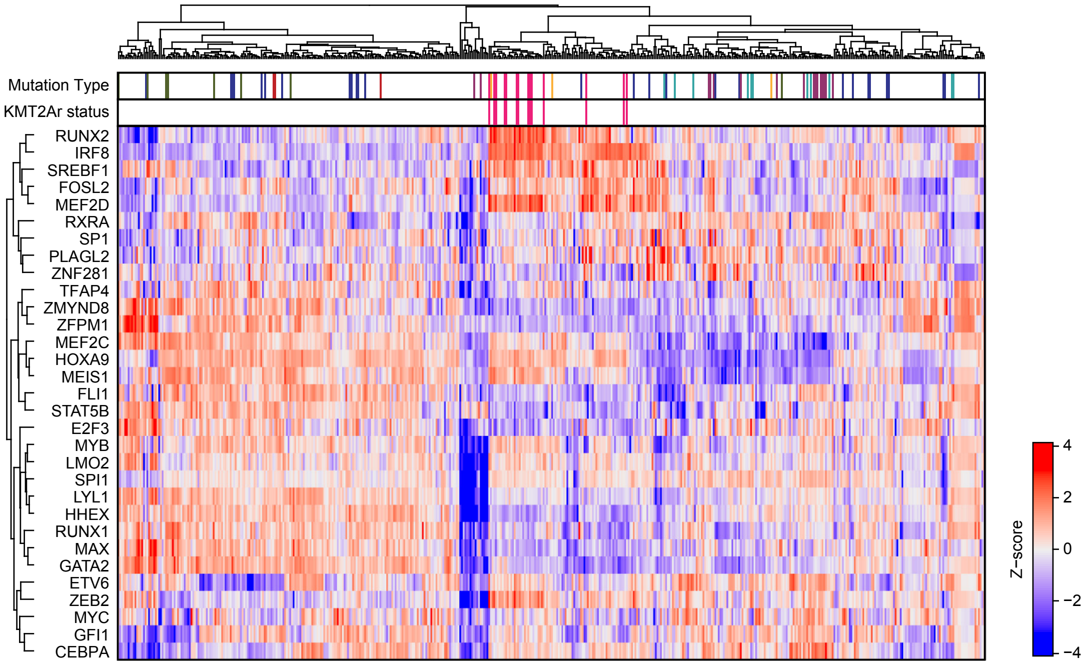

### Figure S11

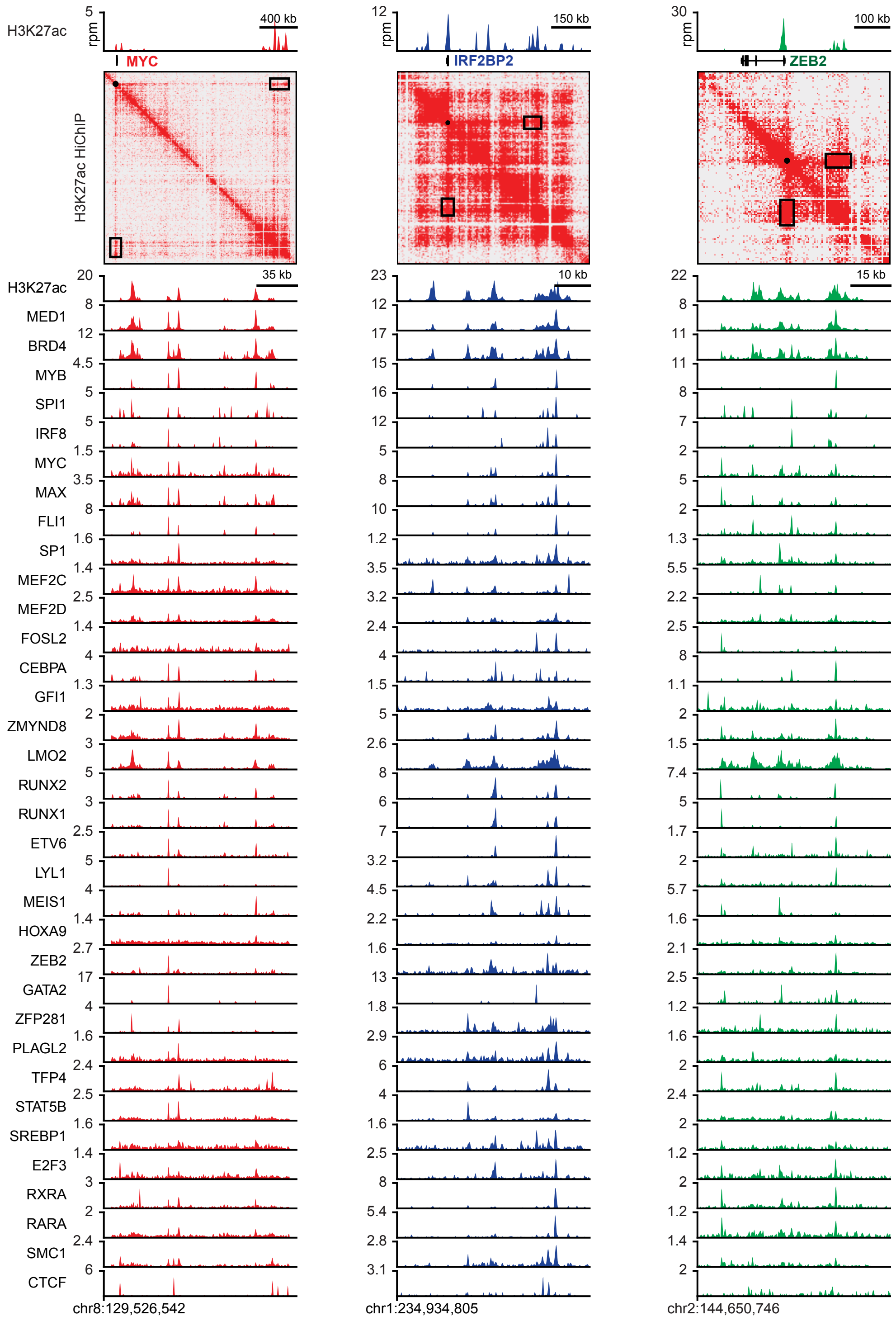
H3K27ac
rpm
MYC
IRF2BP2
ZEB2
400 kb
150 kb
100 kb
H3K27ac HiChIP
35 kb
10 kb
15 kb
H3K27ac
MED1
BRD4
MYB
SPI1
IRF8
MYC
MAX
FLI1
SP1
MEF2C
MEF2D
FOSL2
CEBPA
GFI1
ZMYND8
LMO2
RUNX2
RUNX1
ETV6
LYL1
MEIS1
HOXA9
ZEB2
GATA2
ZFP281
PLAGL2
TFP4
STAT5B
SREBP1
E2F3
RXRA
RARA
SMC1
CTCF
chr8:129,526,542
chr1:234,934,805
chr2:144,650,746

### Figure S12

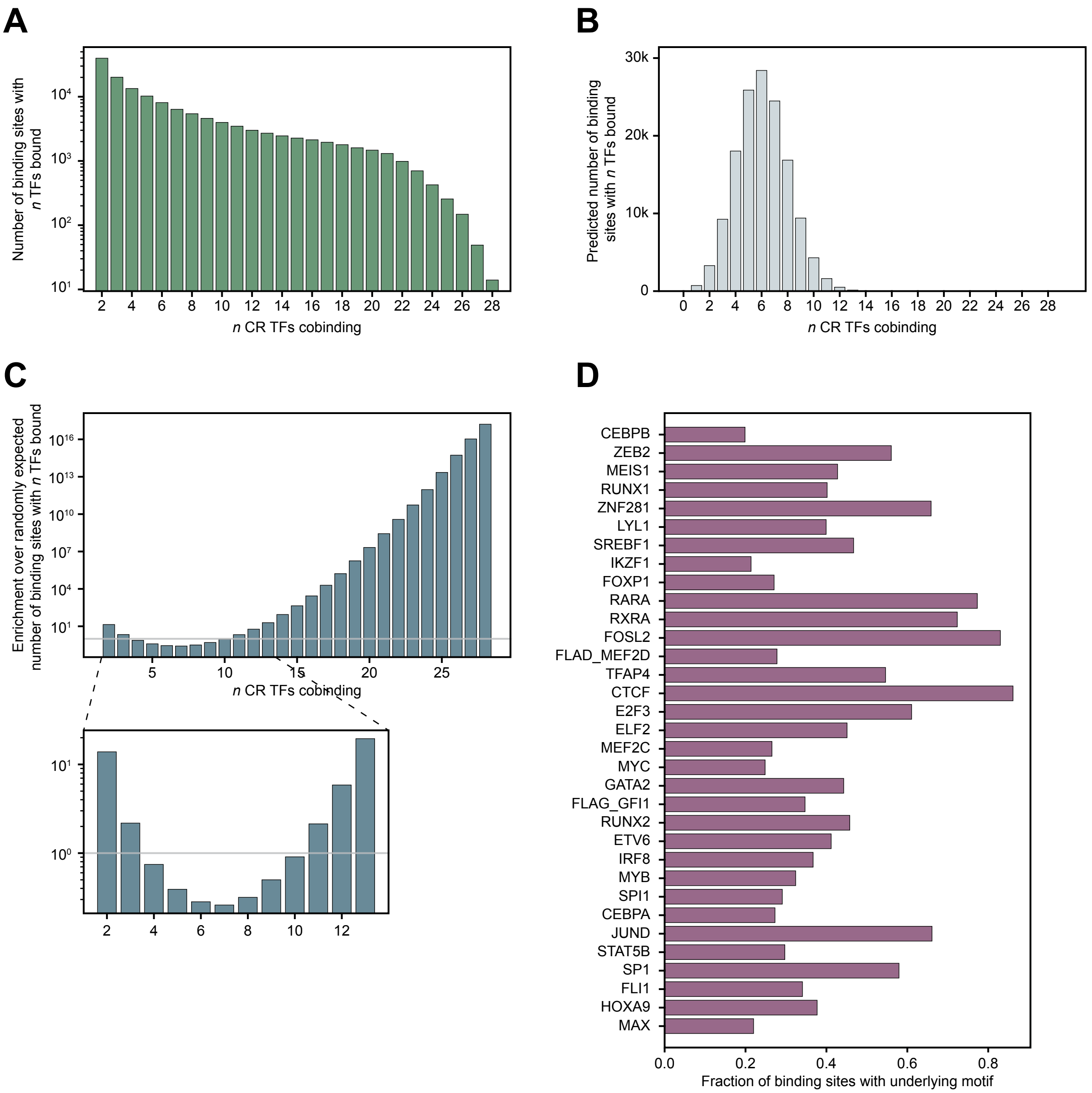
A
B
C
D
Number of binding sites with n TFs bound
n CR TFs cobinding
Predicted number of binding sites with n TFs bound
Enrichment over randomly expected number of binding sites with n TFs bound
CEBPB
ZEB2
MEIS1
RUNX1
ZNF281
LYL1
SREBF1
IKZF1
FOXP1
RARA
RXRA
FOSL2
FLAD_MEF2D
TFAP4
CTCF
E2F3
ELF2
MEF2C
MYC
GATA2
FLAG_GFI1
RUNX2
ETV6
IRF8
MYB
SPI1
CEBPA
JUND
STAT5B
SP1
FLI1
HOXA9
MAX
Fraction of binding sites with underlying motif

### Figure S13

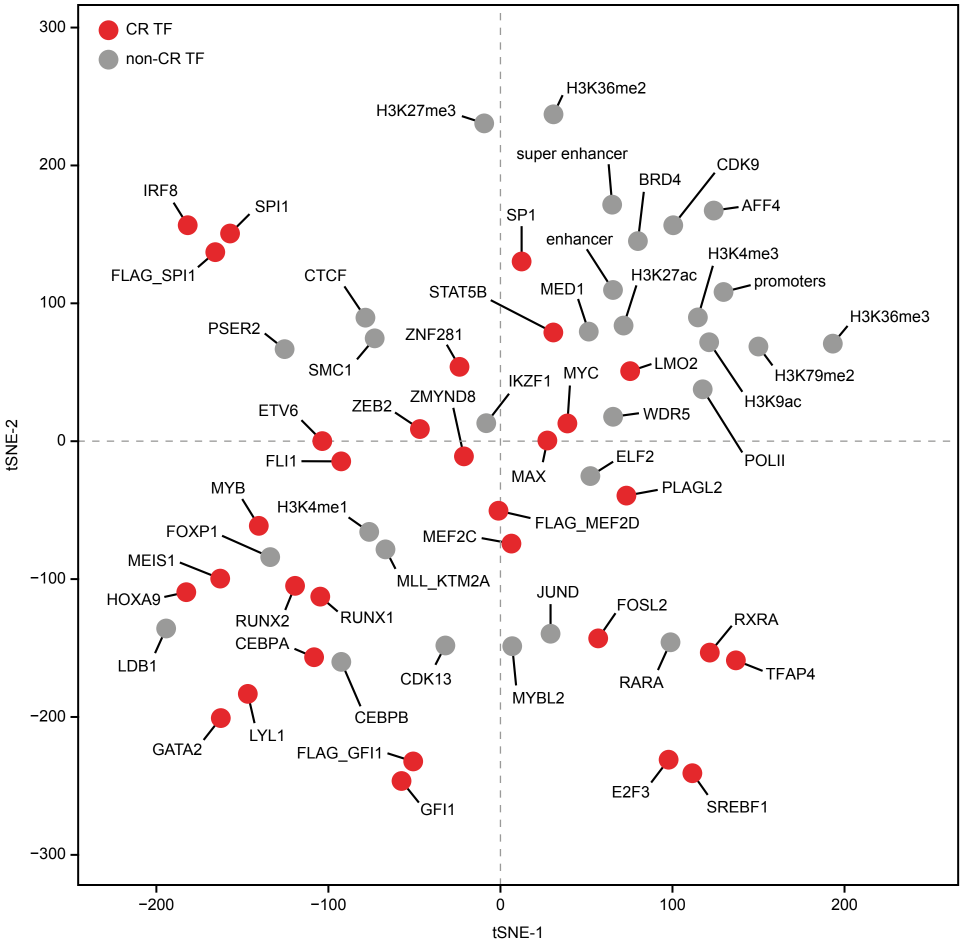

CR TF
non-CR TF
tSNE-2
tSNE-1
300
200
100
0
−100
−200
−300
−200
−100
0
100
200
H3K27me3
H3K36me2
super enhancer
CDK9
BRD4
AFF4
IRF8
SPI1
FLAG_SPI1
SP1
enhancer
H3K4me3
H3K27ac
promoters
CTCF
STAT5B
MED1
H3K36me3
PSER2
SMC1
ZNF281
LMO2
H3K79me2
IKZF1
MYC
H3K9ac
ETV6
ZEB2
ZMYND8
WDR5
FLI1
MAX
ELF2
POLII
MYB
H3K4me1
PLAGL2
FOXP1
FLAG_MEF2D
MEF2C
MEIS1
MLL_KTM2A
HOXA9
RUNX2
RUNX1
JUND
FOSL2
RXRA
LDB1
CEBPA
CDK13
MYBL2
RARA
TFAP4
GATA2
LYL1
CEBPB
FLAG_GFI1
GFI1
E2F3
SREBF1

### Figure S14

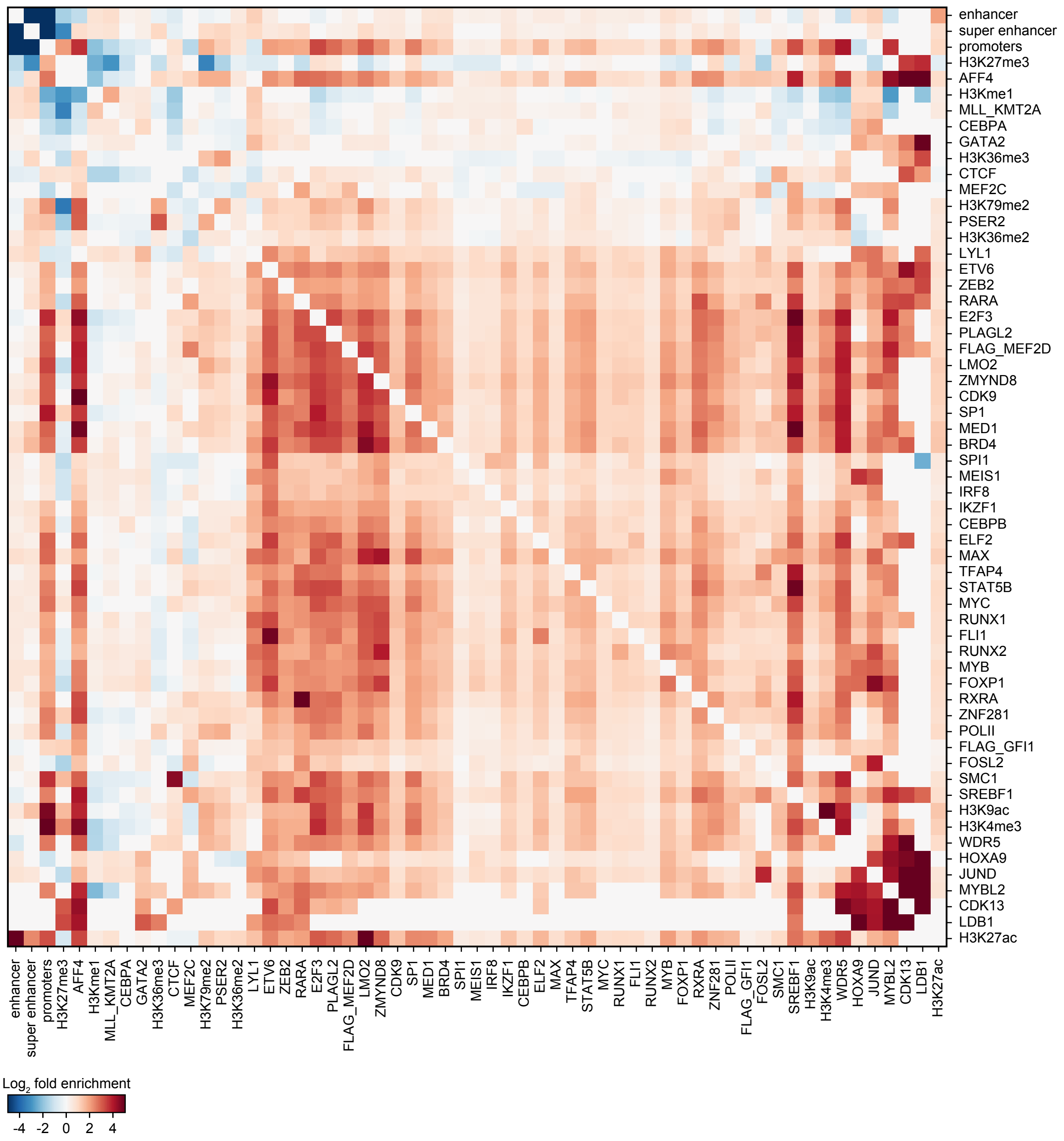

enhancer
super enhancer
promoters
H3K27me3
AFF4
H3Kme1
MLL_KMT2A
CEBPA
GATA2
H3K36me3
CTCF
MEF2C
H3K79me2
PSER2
H3K36me2
LYL1
ETV6
ZEB2
RARA
E2F3
PLAGL2
FLAG_MEF2D
LMO2
ZMYND8
CDK9
SP1
MED1
BRD4
SPI1
MEIS1
IRF8
IKZF1
CEBPB
ELF2
MAX
TFAP4
STAT5B
MYC
RUNX1
FLI1
RUNX2
MYB
FOXP1
RXRA
ZNF281
POLII
FLAG_GFI1
FOSL2
SMC1
SREBF1
H3K9ac
H3K4me3
WDR5
HOXA9
JUND
MYBL2
CDK13
LDB1
H3K27ac
Log2 fold enrichment
-4
-2
0
2
4

### Figure S15

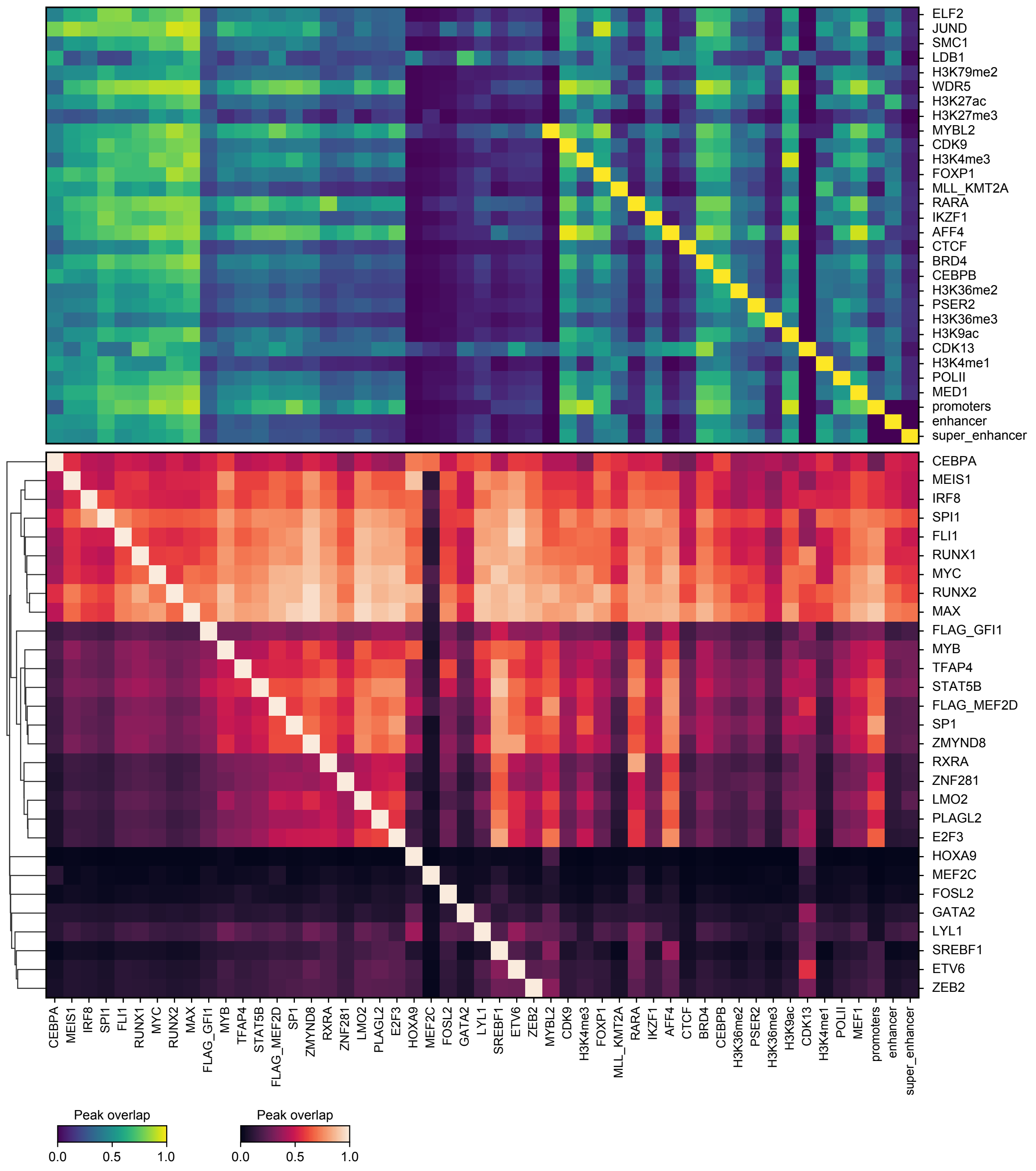
ELF2
JUND
SMC1
LDB1
H3K79me2
WDR5
H3K27ac
H3K27me3
MYBL2
CDK9
H3K4me3
FOXP1
MLL_KMT2A
RARA
IKZF1
AFF4
CTCF
BRD4
CEBPB
H3K36me2
PSER2
H3K36me3
H3K9ac
CDK13
H3K4me1
POLII
MED1
promoters
enhancer
super_enhancer
CEBPA
MEIS1
IRF8
SPI1
FLI1
RUNX1
MYC
RUNX2
MAX
FLAG_GFI1
MYB
TFAP4
STAT5B
FLAG_MEF2D
SP1
ZMYND8
RXRA
ZNF281
LMO2
PLAGL2
E2F3
HOXA9
MEF2C
FOSL2
GATA2
LYL1
SREBF1
ETV6
ZEB2
Peak overlap
0.0
0.5
1.0
Peak overlap
0.0
0.5
1.0

### Figure S16

A

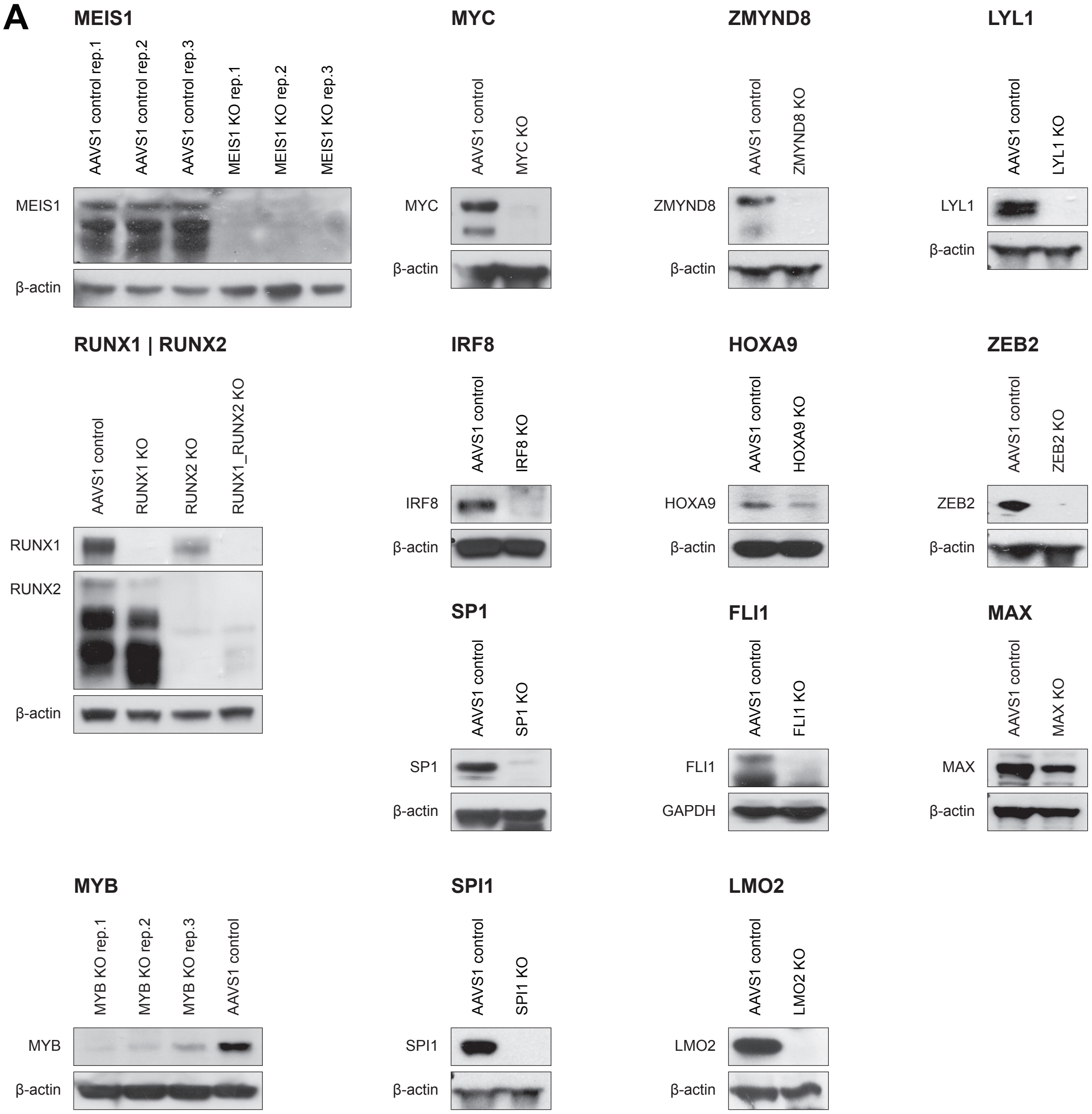


B

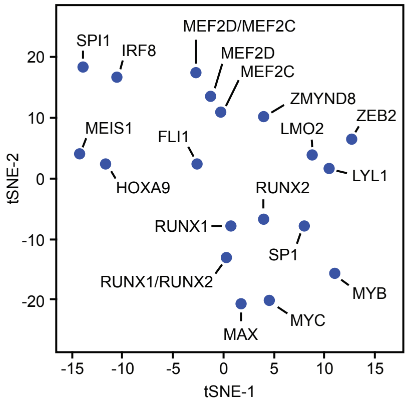

### Figure S17

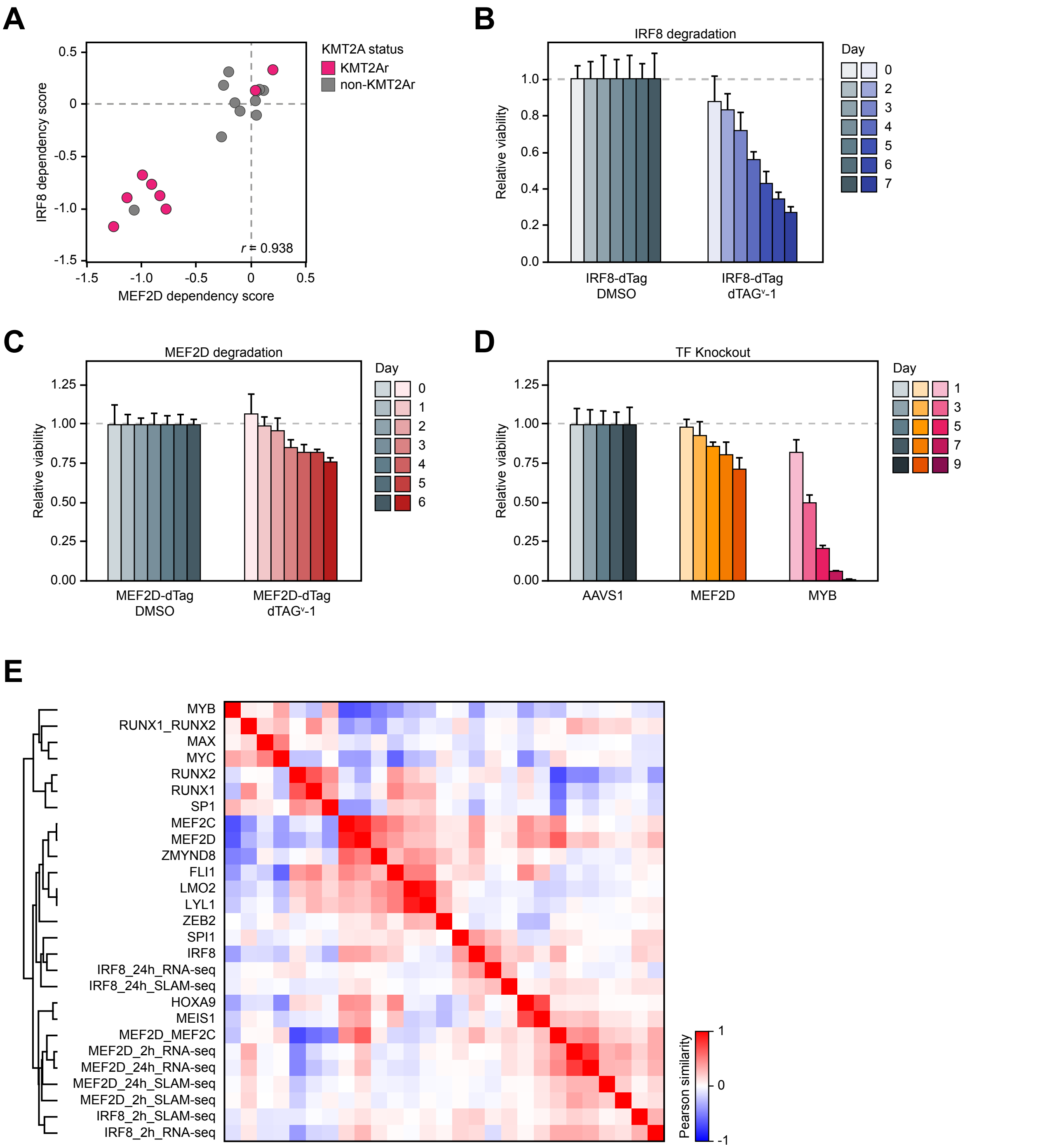
A
KMT2A status
KMT2Ar
non-KMT2Ar
IRF8 dependency score
MEF2D dependency score
r = 0.938
B
IRF8 degradation
Relative viability
Day
IRF8-dTag DMSO
IRF8-dTag dTAGᵛ-1
C
MEF2D degradation
Relative viability
Day
MEF2D-dTag DMSO
MEF2D-dTag dTAGᵛ-1
D
TF Knockout
Relative viability
Day
AAVS1
MEF2D
MYB
E
MYB
RUNX1_RUNX2
MAX
MYC
RUNX2
RUNX1
SP1
MEF2C
MEF2D
ZMYND8
FLI1
LMO2
LYL1
ZEB2
SPI1
IRF8
IRF8_24h_RNA-seq
IRF8_24h_SLAM-seq
HOXA9
MEIS1
MEF2D_MEF2C
MEF2D_2h_RNA-seq
MEF2D_24h_RNA-seq
MEF2D_24h_SLAM-seq
MEF2D_2h_SLAM-seq
IRF8_2h_SLAM-seq
IRF8_2h_RNA-seq
Pearson similarity

### Figure S18

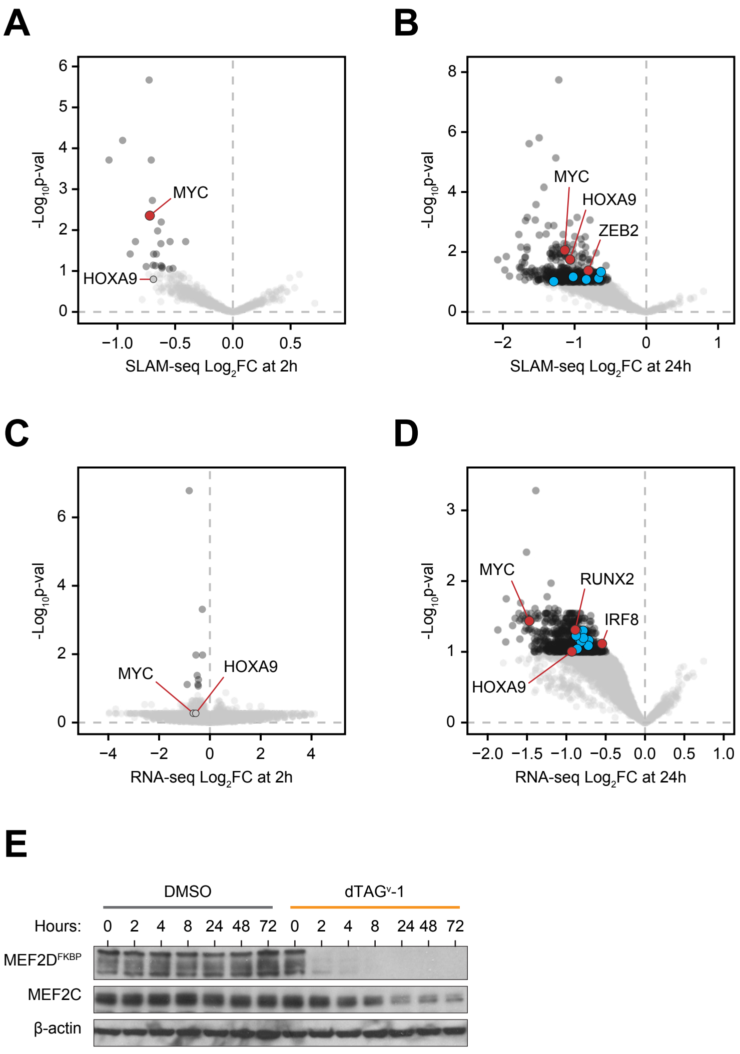

A
-Log10p-val
MYC
HOXA9
SLAM-seq Log2FC at 2h
B
-Log10p-val
MYC
HOXA9
ZEB2
SLAM-seq Log2FC at 24h
C
-Log10p-val
MYC
HOXA9
RNA-seq Log2FC at 2h
D
-Log10p-val
MYC
RUNX2
IRF8
HOXA9
RNA-seq Log2FC at 24h
E
DMSO
dTAGV-1
Hours: 0 2 4 8 24 48 72 0 2 4 8 24 48 72
MEF2DFKBP
MEF2C
β-actin

### Figure S19

## A

| Gene ID | MV411 dep. probability | MV411 drop out score | Pan-lethal | SE in MV411 | TF | 2 hrs SLAM-seq | | | 24 hrs SLAM-seq | | | 2 hrs RNA-seq | | | 24 hrs RNA-seq | | |
|---|---|---|---|---|---|---|---|---|---|---|---|---|---|---|---|---|---|
| | | | | | | $log_2FC$ | pvalue | padj | $log_2FC$ | pvalue | padj | $log_2FC$ | pval | padj | $log_2FC$ | pval | padj |
| FUT4 | 0.009 | -9E-04 | FALSE | 1 | | -0.725 | 2.6E-09 | 2.1E-06 | -0.933 | 0.002 | 0.043 | -0.452 | 2.9E-04 | 0.232 | -0.757 | 0.004 | 0.089 |
| ZFP36 | 0.008 | 0.012 | FALSE | 1 | | -0.955 | 1.6E-07 | 6.4E-05 | -1.630 | 3.2E-09 | 2.4E-06 | -0.659 | 0.038 | 0.534 | -1.477 | 0.025 | 0.124 |
| BHLHE40 | 0.001 | 0.141 | FALSE | 1 | 1 | -0.707 | 8.3E-07 | 1.9E-04 | -1.035 | 0.006 | 0.062 | -0.500 | 0.018 | 0.534 | -1.019 | 0.002 | 0.072 |
| PTGER4 | 0.001 | 0.143 | FALSE | 1 | | -1.073 | 9.5E-07 | 1.9E-04 | -1.128 | 0.009 | 0.069 | -0.944 | 0.003 | 0.446 | -1.086 | 0.003 | 0.082 |
| KLF10 | 0.002 | 0.101 | FALSE | | 1 | -0.697 | 1.2E-05 | 0.002 | -0.697 | 0.048 | 0.143 | -0.545 | 0.009 | 0.521 | -0.757 | 0.073 | 0.167 |
| MYC | 1.000 | -1.935 | TRUE | 1 | 1 | -0.720 | 3.3E-05 | 0.004 | -1.136 | 1.4E-04 | 0.009 | -0.655 | 0.075 | 0.534 | -1.470 | 2.7E-04 | 0.037 |
| F3 | 0.141 | -0.259 | FALSE | | | -0.620 | 5.5E-05 | 0.006 | -0.828 | 0.010 | 0.070 | -0.541 | 0.064 | 0.534 | -0.953 | 0.002 | 0.076 |
| TNFAIP3 | 0.003 | 0.093 | FALSE | | | -0.652 | 1.0E-04 | 0.010 | -0.681 | 0.093 | 0.209 | -0.520 | 0.006 | 0.503 | -1.051 | 4.9E-04 | 0.046 |
| TXNIP | 0.438 | -0.449 | FALSE | | | -0.544 | 2.4E-04 | 0.019 | -1.039 | 6.0E-04 | 0.019 | -0.468 | 0.058 | 0.534 | -0.960 | 0.001 | 0.058 |
| PMAIP1 | 4E-04 | 0.252 | FALSE | 1 | | -0.844 | 2.3E-04 | 0.019 | -0.962 | 0.003 | 0.046 | -0.785 | 8.5E-04 | 0.299 | -1.277 | 0.002 | 0.065 |
| PIM1 | 0.001 | 0.138 | FALSE | 1 | | -0.409 | 2.6E-04 | 0.019 | -0.813 | 0.003 | 0.047 | -0.455 | 0.008 | 0.521 | -0.895 | 0.001 | 0.055 |
| IER2 | 0.003 | 0.072 | FALSE | 1 | | -0.624 | 3.2E-04 | 0.021 | -1.219 | 8.0E-12 | 1.8E-08 | -0.459 | 4.7E-05 | 0.086 | -1.275 | 5.6E-04 | 0.047 |
| ID2 | 0.204 | -0.309 | FALSE | 1 | | -0.890 | 6.1E-04 | 0.038 | -1.545 | 3.5E-06 | 8.4E-04 | -0.905 | 0.024 | 0.534 | -1.212 | 0.007 | 0.097 |
| TP53RK | 0.908 | -0.843 | TRUE | 1 | | -0.663 | 6.6E-04 | 0.038 | -0.880 | 0.017 | 0.090 | -0.478 | 1.1E-04 | 0.164 | -0.799 | 0.002 | 0.072 |
| KLF6 | 5E-04 | 0.222 | FALSE | | 1 | -0.687 | 7.1E-04 | 0.039 | -1.656 | 2.4E-06 | 6.9E-04 | -0.590 | 0.035 | 0.534 | -1.293 | 6.7E-04 | 0.049 |
| KLF11 | 0.109 | -0.228 | FALSE | | 1 | -0.691 | 1.4E-03 | 0.072 | -1.325 | 0.003 | 0.046 | -0.706 | 0.006 | 0.498 | -0.975 | 8.4E-04 | 0.052 |
| CDKN1B | 0.006 | 0.035 | FALSE | 1 | | -0.671 | 1.6E-03 | 0.074 | -1.308 | 2.1E-04 | 0.011 | -0.751 | 0.062 | 0.534 | -1.478 | 6.3E-04 | 0.049 |
| CDC42EP3 | 0.012 | -0.023 | FALSE | 1 | | -0.622 | 1.6E-03 | 0.074 | -0.980 | 0.002 | 0.041 | -0.584 | 0.008 | 0.521 | -1.041 | 5.3E-04 | 0.047 |
| ZBTB33 | 0.032 | -0.103 | FALSE | 1 | 1 | -0.751 | 1.8E-03 | 0.078 | -1.280 | 0.009 | 0.069 | -0.637 | 0.041 | 0.534 | -0.945 | 0.147 | 0.255 |
| RANBP6 | 0.038 | -0.120 | FALSE | | | -0.612 | 1.9E-03 | 0.079 | -0.990 | 0.027 | 0.113 | -0.477 | 0.011 | 0.534 | -0.705 | 0.035 | 0.129 |
| CPEB2 | 2E-04 | 0.293 | FALSE | 1 | | -0.615 | 2.2E-03 | 0.085 | -1.259 | 5.8E-05 | 0.005 | -0.890 | 0.002 | 0.376 | -1.474 | 1.7E-04 | 0.033 |
| SATB2 | 0.063 | -0.168 | FALSE | 1 | 1 | -0.519 | 2.3E-03 | 0.086 | -1.119 | 0.007 | 0.062 | -0.369 | 2.9E-04 | 0.232 | -0.766 | 0.007 | 0.099 |
| DUSP6 | 0.005 | 0.037 | FALSE | 1 | | -0.550 | 2.5E-03 | 0.090 | -0.914 | 0.005 | 0.056 | -0.695 | 0.055 | 0.534 | -1.520 | 1.5E-04 | 0.033 |

## B

| Gene ID | MV411 dep. probability | MV411 drop out score | Pan-lethal | SE in MV411 | TF | 2 hrs SLAM-seq | | | 24 hrs SLAM-seq | | | 2 hrs RNA-seq | | | 24 hrs RNA-seq | | |
|---|---|---|---|---|---|---|---|---|---|---|---|---|---|---|---|---|---|
| | | | | | | $log_2FC$ | pvalue | padj | $log_2FC$ | pvalue | padj | $log_2FC$ | pval | padj | $log_2FC$ | pval | padj |
| MYC | 1.000 | -1.935 | TRUE | 1 | 1 | -0.720 | 3.3E-05 | 0.004 | -1.136 | 1.4E-04 | 0.009 | -0.655 | 0.075 | 0.534 | -1.470 | 2.7E-04 | 0.037 |
| LMO2 | 0.013 | -0.032 | FALSE | 1 | 1 | -0.446 | 0.006 | 0.138 | -0.770 | 0.037 | 0.128 | -0.171 | 0.001 | 0.334 | -0.479 | 0.017 | 0.116 |
| ZEB2 | 0.931 | -0.914 | FALSE | 1 | 1 | -0.425 | 0.008 | 0.152 | -0.809 | 0.002 | 0.041 | -0.322 | 0.010 | 0.527 | -0.877 | 0.001 | 0.059 |
| HOXA9 | 0.305 | -0.374 | FALSE | 1 | 1 | -0.677 | 0.011 | 0.172 | -1.063 | 5.5E-04 | 0.018 | -0.539 | 0.043 | 0.534 | -0.930 | 0.008 | 0.099 |
| MYB | 0.999 | -1.468 | FALSE | 1 | 1 | -0.356 | 0.047 | 0.307 | -1.020 | 0.008 | 0.068 | -0.440 | 0.056 | 0.534 | -0.782 | 0.001 | 0.064 |
| IRF8 | 0.946 | -0.984 | FALSE | 1 | 1 | -0.279 | 0.091 | 0.376 | -0.837 | 0.014 | 0.083 | -0.019 | 0.803 | 0.929 | -0.546 | 0.003 | 0.077 |
| RUNX1 | 0.167 | -0.281 | FALSE | 1 | 1 | -0.195 | 0.152 | 0.480 | -0.671 | 0.041 | 0.134 | -0.487 | 0.004 | 0.450 | -0.861 | 0.005 | 0.091 |
| E2F3 | 0.200 | -0.306 | FALSE | 1 | 1 | -0.234 | 0.176 | 0.499 | -0.498 | 0.098 | 0.213 | -0.444 | 0.001 | 0.299 | -0.840 | 0.002 | 0.065 |
| MEIS1 | 0.376 | -0.414 | FALSE | 1 | 1 | -0.274 | 0.184 | 0.513 | -0.865 | 0.037 | 0.128 | -0.184 | 0.222 | 0.574 | -0.757 | 0.010 | 0.106 |
| ETV6 | 1E-04 | 0.378 | FALSE | 1 | 1 | -0.160 | 0.296 | 0.614 | -0.569 | 0.084 | 0.197 | -0.029 | 0.767 | 0.916 | -0.563 | 0.011 | 0.108 |
| GFI1 | 0.916 | -0.864 | FALSE | 1 | 1 | -0.123 | 0.288 | 0.614 | -0.665 | 0.011 | 0.076 | -0.172 | 0.036 | 0.534 | -0.714 | 0.002 | 0.069 |
| RUNX2 | 0.132 | -0.250 | FALSE | 1 | 1 | -0.151 | 0.312 | 0.614 | -0.810 | 0.027 | 0.114 | -0.393 | 0.005 | 0.492 | -0.887 | 6.4E-04 | 0.049 |
| PLAGL2 | 0.316 | -0.380 | FALSE | | 1 | -0.149 | 0.383 | 0.665 | -0.444 | 0.094 | 0.209 | -0.161 | 0.009 | 0.521 | -0.720 | 0.003 | 0.082 |
| SP1 | 0.864 | -0.764 | FALSE | | 1 | -0.156 | 0.395 | 0.672 | -0.653 | 0.038 | 0.130 | -0.127 | 0.164 | 0.544 | -0.624 | 0.024 | 0.123 |
| MEF2C | 0.523 | -0.496 | FALSE | 1 | 1 | -0.205 | 0.428 | 0.693 | -0.835 | 0.037 | 0.130 | -0.369 | 0.045 | 0.534 | -0.750 | 0.020 | 0.120 |
| CEBPA | 0.485 | -0.475 | FALSE | 1 | 1 | 0.127 | 0.484 | 0.718 | -0.293 | 0.137 | 0.257 | 0.103 | 0.377 | 0.687 | -0.389 | 0.073 | 0.167 |
| RXRA | 0.582 | -0.530 | FALSE | 1 | 1 | 0.099 | 0.555 | 0.761 | -0.371 | 0.248 | 0.366 | 0.104 | 0.249 | 0.597 | -0.300 | 0.225 | 0.344 |
| ZMYND8 | 0.920 | -0.875 | FALSE | 1 | | -0.059 | 0.738 | 0.884 | -0.629 | 0.041 | 0.134 | -0.058 | 0.754 | 0.915 | -0.786 | 7.6E-04 | 0.050 |
| MEF2D | 0.877 | -0.784 | FALSE | 1 | 1 | -0.018 | 0.913 | 0.962 | -0.632 | 0.003 | 0.046 | 0.021 | 0.823 | 0.939 | -0.444 | 0.060 | 0.153 |
| MAX | 0.395 | -0.425 | TRUE | | 1 | 0.179 | 0.460 | | -0.708 | 0.074 | 0.183 | -0.026 | 0.738 | 0.906 | -0.471 | 0.033 | 0.128 |
| FLI1 | 0.318 | -0.381 | FALSE | 1 | 1 | -0.167 | 0.537 | | -1.290 | 0.019 | 0.096 | -0.289 | 0.060 | 0.534 | -0.571 | 0.047 | 0.139 |
| SPI1 | 0.943 | -0.967 | FALSE | 1 | 1 | -0.182 | 0.603 | | 0.038 | 0.938 | 0.951 | 0.236 | 0.050 | 0.534 | -0.286 | 0.198 | 0.314 |
| STAT5B | 0.898 | -0.820 | FALSE | | 1 | 0.572 | 0.143 | | -0.287 | 0.661 | | 0.110 | 0.261 | 0.604 | -0.306 | 0.114 | 0.215 |
| SREBF1 | 0.898 | -0.820 | FALSE | 1 | 1 | 0.657 | 0.280 | | -0.133 | 0.813 | | 0.243 | 0.180 | 0.549 | -0.131 | 0.561 | 0.667 |
| TFAP4 | 0.941 | -0.959 | FALSE | | 1 | 0.089 | 0.833 | | -0.636 | 0.473 | | -0.064 | 0.587 | 0.824 | -0.624 | 0.018 | 0.116 |
| ZNF281 | 0.074 | -0.185 | FALSE | | 1 | -0.160 | 0.638 | | -0.746 | 0.099 | 0.214 | -0.557 | 0.146 | 0.538 | -1.042 | 0.021 | 0.121 |
| LYL1 | 0.142 | -0.260 | FALSE | 1 | 1 | 0.401 | 0.437 | | 0.018 | 0.966 | 0.974 | 0.243 | 0.081 | 0.534 | -0.197 | 0.425 | 0.543 |
| FOSL2 | 0.400 | -0.428 | FALSE | 1 | 1 | 0.019 | 0.944 | | -0.553 | 0.163 | 0.284 | -0.358 | 0.012 | 0.534 | -0.812 | 0.002 | 0.072 |
| GATA2 | 2E-04 | 0.296 | FALSE | 1 | 1 | 0.112 | 0.667 | | -1.029 | 0.047 | 0.141 | -0.080 | 0.342 | 0.660 | -0.627 | 0.024 | 0.123 |
| HHEX | 0.051 | -0.149 | FALSE | | 1 | -0.344 | 0.201 | | -1.312 | 0.032 | 0.121 | -0.495 | 0.007 | 0.521 | -0.871 | 7.8E-04 | 0.050 |
| ZFPM1 | 0.017 | -0.053 | FALSE | 1 | 1 | -1.127 | 0.253 | | -0.428 | 0.671 | | 0.519 | 0.030 | 0.534 | -0.345 | 0.262 | 0.383 |

### Figure S20

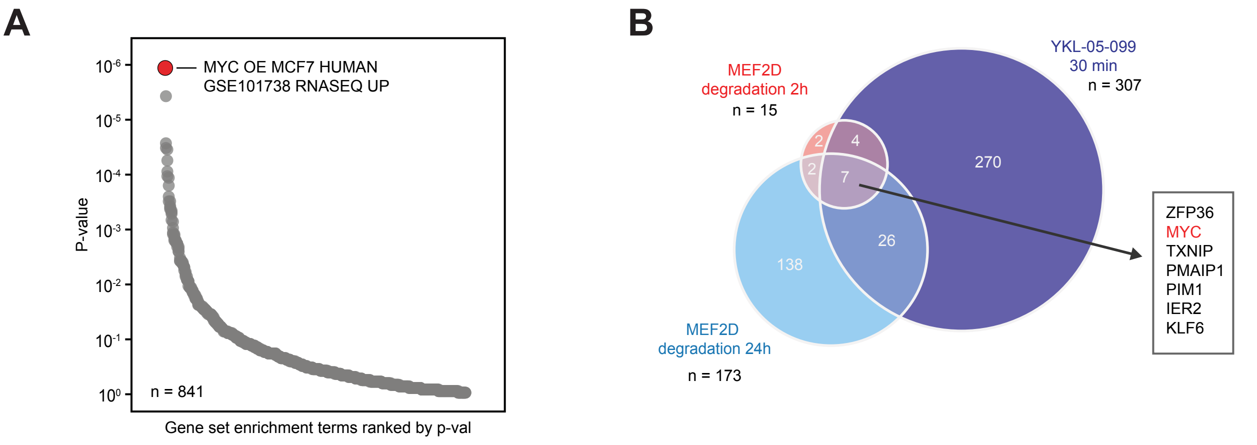


**C**

JQ1, μM

| YKL-05-009, nM | 12.5 | 6.25 | 3.125 | 1.563 | 0.781 | 0.391 |
|---|---|---|---|---|---|---|
| 400 | 0.05 | 0.091 | 0.022 | -0.05 | 0.05 | 0.031 |
| 200 | 0.08 | 0.136 | 0.018 | 0.058 | 0.064 | 0.06 |
| 100 | 0.049 | 0.48 | 0.059 | -0 | 0.14 | 0.076 |
| 50 | 0.029 | 0.024 | 0.043 | 0.015 | 0.064 | 0.06 |
| 25 | 0.057 | 0.045 | 0.039 | -0 | 0.117 | 0.136 |
| 12.5 | 0.049 | 0.068 | 0.07 | 0.011 | 0.069 | 0.072 |

**D**

MYCi361, μM

| YKL-05-009, nM | 3.125 | 1.563 | 0.781 | 0.391 | 0.195 | 0.098 |
|---|---|---|---|---|---|---|
| 400 | -0.05 | 0.03 | 0.089 | 0.102 | 0.133 | 0.032 |
| 200 | -0.04 | 0.095 | 0.091 | 0.145 | 0.095 | 0.034 |
| 100 | 0.033 | 0.016 | 0.037 | 0.079 | 0.15 | 0.072 |
| 50 | -0.04 | -0.02 | 0.252 | 0.063 | 0.042 | 0.05 |
| 25 | 0.04 | 0.039 | 0.093 | 0.06 | 0.059 | 0.038 |
| 12.5 | 0.034 | 0.005 | 0.029 | 0.023 | 0.044 | 0.052 |

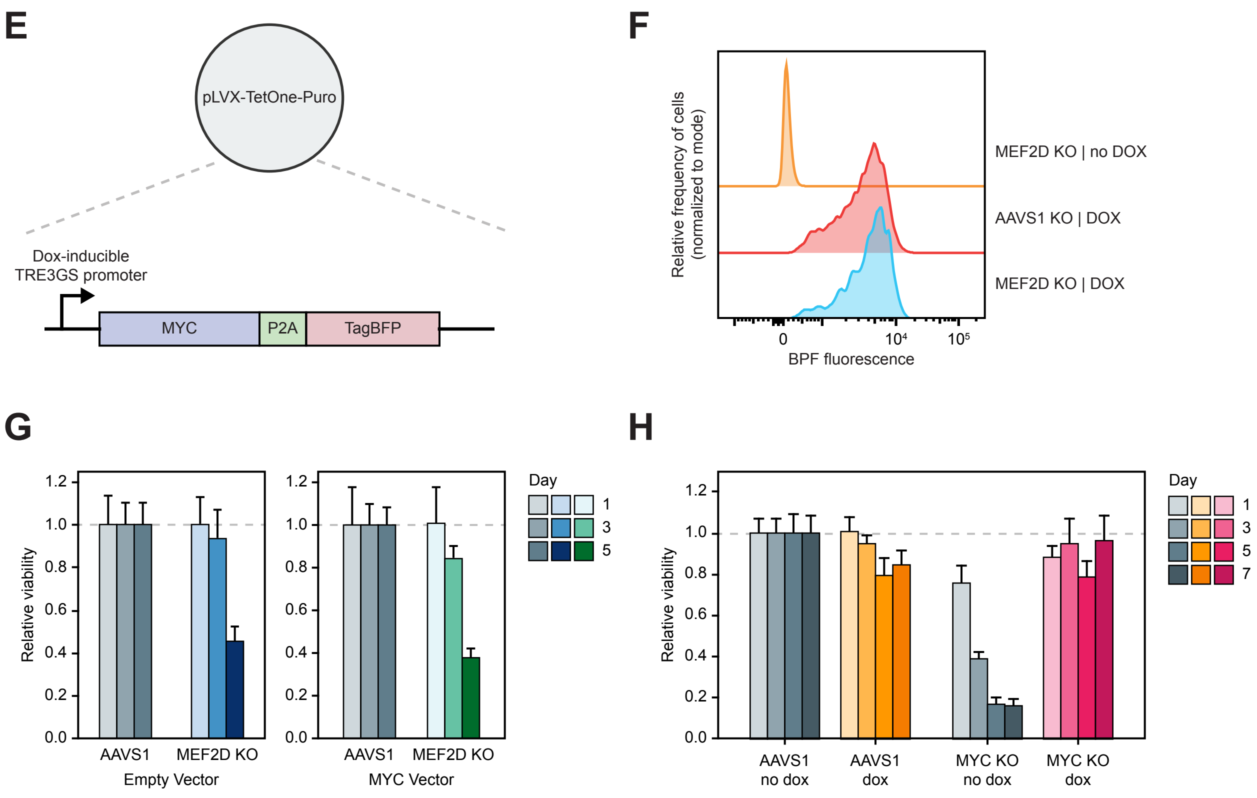

### Figure S21

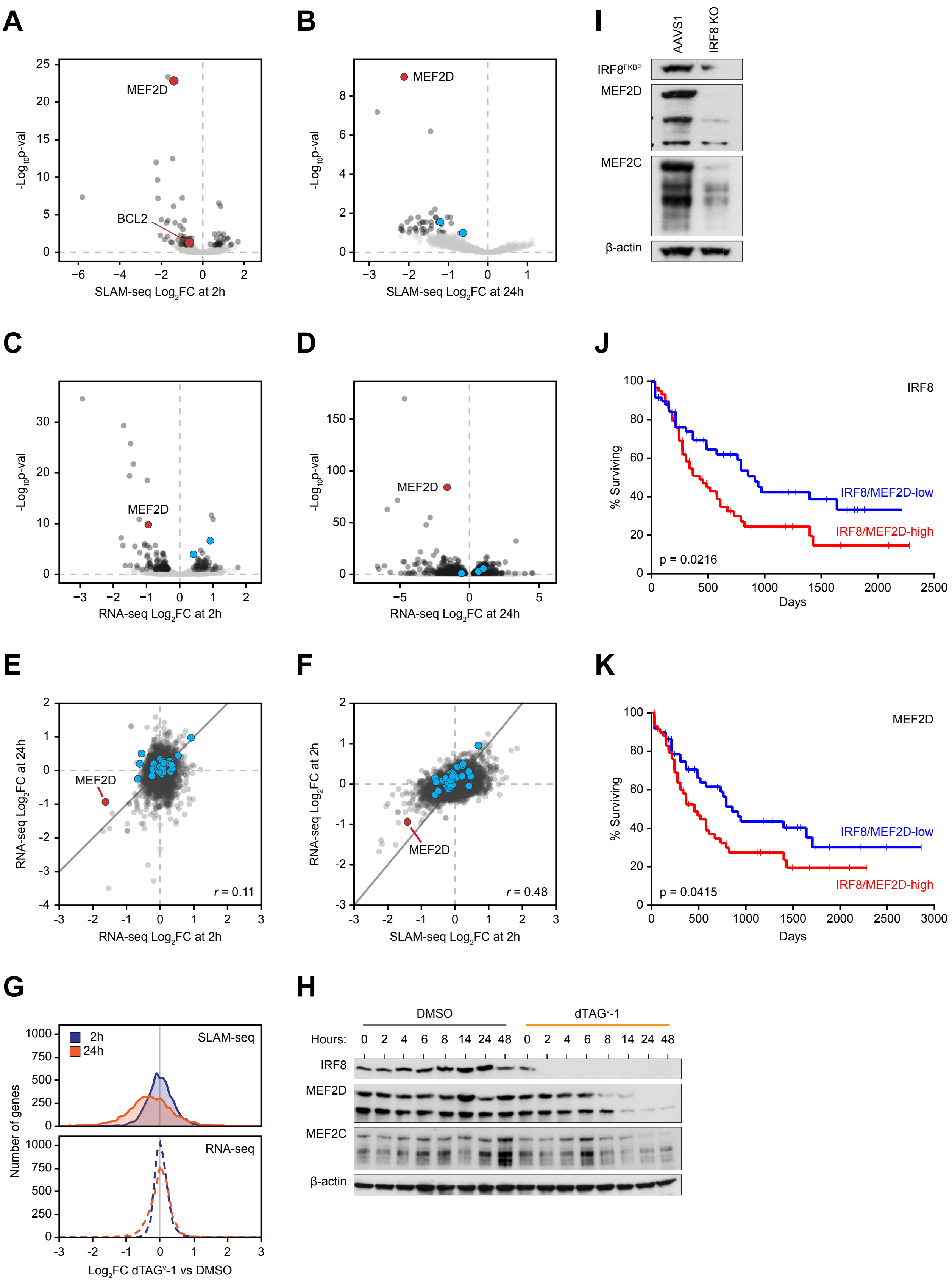

A
MEF2D
BCL2
-Log10p-val
SLAM-seq Log2FC at 2h
B
MEF2D
SLAM-seq Log2FC at 24h
I
AAVS1
IRF8 KO
IRF8FKBP
MEF2D
MEF2C
β-actin
C
MEF2D
RNA-seq Log2FC at 2h
D
MEF2D
RNA-seq Log2FC at 24h
J
IRF8
% Surviving
IRF8/MEF2D-low
IRF8/MEF2D-high
p = 0.0216
Days
E
RNA-seq Log2FC at 24h
MEF2D
r = 0.11
RNA-seq Log2FC at 2h
F
RNA-seq Log2FC at 2h
MEF2D
r = 0.48
SLAM-seq Log2FC at 2h
K
MEF2D
% Surviving
IRF8/MEF2D-low
IRF8/MEF2D-high
p = 0.0415
Days
G
2h
24h
SLAM-seq
RNA-seq
Number of genes
Log2FC dTAGv-1 vs DMSO
H
DMSO
dTAGv-1
Hours: 0 2 4 6 8 14 24 48 0 2 4 6 8 14 24 48
IRF8
MEF2D
MEF2C
β-actin

### Figure S22

| Gene ID | MV411 dep. probability | MV411 drop out score | 2 hrs SLAM-seq log$_2$FC | 2 hrs SLAM-seq pvalue | 2 hrs SLAM-seq padj | 24 hrs SLAM-seq log$_2$FC | 24 hrs SLAM-seq pvalue | 24 hrs SLAM-seq padj | 2 hrs RNA-seq log$_2$FC | 2 hrs RNA-seq pval | 2 hrs RNA-seq padj | 24 hrs RNA-seq log$_2$FC | 24 hrs RNA-seq pval | 24 hrs RNA-seq padj |
|---|---|---|---|---|---|---|---|---|---|---|---|---|---|---|
| CXorf21 | 0.027 | -0.089 | -1.672 | 9.9E-28 | 4.4E-24 | -2.795 | 6.0E-11 | 6.6E-08 | -0.971 | 1.4E-22 | 2.9E-19 | -1.824 | 1.8E-04 | 0.009 |
| MEF2D | 0.877 | -0.784 | -1.393 | 6.7E-27 | 1.5E-23 | -2.118 | 4.6E-13 | 1.0E-09 | -0.946 | 1.2E-13 | 1.5E-10 | -1.603 | 6.8E-89 | 5.0E-85 |
| FAM107B | 4E-04 | 0.237 | -1.453 | 2.2E-16 | 3.2E-13 | -1.241 | 0.002 | 0.113 | -0.690 | 8.4E-06 | 0.003 | -0.771 | 0.028 | 0.296 |
| CCR2 | 0.268 | -0.351 | -2.247 | 8.7E-16 | 9.6E-13 | -3.651 | 0.001 | | -1.684 | 8.2E-34 | 5.0E-30 | -1.842 | 2.0E-13 | 6.3E-11 |
| DCANP1 | 0.187 | -0.296 | -2.179 | 2.2E-13 | 1.9E-10 | -2.188 | 0.003 | | -1.394 | 6.2E-26 | 1.9E-22 | -1.295 | 2.7E-10 | 4.8E-08 |
| KCNA3 | 0.006 | 0.033 | -5.798 | 5.2E-11 | 3.8E-08 | -5.960 | 1.8E-06 | | -2.926 | 2.1E-39 | 2.6E-35 | -5.905 | 3.3E-67 | 1.2E-63 |
| CBX6 | 0.007 | 0.017 | -0.989 | 7.7E-11 | 4.9E-08 | -1.325 | 6.2E-05 | 0.015 | -0.378 | 0.002 | 0.145 | -0.971 | 0.005 | 0.111 |
| TIFAB | 0.007 | 0.016 | -2.156 | 1.0E-10 | 5.6E-08 | -4.186 | 0.003 | | -1.478 | 4.2E-30 | 1.7E-26 | -1.755 | 1.6E-19 | 1.6E-16 |
| LDLR | 0.579 | -0.528 | 0.770 | 5.0E-10 | 2.5E-07 | -0.116 | 0.617 | 0.859 | 0.957 | 1.2E-15 | 2.1E-12 | 0.132 | 0.351 | 0.800 |
| MALAT1 | | | 0.856 | 1.4E-09 | 6.0E-07 | -0.220 | 0.432 | 0.760 | 0.562 | 0.018 | 0.404 | -0.296 | 0.477 | 0.855 |
| CCR1 | 0.019 | -0.061 | -1.410 | 1.6E-09 | 6.3E-07 | -1.636 | 2.3E-04 | 0.028 | -0.902 | 7.6E-09 | 4.9E-06 | -1.001 | 0.001 | 0.042 |
| CPM | 0.007 | 0.020 | -1.991 | 1.0E-07 | 3.8E-05 | -1.815 | 1.6E-02 | | -0.494 | 1.3E-05 | 3.4E-03 | -1.606 | 7.2E-16 | 4.1E-13 |
| IKZF1 | 0.434 | -0.447 | -0.889 | 1.4E-07 | 4.8E-05 | -0.977 | 0.008 | 0.175 | -0.532 | 3.9E-04 | 0.045 | -0.301 | 0.261 | 0.735 |
| SERPINB8 | 0.006 | 0.029 | -1.678 | 1.6E-07 | 4.9E-05 | -1.501 | 0.045 | | -0.693 | 1.2E-04 | 0.019 | -1.033 | 0.026 | 0.286 |
| POU2F2 | 8E-05 | 0.393 | -1.745 | 3.6E-07 | 1.0E-04 | -2.154 | 0.001 | 0.076 | -1.510 | 1.6E-23 | 3.9E-20 | -3.523 | 5.7E-26 | 9.2E-23 |
| ABHD15 | 0.069 | -0.178 | -1.336 | 3.6E-07 | 1.0E-04 | -1.321 | 0.043 | | -0.494 | 2.4E-07 | 1.1E-04 | -0.628 | 2.5E-05 | 0.002 |
| IL6R | 0.021 | -0.067 | -1.098 | 9.6E-07 | 2.5E-04 | -0.493 | 0.352 | 0.717 | -0.517 | 1.4E-07 | 7.0E-05 | -0.457 | 0.056 | 0.407 |
| TMEM127 | 0.698 | -0.605 | -0.955 | 3.0E-06 | 7.4E-04 | -1.121 | 0.029 | 0.314 | -0.345 | 8.0E-04 | 0.077 | -0.140 | 0.360 | 0.804 |
| TRIB1 | 0.005 | 0.045 | 0.783 | 1.1E-05 | 0.003 | -0.084 | 0.703 | 0.897 | 0.687 | 1.1E-07 | 5.7E-05 | -0.075 | 0.750 | 0.947 |
| SCD | 0.919 | -0.871 | 0.566 | 1.4E-05 | 0.003 | -0.344 | 0.289 | 0.665 | 0.040 | 0.691 | 0.912 | 0.256 | 0.119 | 0.561 |
| HMGCS1 | 0.965 | -1.096 | 1.003 | 1.5E-05 | 0.003 | -0.891 | 0.033 | 0.335 | 0.784 | 1.1E-05 | 0.003 | -0.378 | 0.074 | 0.462 |
| OPN3 | 0.002 | 0.128 | -2.046 | 2.3E-05 | 0.004 | -2.730 | 0.034 | | -0.561 | 1.9E-04 | 0.027 | -0.488 | 0.103 | 0.531 |
| ADGRG5 | 0.016 | -0.044 | -1.060 | 2.3E-05 | 0.004 | -1.347 | 0.104 | | -0.579 | 2.6E-04 | 0.034 | -0.288 | 0.008 | 0.148 |
| CSF1R | 0.229 | -0.326 | -1.702 | 3.5E-05 | 0.006 | -1.519 | 0.109 | | -0.705 | 3.1E-07 | 1.3E-04 | -0.881 | 0.009 | 0.158 |
| MSMO1 | 0.002 | 0.110 | 0.887 | 4.0E-05 | 0.007 | -1.452 | 0.008 | 0.187 | 1.004 | 9.2E-15 | 1.3E-11 | 0.107 | 0.278 | 0.748 |
| TGFBR1 | 0.003 | 0.078 | -0.899 | 4.2E-05 | 0.007 | -1.727 | 3.1E-04 | 0.032 | -0.332 | 0.079 | 0.649 | -0.718 | 0.060 | 0.420 |
| P2RY2 | 0.006 | 0.025 | -0.538 | 6.6E-05 | 0.010 | -0.101 | 0.723 | 0.904 | -0.429 | 1.6E-04 | 0.024 | 0.068 | 0.643 | 0.916 |
| DDIT3 | 0.160 | -0.275 | 1.216 | 6.5E-05 | 0.010 | -0.146 | 0.678 | 0.884 | 0.330 | 0.028 | 0.476 | -0.153 | 0.452 | 0.845 |
| ZBTB33 | 0.032 | -0.103 | -0.944 | 7.6E-05 | 0.011 | -1.126 | 0.004 | 0.149 | -0.902 | 3.4E-05 | 0.008 | -1.022 | 0.005 | 0.112 |
| B3GNT7 | 0.004 | 0.056 | -0.684 | 7.6E-05 | 0.011 | -0.416 | 0.270 | 0.646 | -0.634 | 2.7E-06 | 9.7E-04 | -0.229 | 0.008 | 0.140 |
| CALR | 0.040 | -0.125 | 0.663 | 7.8E-05 | 0.011 | 0.583 | 0.096 | 0.477 | 0.093 | 0.447 | 0.820 | 0.623 | 2.6E-04 | 0.011 |
| DUSP7 | 0.004 | 0.056 | -0.633 | 8.4E-05 | 0.012 | -0.518 | 0.326 | 0.694 | -0.138 | 0.222 | 0.721 | -0.656 | 0.058 | 0.417 |
| INHBA | 0.012 | -0.023 | -0.939 | 9.7E-05 | 0.013 | -1.763 | 5.9E-05 | 0.015 | -0.277 | 0.083 | 0.650 | -1.321 | 2.1E-09 | 3.2E-07 |
| JUN | 0.374 | -0.413 | 1.026 | 1.1E-04 | 0.013 | 0.314 | 0.272 | 0.646 | 0.765 | 2.4E-05 | 0.005 | -0.152 | 0.584 | 0.895 |
| 284454 | | | 1.170 | 1.0E-04 | 0.013 | 0.681 | 0.023 | 0.298 | 0.639 | 0.042 | 0.548 | 0.039 | 0.871 | 0.979 |
| SH2B3 | 0.002 | 0.107 | -0.791 | 1.3E-04 | 0.016 | -1.384 | 0.124 | 0.503 | -0.461 | 3.6E-04 | 0.042 | -0.374 | 0.274 | 0.746 |
| LYZ | 0.041 | -0.126 | -1.829 | 1.4E-04 | 0.017 | -0.611 | 0.503 | | -0.275 | 0.044 | 0.556 | 0.546 | 0.135 | 0.588 |
| TIFA | 0.003 | 0.093 | -1.191 | 1.6E-04 | 0.019 | -1.692 | 0.012 | 0.210 | -0.653 | 3.6E-04 | 0.042 | -1.095 | 0.003 | 0.076 |
| ZNF616 | 4E-04 | 0.249 | -0.983 | 1.7E-04 | 0.019 | -0.824 | 0.450 | | -0.213 | 0.223 | 0.721 | 0.232 | 0.363 | 0.806 |
| RNF41 | 0.193 | -0.301 | -0.678 | 1.8E-04 | 0.020 | -0.898 | 0.050 | 0.368 | -0.183 | 0.074 | 0.644 | -0.223 | 0.028 | 0.293 |
| EGR1 | 0.007 | 0.017 | 0.837 | 2.9E-04 | 0.031 | -0.124 | 0.578 | 0.845 | 0.606 | 8.9E-05 | 0.017 | -0.342 | 0.240 | 0.715 |
| HELQ | 0.005 | 0.039 | -0.792 | 3.1E-04 | 0.032 | -0.613 | 0.173 | 0.564 | -0.351 | 0.012 | 0.344 | -0.180 | 0.562 | 0.887 |
| BIN3-IT1 | | | 1.691 | 3.2E-04 | 0.033 | 0.252 | 0.652 | 0.872 | 0.828 | 0.025 | 0.453 | -0.098 | 0.817 | 0.965 |
| MBP | 0.002 | 0.112 | -0.650 | 3.5E-04 | 0.035 | -0.755 | 0.010 | 0.205 | -0.385 | 0.015 | 0.383 | -0.184 | 0.377 | 0.813 |
| ZNF765 | 3E-04 | 0.288 | -1.544 | 3.6E-04 | 0.035 | -0.895 | 0.492 | | -0.265 | 0.250 | 0.736 | 0.067 | 0.889 | 0.982 |
| KLF2 | 0.002 | 0.121 | 0.989 | 3.7E-04 | 0.035 | 0.307 | 0.239 | 0.625 | 0.833 | 6.8E-05 | 0.014 | -0.585 | 0.023 | 0.266 |
| HES6 | 0.010 | -0.009 | 0.760 | 3.9E-04 | 0.037 | 0.523 | 0.199 | 0.580 | 0.281 | 0.187 | 0.703 | 0.065 | 0.845 | 0.972 |
| RHOU | 0.030 | -0.099 | -1.481 | 4.3E-04 | 0.039 | 0.229 | 0.771 | | -0.475 | 0.003 | 0.188 | 0.131 | 0.627 | 0.911 |
| TMED10 | 0.483 | -0.473 | -0.780 | 4.7E-04 | 0.042 | -0.838 | 0.120 | 0.497 | -0.254 | 0.093 | 0.653 | -0.215 | 0.439 | 0.839 |
| BCL2 | 0.576 | -0.526 | -0.649 | 5.4E-04 | 0.048 | -1.205 | 0.005 | 0.159 | -0.340 | 0.038 | 0.534 | -0.713 | 3.2E-07 | 3.4E-05 |
| FLVCR2 | 0.034 | -0.110 | -0.988 | 5.7E-04 | 0.048 | -1.404 | 0.012 | 0.211 | -0.418 | 1.2E-04 | 0.019 | -0.426 | 9.7E-05 | 0.005 |
| CRKL | 0.902 | -0.830 | -0.611 | 5.6E-04 | 0.048 | -0.039 | 0.967 | 0.986 | -0.155 | 0.240 | 0.732 | 0.192 | 0.614 | 0.906 |
| HNRNPA3 | 0.138 | -0.256 | 0.733 | 6.3E-04 | 0.053 | 0.223 | 0.478 | 0.791 | -0.119 | 0.464 | 0.825 | -0.133 | 0.604 | 0.902 |
| LRRC8C | 0.041 | -0.128 | -0.817 | 6.8E-04 | 0.055 | -2.041 | 1.8E-04 | 0.027 | -0.366 | 0.060 | 0.611 | -1.075 | 0.015 | 0.208 |

### Figure S22 cont.

| Gene ID | MV411 dep. probability | MV411 drop out score | 2 hrs SLAM-seq log$_2$FC | 2 hrs SLAM-seq pvalue | 2 hrs SLAM-seq padj | 24 hrs SLAM-seq log$_2$FC | 24 hrs SLAM-seq pvalue | 24 hrs SLAM-seq padj | 2 hrs RNA-seq log$_2$FC | 2 hrs RNA-seq pval | 2 hrs RNA-seq padj | 24 hrs RNA-seq log$_2$FC | 24 hrs RNA-seq pval | 24 hrs RNA-seq padj |
|---|---|---|---|---|---|---|---|---|---|---|---|---|---|---|
| LINC00599 | | | 0.849 | 7.3E-04 | 0.059 | 0.435 | 0.169 | 0.562 | 0.725 | 0.212 | 0.715 | 0.685 | 0.010 | 0.167 |
| NLRP3 | 0.002 | 0.130 | -1.080 | 8.1E-04 | 0.063 | -1.668 | 1.8E-04 | 0.027 | -0.688 | 7.2E-04 | 0.071 | -0.959 | 1.4E-08 | 2.0E-06 |
| ZNF189 | 5E-04 | 0.229 | -0.858 | 8.2E-04 | 0.063 | -1.239 | 0.036 | 0.347 | -0.349 | 0.056 | 0.603 | 0.110 | 0.787 | 0.960 |
| GLUL | 0.002 | 0.123 | 0.357 | 9.0E-04 | 0.066 | 0.521 | 0.058 | 0.383 | 0.141 | 0.191 | 0.708 | 1.087 | 1.1E-05 | 8.1E-04 |
| FADS1 | 0.006 | 0.034 | 0.710 | 9.1E-04 | 0.066 | -0.743 | 0.145 | 0.531 | -0.093 | 0.419 | 0.809 | 0.232 | 0.022 | 0.261 |
| MIR9-3HG | | | 0.780 | 9.1E-04 | 0.066 | -0.337 | 0.355 | 0.720 | 0.125 | 0.467 | 0.826 | 0.078 | 0.607 | 0.903 |
| GORASP1 | 0.091 | -0.207 | 1.392 | 8.9E-04 | 0.066 | 0.727 | 0.366 | | 0.303 | 0.040 | 0.542 | 0.998 | 2.1E-11 | 4.9E-09 |
| ZNF766 | 5E-04 | 0.215 | -0.591 | 9.6E-04 | 0.068 | -0.855 | 0.020 | 0.270 | 0.040 | 0.735 | 0.925 | 0.120 | 0.454 | 0.846 |
| TMEM70 | 0.087 | -0.203 | -0.996 | 0.001 | 0.072 | -1.093 | 0.029 | 0.312 | -0.213 | 0.248 | 0.735 | -0.138 | 0.544 | 0.880 |
| GNA13 | 0.003 | 0.073 | -0.838 | 0.001 | 0.072 | -1.282 | 0.011 | 0.208 | -0.540 | 0.008 | 0.291 | -0.629 | 0.242 | 0.717 |
| KCNQ1OT1 | | | 0.770 | 0.001 | 0.073 | 0.088 | 0.823 | 0.946 | 0.808 | 0.003 | 0.193 | -0.002 | 0.994 | 0.999 |
| RGS16 | 0.003 | 0.080 | 0.657 | 0.001 | 0.078 | -0.996 | 0.005 | 0.149 | 0.694 | 0.001 | 0.100 | -0.649 | 0.123 | 0.570 |
| INSIG1 | 0.005 | 0.046 | 0.584 | 0.001 | 0.079 | -0.431 | 0.227 | 0.609 | 0.705 | 0.000 | 0.004 | 0.100 | 0.727 | 0.940 |
| RCSD1 | 0.060 | -0.164 | -0.778 | 0.001 | 0.080 | -0.990 | 0.027 | 0.310 | -0.279 | 0.018 | 0.405 | -0.022 | 0.923 | 0.987 |
| HNRNPA2B1 | 0.314 | -0.379 | 0.585 | 0.001 | 0.086 | 0.488 | 0.189 | 0.571 | -0.034 | 0.775 | 0.938 | 0.109 | 0.420 | 0.832 |
| CXCL8 | 0.028 | -0.094 | 0.819 | 0.001 | 0.086 | -0.134 | 0.722 | 0.904 | 0.898 | 1.6E-05 | 0.004 | 0.062 | 0.902 | 0.984 |
| DMXL2 | 0.091 | -0.207 | 1.322 | 0.001 | 0.090 | 0.189 | 0.723 | 0.904 | -0.009 | 0.952 | 0.990 | 0.547 | 0.222 | 0.699 |
| BNIP2 | 0.060 | -0.164 | -0.792 | 0.002 | 0.093 | -0.811 | 0.025 | 0.298 | -0.523 | 0.015 | 0.388 | -0.325 | 0.350 | 0.799 |
| RAB11FIP1 | 3E-04 | 0.279 | -0.560 | 0.002 | 0.097 | -0.832 | 0.002 | 0.113 | -0.352 | 4.7E-04 | 0.052 | -0.210 | 0.221 | 0.697 |
